# A dynamic activation–memory cycle organizes the human IgG⁺ memory B cell compartment

**DOI:** 10.1101/2025.10.30.685644

**Authors:** Laura T.M. Graus, Andreas Chrysostomou, Denise Guerra, Gius Kerster, Karlijn van der Straten, Jacqueline van Rijswijk, Khadija Tejjani, Tim Beaumont, Maria Prins, Menno D. de Jong, Godelieve J. de Bree, Rogier W. Sanders, Marit J. van Gils, Mathieu Claireaux

## Abstract

B cells underpin durable immunity by generating long-lived memory B cells and antibody-secreting cells (ASCs) that re-engage upon antigen re-encounter. How human IgG⁺ memory B cells are organized to establish, maintain, and reactivate humoral immunity remains incompletely understood. Here, we combine multimodal single-cell profiling with longitudinal characterization of antigen-specific memory B cells following SARS-CoV-2 infection and vaccination to define the developmental relationships and biological functions of human IgG⁺ memory B cell subsets. We identify a coordinated activation–memory cycle shared by germinal center– and extrafollicular-derived memory B cells marked by differential CD45RB expression. Within this cycle, activated B cells represent specialized effector cells that acquire inflammatory responsiveness, migratory capacity, and differentiation potential toward ASCs. Activated memory B cells persist as an intermediate that retains migratory capacity while shifting toward homeostatic survival, preserving recall competence and contributing to the regeneration of long-lived resting memory. The resting memory compartment comprises two complementary populations: CD73⁺ resting memory B cells form the principal long-lived recall reservoir, whereas CD24⁺ resting memory B cells adopt a more regulatory resting state and shows limited participation in the SARS-CoV-2 recall response. We further identify a CD24⁺ intermediate population as the earliest transitional state emerging upon memory B cell reactivation, bridging resting memory and the effector recall response. Together, these findings establish the human IgG⁺ memory B cell compartment as a dynamic activation–memory cycle rather than a collection of static subsets, providing a framework for understanding humoral immunity and interpreting B cell responses in vaccination, infection, and immune-mediated disease.

**One sentence summary:** Our results provide novel insights on B cell recall responses after SARS-CoV-2 infection or vaccination, formulating distinct classical and non-classical re-activation trajectories from a resting memory state through intermediate phenotypes towards an activated state.

## Introduction

B cells play a crucial role in establishing long-lasting targeted immune protection against foreign pathogens through the generation of antibody-secreting cells (ASCs) and memory B cells (MBCs). Upon pathogen encounter or vaccination, pathogen-derived antigens are recognized by the B cell receptor (BCR), triggering B cell activation and differentiation within secondary lymphoid organs. Early activated B cells can follow two main pathways. Some differentiate through the extrafollicular route, producing short-lived ASCs and early MBCs for rapid and broad responses. Others enter the germinal centers (GCs) within secondary lymph nodes, where they undergo affinity maturation to improve BCR affinity^1–8^. These GC processes drive the selection of B cell clones with the highest affinity for the antigen, while B cells with insufficient affinity are eliminated. Selected B cells persist and either participate in further rounds of proliferation and antigen specificity refinement within the GC, or exit as precursors of longlived ASCs and MBCs. Long-lived ASC precursors migrate to the bone marrow where they mature into plasma cells and sustain long-term antibody production. MBCs recirculate into the periphery and respond rapidly upon antigen re-exposure by differentiating into ASCs or reentering GCs for additional maturation^5,9–12^.

While this framework defines the principal routes through which B cells generate ASCs and MBCs, how the resulting human memory compartment is organized into functionally distinct subsets, how these subsets are mobilized upon antigen re-encounter, and how they subsequently return to quiescence remain incompletely understood.

Studies of peripheral blood have nonetheless provided an understanding of B cell heterogeneity, with circulating CD19^⁺^ B cells traditionally classified based on IgD and CD27 surface proteins expression into four subsets: naive B cells (CD19^⁺^ IgD^⁺^ CD27^-^), unswitched MBCs (CD19^⁺^ IgD^⁺^ CD27^⁺^), classical/switched MBCs (CD19^⁺^ IgD^−^ CD27^⁺^) and double negative (DN) MBCs (CD19^⁺^ IgD^−^ CD27^−^)^13–15^. DN B cells can be further classified into different subsets, including DN1 (IgD^-^ CD27^-^ CD21^⁺^ CD11c^-^ T-bet^-^), DN2 (IgD^-^ CD27^-^ CD21^-^ CD11c^⁺^ T-bet^⁺^), and DN3 (IgD^-^CD27^-^CD21^-^ CD11c^-^T-bet^low^) B cells. Together with the activated naive subsets (CD19^⁺^ IgD^⁺^ CD27^-^ CD11c^⁺^ T-bet^⁺^), DN2 and DN3 define the atypical B cell (AtypB) compartment that is expanded in the elderly and individuals with autoimmune and/or chronic disease^16–20^. AtypB also contribute to early B cell responses following viral infections, including SARS-CoV-2, where they correlate with disease severity and are thought to arise through the extrafollicular pathway^21–24^. DN1 B cells are considered to be a durable product of the extrafollicular response. This subset is characterized by low somatic hypermutation (SHM) levels and increased clonal expansion compared to GC-dependent responses^25^. DN1 B cells are often associated with autoimmune diseases, but are also linked to the preservation of immunological breadth providing an advantage in vaccine responses^25,26^. In contrast, the classical MBC compartment mostly represents the GC-derived responses and can be further subdivided using CD21 and CD71 distinguishing CD27⁺CD21⁻CD71^⁺^ activated B cells (ActB) from CD27⁺CD21⁺ MBCs. GC-derived ActB peak shortly after infection or vaccination, but may emigrate from the GC up to 6 months after infection^27,28^. Eventually, these B cells contract over time, while MBCs increase in frequency. Importantly, ActB are interpreted as recent GC emigrates and potential precursors of ASCs^29,30^. More recent broader phenotyping using additional markers such as CD45RB, CD39, CD73, and CD95 has revealed a more heterogeneous MBC population with variable activation and differentiation states^31,32^. CD45RB was notably enriched within classical MBCs and has been associated with memory populations of GC origin^31,32^. Moreover, our recent work^33^ demonstrates substantial diversity especially within the CD27⁺CD21⁺ classical MBC compartment. This population comprises at least two major subsets: (i) an Activated Memory B cell (ActMem) fraction (CD73⁻CD24⁻), predominantly composed of B cells that respond to recent antigen exposure, and (ii) a resting MBC fraction (CD73⁺ and/or CD24⁺), enriched for B cells recognizing antigens encountered in the past.

Further stratification by CD95 expression reveals additional diversity within the resting pool, including a CD95⁻CD73⁺CD24^lo^ subset consistent with long-lived memory.

Despite these advances, the developmental position, re-activation potential, and differentiation trajectories of these subsets toward ASCs and MBCs remain poorly defined. Resolving these trajectories should reveal the cellular basis of durable humoral immunity, inform rational vaccine design, and provide a reference map to understand why these processes become dysregulated in autoimmune disease and allergy.

To address this, we set out to track spike-specific B cells using SARS-CoV-2 breakthrough infection and vaccination as models of antigen-driven recall. We performed multi-omics singlecell sequencing, integrating transcriptome, surface proteomics, antigen specificity, and BCR repertoire information, and verified our results with independent complementary longitudinal validation by spectral flow cytometry. Our data provides novel insights of extrafollicular and GC-dependent activation pathways and shows the parallel occurrence of both pathways within the activated and resting MBC compartments, distinguishable by CD45RB expression^32^. Furthermore, we defined a B cell subset functioning as a gateway for B cell re-activation. The phenotypic and functional profile of B cell subsets within the MBC subsets were further validated by longitudinal analysis providing additional evidence of these distinct B cell re-activation trajectories. Together, these integrated analyses offer additional insights into the complexity and interconnection between different MBC subsets that contribute to protective immunity following infection, providing novel leads for vaccine design and implications for optimal timing of vaccine regimens.

## Results

### A multi-omics approach allows for simultaneous analysis of B cell transcriptome, phenotype and BCR repertoire>

To investigate the diversity and developmental trajectories of recall B cell responses, we used SARS-CoV-2 breakthrough infection as a model of antigen-driven recall immunity. B cells were analyzed from 12 individuals in the COSCA cohort^34,35^ who had received at least two doses of a COVID-19 mRNA vaccine and subsequently experienced breakthrough infection with Delta or Omicron (BA.1, BA.2, BA.4, or BA.5). Samples were collected 4–7 weeks post-infection, a window that captures peak recall responses including GC-derived outputs and early memory B cell re-activation.

SARS-CoV-2-specific B cells were isolated by FACS alongside antigen-unspecific B cells to enrich for antigen-driven responses while retaining immunological context. These cells were then jointly profiled using a multi-omics single-cell approach on the 10x Genomics Chromium platform, integrating gene expression, BCR sequencing, and Cellular Indexing of Transcriptomes and Epitopes (CITE-seq) to simultaneously resolve transcriptional state, cell surface phenotype, and clonal repertoire within the same cells. The CITE-seq library included barcoded hashtag antibodies for sample multiplexing and donor identification, a 140-antibody panel for B cell surface marker phenotyping, and barcoded SARS-CoV-2 spike and receptorbinding domain (RBD) proteins to assess B cell specificity toward the ancestral (WT) virus and breakthrough variants (Figure 1). To enable sensitive detection of antigen-specific B cells across a range of affinities, SARS-CoV-2 proteins were conjugated to barcoded dextramer-based probes, which preserve native antigen conformation and are compatible with 10x Genomics single-cell multi-omics and flow cytometry workflows. This approach allowed for reliable identification of both high- and low-affinity antigen-specific B cells by flow cytometry and single-cell sequencing (See supplementary methods and supplementary figures S1 and S2 for comparison of different probe strategies).

**Figure 1.**
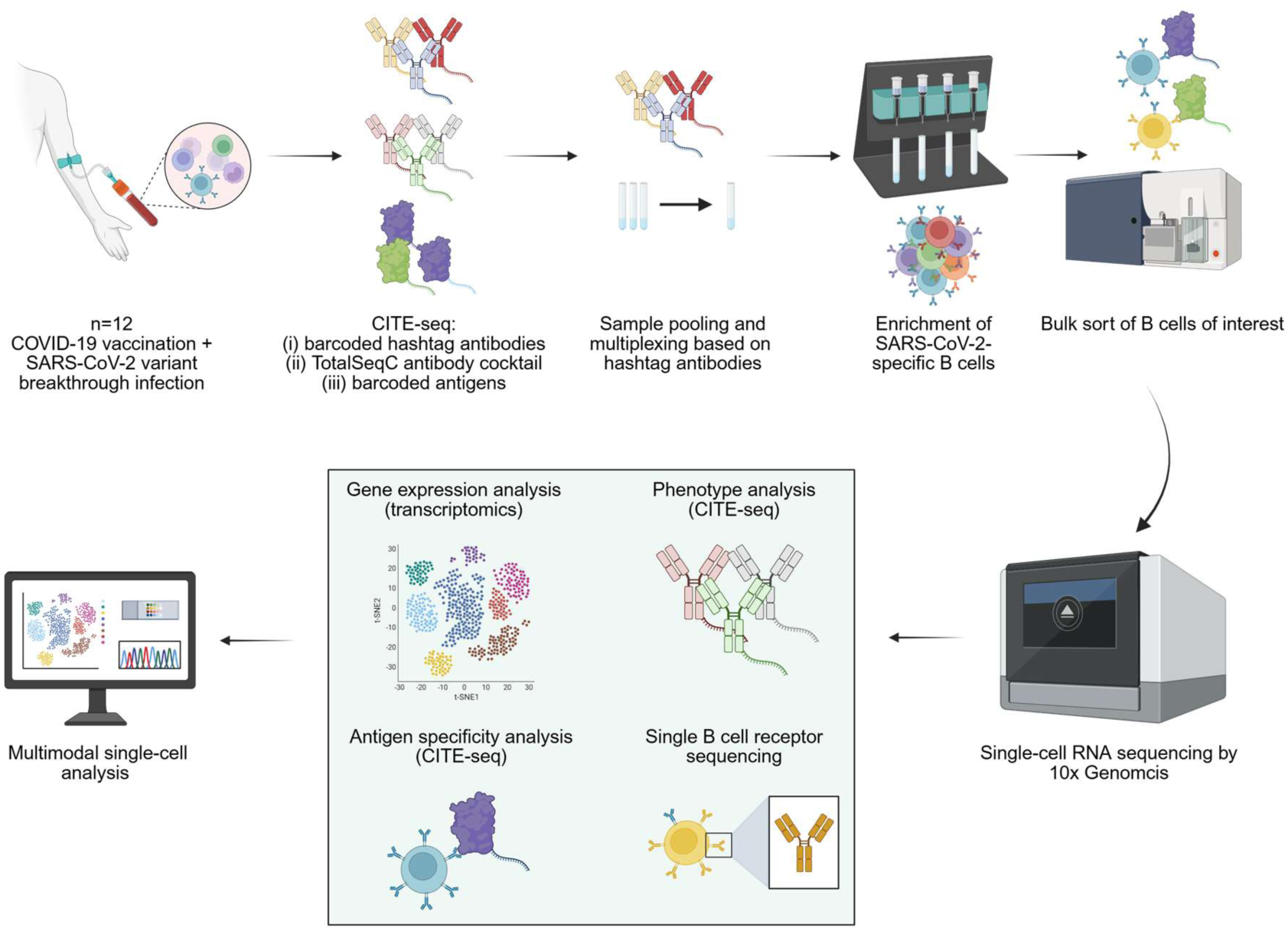
Multi-omics approach for integrated single B cell transcriptome, phenotype, antigen-specificity, and BCR repertoire analysis. Schematic representation of our experimental pipeline to detect, sort and sequence B cells by 10x Genomics. Three parallel single B cell libraries were obtained, corresponding to transcriptomics, proteomics, antigen specificity, and BCR repertoire data. Created in BioRender. Van Gils, M. (2026) https://BioRender.com/rlf2299.

### Multimodal integration identifies functionally distinct memory B cell subpopulations and reveals their underlying relationships

Transcriptomic and cell-surface protein measurements were integrated using a Weighted Nearest Neighbor (WNN) analysis^36^ to define B cell clusters. This approach resolved B cells into two major compartments: a memory/atypical B cell-enriched compartment and a naive/transitional B cell compartment (Figure 2A and Figure S3A). B cells from participant COSCA-365 formed a distinct cluster driven by donor-specific transcriptional effects, and were excluded from further analyses. Other participants were evenly distributed within all B cell subsets (Figure S3B). Subsequent analyses focused on IgG⁺ and atypical B cells (366 antigen-specific and 764 antigen-unspecific cells), excluding naive and IgM⁺ MBCs, as class-switched populations represented the dominant recall compartment.

**Figure 2.**
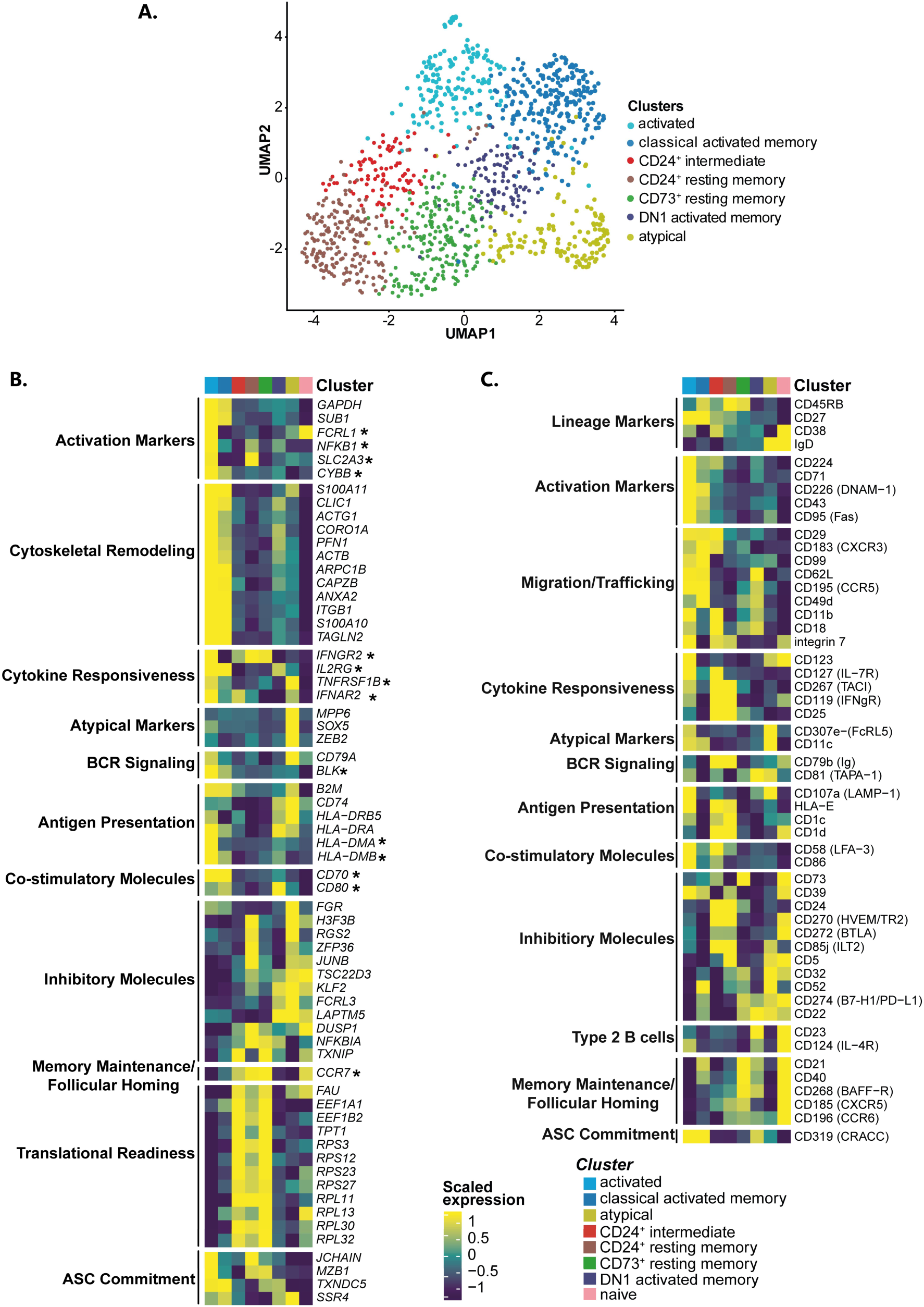
Characterization of distinct B cell populations through multimodal data analysis. **(A)** UMAP plot derived from WNN multimodal analysis, showing the IgG⁺ MBC compartment. The clustering was performed on the UMAP embeddings, using the unsupervised Leiden algorithm. The annotation was conducted based on known cell surface protein markers. **(B)** mRNA markers per MBC cluster as defined by unsupervised and supervised analysis. The markers were identified through spatial autocorrelation analysis using Moran’s I test. The markers are ordered based on their functionality. Supervised-selected genes are highlighted by an asterisk. Here, a subset of mRNA markers is shown based on biological relevance, while the full panels are reported in Supplementary Figure S4B-C. **(C)** Exploratory surface proteins per MBC cluster. The markers are significantly differentially expressed, with the significance being tested by the MAST test. The markers are ordered based on their functionality. Here, a subset of the entirety of the antibody-derived tags (ADT) markers is shown based on biological relevance, while the full ADT panel with biologically relevant surface proteins are reported in Supplementary Figure S4A.

WNN analysis resolved seven major IgG⁺ MBC subpopulations (Figure 2A) annotated using integrated B cell surface-marker patterns and previously established reference frameworks^33,37,38^: ActB, ActMem, comprising classical and DN1 fractions, RestMem consisting of CD73⁺ RestMem and CD24⁺ RestMem, AtypB, and an intermediate population of B cells bridging the resting and activated compartments, termed CD24⁺ intermediate B cells (CD24⁺ Int). Surface protein expression, transcriptomic signatures, and pathway activity were integrated to characterize each population and study their interrelationships (Figure 2C–D, Supplementary Figure S4, Supplementary Tables S3-S6). Throughout, italicized names refer to gene expression derived from transcriptomics; non-italicized names refer to surface protein expression measured by CITE-seq.

### Activated B cells are considered an effector B cell population characterized by enhanced migratory and activation-related capabilities

The ActB subset displayed a classical activated B cell phenotype characterized by high expression of CD27, CD43, and CD71 and low levels of CD21^29,30,33,37,38^. Furthermore, ActB showed reduced levels of *CCR7*, CXCR5, BAFF-R, and CD40, consistent with loss of the resting memory profile associated with lymphoid homing, follicular localization, and B cell maintenance. Concurrently, ActB upregulated a migration-associated module including CD62L, CD29, CD49d, integrin β7, CD11b, and CXCR3, reflecting enhanced migratory capacity toward both secondary lymphoid organs and peripheral tissue during inflammation. ActB displayed the strongest activation signature of all populations, marked by high expression of activation markers (CD71, CD43, CD95, CD224, CD226, and *FCRL1)*. In parallel, ActB combined expression of key BCR signaling components (e.g., *CD79a*, CD79B, CD81, and *BLK*), with increased expression actin-associated and cytoskeletal remodeling genes involved in BCR signaling and cellular migration (e.g., PFN1, ARPC1B, CORO1A, CAPZB, TAGLN2, TMSB4X, ACTB, and ACTG1). ActB also expressed elevated levels of molecules associated with inflammatory survival and cytokine responsiveness (e.g., CD267, *TNFRSF1B*, *IFNGR2*, CD123, and *IL2RG*) reflecting a switch from homeostatic to activation-driven survival signals.. Also, ActB exhibited expression of antigen-processing and presentation machinery, including LAMP1, HLA molecules, *B2M* and CD74^39–41^, together with T cell co-stimulatory molecules CD58, *CD70*, *CD80* and *CD86*^24,42–47^. Finally, a subset of ActB expressed the plasmablast-associated markers CD319, JCHAIN and MZB. (Figure 2C–D, Supplementary Figure S4, Supplementary Tables S3-S6).

Together, these features define the ActB subset as a major activated effector B cell population characterized by enhanced migratory, antigen-presenting, and T cell co-stimulatory capacities, accompanied by reduced follicular homing and memory-associated features. In agreement with previous studies, the expression of plasmablast-associated markers in a subset of ActB is consistent with ASC differentiation^48–50^. This positions the ActB subset as recently activated B cells that arise upon antigen encounter.

### Atypical B cells are a heterogeneous subset of activated B cells with a distinct expression profile

AtypB represent a phenotypically heterogeneous compartment previously associated with extrafollicular activation, chronic immune stimulation, and inflammatory disease^51,52^. Visual inspection of marker expression within the AtypB compartment identified three subclusters: activated CD73⁺, activated naive and DN2 B cells. Among these, only activated naive (IgD⁺) and DN2 (IgD^-^) cells displayed the canonical atypical B cell phenotype, characterized by loss of CD27, CD45RB, CD21, together with expression of CD11c, FCRL5, *SOX5* and *ZEB2*^18^. Accordingly, the AtypB-associated transcriptional and phenotypic signatures described below refer predominantly to activated naive and DN2 populations (Supplementary Figure S5-6). Similar to ActB, AtypB displayed reduced expression of *CCR7*, CXCR5, BAFF-R, and CD40, consistent with loss of the memory profile associated with follicular homing and B cell maintenance. In contrast to ActB however, AtypB lacked the migration- and adhesion-associated module linked to lymphoid recirculation and peripheral tissue trafficking. AtypB exhibited a partially activated phenotype characterized by intermediate expression of selected activation markers (e.g. CD71, CD43), but lacked the broad activation and cytokine-responsiveness gene profile observed in ActB. Consistent with this intermediate activation state, AtypB also expressed lower levels of actin-associated and cytoskeletal remodeling genes than ActB. In contrast, AtypB displayed a prominent inhibitory profile encompassing multiple inhibitory receptors (e.g. *LAPTM5,* CD22, *CD72,* CD32, FCRL5, *FCRL3,* CD85j), transcriptional regulators (e.g. *JUND, TSC22D3*), and post-transcriptional and chromatin-associated regulators (ZFP36, H3F3B). Pathway analysis further supported this restrained activation state, revealing a trend towards enrichment of EGFR, JAK-STAT, PI3K and TNFa signaling, albeit at lower levels than in ActB. Despite this inhibitory phenotype, AtypB displayed a prominent antigen-processing and presentation profile, characterized by expression of molecules involved in antigen processing, MHC class II loading, and antigen presentation (e.g., CD107a, CD74, *B2M, HLA-DRA, HLA-DR, and HLA-DRB5*). However, unlike ActB, AtypB expressed only low levels of the T cell co-stimulatory molecules CD80, CD86, CD58, and CD40, suggesting a reduced capacity to fully activate T cells and receive T cell help (Figure 2C–D, Supplementary Figure S4, Supplementary Tables S3-S6).

Together, these findings are consistent with AtypB representing a restrained effector state, distinguished from the highly activated ActB population by an extensive inhibitory profile despite retaining antigen-presenting capacity — in line with previous descriptions of atypical B cells in chronic immune stimulation^53^.

### Resting memory is defined as a quiescent state comprised of phenotypically and functionally distinct populations

In contrast to both the ActB and AtypB populations, the RestMem compartment was defined by expression of CD21 and absence of CD71, with differential expression of CD73 and CD24 distinguishing two subsets: CD73⁺ RestMem and CD24⁺ RestMem. Both populations expressed higher levels of CXCR5, CCR6, CCR7, BAFF-R, and CD40, consistent with preservation of a resting memory profile supporting follicular recirculation, B cell maintenance, and T cell interactions. In addition, both subsets exhibited a prominent translation-associated profile (e.g., EEF1, FAU, TPT1, and ribosomal RPL/RPS genes), indicative of a preserved biosynthetic readiness despite their quiescent state. Indeed, both RestMem subsets shared a core quiescent profile characterized by low expression of activation markers (e.g., CD71, CD43, CD224, and CD226), T cell co-stimulatory molecules (CD80 and CD86), BCR signaling components (e.g., CD79a, CD79b, and CD81), classical antigen-presentation molecules (e.g., HLA-DR, HLA-DM, and B2M), actin-associated and cytoskeletal remodeling genes, and trafficking-associated receptors (e.g., CD62L, CD29, and CXCR3) compared with ActB (Figure 2C–D, Supplementary Figure S4, Supplementary Tables S3-S6).

Despite this common resting phenotype, CD73⁺ and CD24⁺ RestMem displayed distinct profiles. CD73⁺ RestMem preferentially expressed trafficking molecules CCR1, CCR5, CD49d, CD11b, and CD18, together with higher levels of BAFF-R and CD40, consistent with a homeostatic memory population poised for long term maintenance and reactivation. In contrast, CD24⁺ RestMem preferentially expressed integrin β7 together with the cytokine-responsive molecules TACI, IFNGR, and CD25, consistent with reduced homing capacity and an innate-like or regulatory B cell phenotype. CD24⁺ RestMem also displayed enhanced expression of the non-classical antigen-presentation molecules CD1c, CD1d, and HLA-E, further distinguishing this subset from the other B cell populations described thus far.

Analysis of regulatory molecules further highlighted this heterogeneity. Both populations expressed CD32, CD52, DUSP1, and NFKBIA. However, CD73⁺ RestMem preferentially expressed CD22, PD-L1, and TXNIP, whereas CD24⁺ RestMem expressed inhibitory receptors BTLA, HVEM, CD39, CD85j, CD5, together with regulatory molecules JUNB, JUND, ZFP36, RGS2, and H3F3B, several of which were shared with AtypB. Consistent with these phenotypic differences, pathway analysis revealed broadly reduced activation and growth signaling across both subsets and increased TGFβ signaling within the CD24⁺ RestMem (Figure 2C– D, Supplementary Figure S4, Supplementary Tables S3-S6).

Together, these findings define CD73⁺ and CD24⁺ RestMem as functionally distinct resting programs within a shared quiescent state. CD73⁺ RestMem are characterized by a homeostatic maintenance, whereas CD24⁺ RestMem displayed a tightly regulated, environmentally responsive, immunoregulatory profile, highlighting divergent roles within the quiescent MBC compartment.

### Activated Memory B cells represent a transitional state linking the activated and resting memory compartments

The ActMem compartment comprising classical and DN1 subsets was defined by intermediate CD71 expression while retaining CD21 expression. Consistent with their phenotypic distinction, the classical ActMem subset was enriched for CD45RB⁺CD27⁺ cells, whereas DN1 Act-Mem was enriched for CD45RB⁻CD27⁻ cells. Both populations displayed a transitional phenotype between ActB and RestMem cells, characterized by intermediate expression of activation molecules (e.g., CD71, CD43, *FCRL1, NFKB1)*, while maintaining migration- and cytoskeletal remodeling-associated functionality. Compared with ActB, both ActMem subsets showed reduced inflammatory cytokine responsiveness, antigen-presentation machinery (CD74, HLA-DR), and T cell co-stimulatory molecules (CD80, CD70). Concurrently, they acquired multiple features associated with RestMem, including increased expression of CD21, CD40 and BAFFR together with inhibitory molecules (e.g. CD22, PDL1, NFKBIA, TXNIP), and translation-associated genes (RPS/RPL genes).

Despite these shared characteristics, classical ActMem retained higher levels of most activation molecules and cytoskeletal remodeling genes than DN1 ActMem. The two populations also displayed distinct trafficking-associated (classical ActMem: CD29, CXCR3, CD99; DN1 ActMem: CD11b, CD18), and regulatory profiles (classical ActMem: CD32, CD52; DN1 Act-Mem: *JUND, TSC22D3, KLF2, FCRL3*). These differences indicate distinct migration and regulatory profiles within the ActMem compartment. Notably, the DN1 ActMem subset was highly enriched for cells expressing CD23 and IL4R, markers previously associated with Type 2 B cells, involved in allergy as reservoir of IgE memory^54^, further supporting the distinct functional identity of this population. Pathway analysis mirrored the phenotypic findings, with both ActMem populations displaying intermediate pathway activity between ActB and RestMem across the majority of signaling pathways (Figure 2C–D, Supplementary Figure S4, Supplementary Tables S3-S6).

Together, these findings establish classical ActMem and DN1 ActMem as two distinct transitional MBC populations linking activated and resting memory compartments. While both populations acquire features associated with memory reformation, their divergent activation, trafficking, and regulatory profiles support distinct activation states and potential differentiation pathways within the human MBC populations.

### CD24**⁺** intermediate is a phenotypically distinct transitional population functioning as cellular gateway towards B cell activation

CD24⁺ Int represent an additional transitional phenotype between RestMem and ActB as apparent by their intermediate expression of activated as well as resting-associated markers. This subset was defined by expression of CD24 and CD1d, intermediate expression of CD71, and absence of CD21. CD24^+^ Int have previously been described as a phenotypically distinct activated subset. However, these B cells have been annotated as CD24^+^ Int in our data as we do not consider these cells as fully activated^37^. CD24⁺ Int were characterized by intermediate expression of activation-associated molecules (e.g. CD224, CD226, and *NFKB1)* highlighting their transitional state. CD24⁺ Int acquired several migration-associated features shared with ActB, including CD29, CD11b, CD18, and Integrin 7, although they lacked CD62L expression. In contrast to ActB, however, they expressed low levels of cytoskeletal remodeling genes.

They also retained broad cytokine responsiveness (e.g., IL7R, TACI, IFNgR, CD25, *IFNAR*), a profile shared with both ActB and CD24⁺ RestMem. Similar to CD24⁺ RestMem, CD24⁺ Int preferentially expressed high levels of non-classical presentation molecules (CD1c, CD1d, HLA-E), while displaying reduced expression of classical antigen-presentation machinery. Likewise, T cell co-stimulatory molecules exhibited an intermediate profile, with higher expression of CD58 and CD86 than RestMem but lower CD70 expression than ActB. CD24⁺ Int cells also retained elements of the regulatory profile observed in the RestMem compartment, including expression of HVEM, BTLA, CD85j, and TNXIP, together with a translation-associated RPL/RPS resembling that of RestMem cells. Consistent with their intermediate positioning, the CD24⁺ Int displayed a partial memory profile, lacking CD21 and CCR6, while expressing intermediate levels of CD40, BAFFR and CXCR5, together with high *CCR7* gene expression. Pathway analysis further placed CD24⁺ Int closer to RestMem than ActB (Figure 2C–D, Supplementary Figure S4, Supplementary Tables S3-S6).

Overall, these findings position CD24⁺ Int as a transitional population between resting memory and activated B cell states similar to the ActMem compartment, while showing a distinct antigen-presentation profile. While maintaining homeostatic, and regulatory, biosynthetic features, they simultaneously acquired activation, migration and cytokine-responsive profiles characteristic of ActB. Potentially, this subset functions as a gateway towards B cell re-activation.

In conclusion, our analyses resolved the IgG⁺ memory B cell compartment into eight phenotypically and transcriptionally distinct states: a fully activated compartment (ActB), two transitional populations bridging activation and resting memory (classical ActMem and DN1 Act-Mem), two resting memory subsets with divergent regulatory profiles (CD73⁺ and CD24⁺ Rest-Mem), a re-activation intermediate (CD24⁺ Int), and an AtypB compartment (activated naive and DN2). The progressive acquisition and loss of activation, trafficking, antigen-presentation, and regulatory profiles across these populations suggested an ordered differentiation continuum underlying human memory B cell responses.

### Multilayer analysis supports an activation-to-resting memory trajectory and resolves parallel germinal center and extrafollicular contributions

To resolve developmental relationships between IgG⁺ B cell populations, we performed gene expression–based lineage inference and integrated independent evidence from BCR SHM, antigen specificity, and monoclonal antibody (mAb) profiling. Lineage inference positioned ActB at the origin of the principal trajectory, progressing through ActMem compartments toward CD73⁺ and CD24⁺ RestMem as the quiescent endpoint, while identifying CD24⁺ Int as a re-entry node linking RestMem to ActB states (Figure 3A). In contrast, AtypB fell largely outside the main trajectory, consistent with a distinct differentiation pathway^25^.

**Figure 3.**
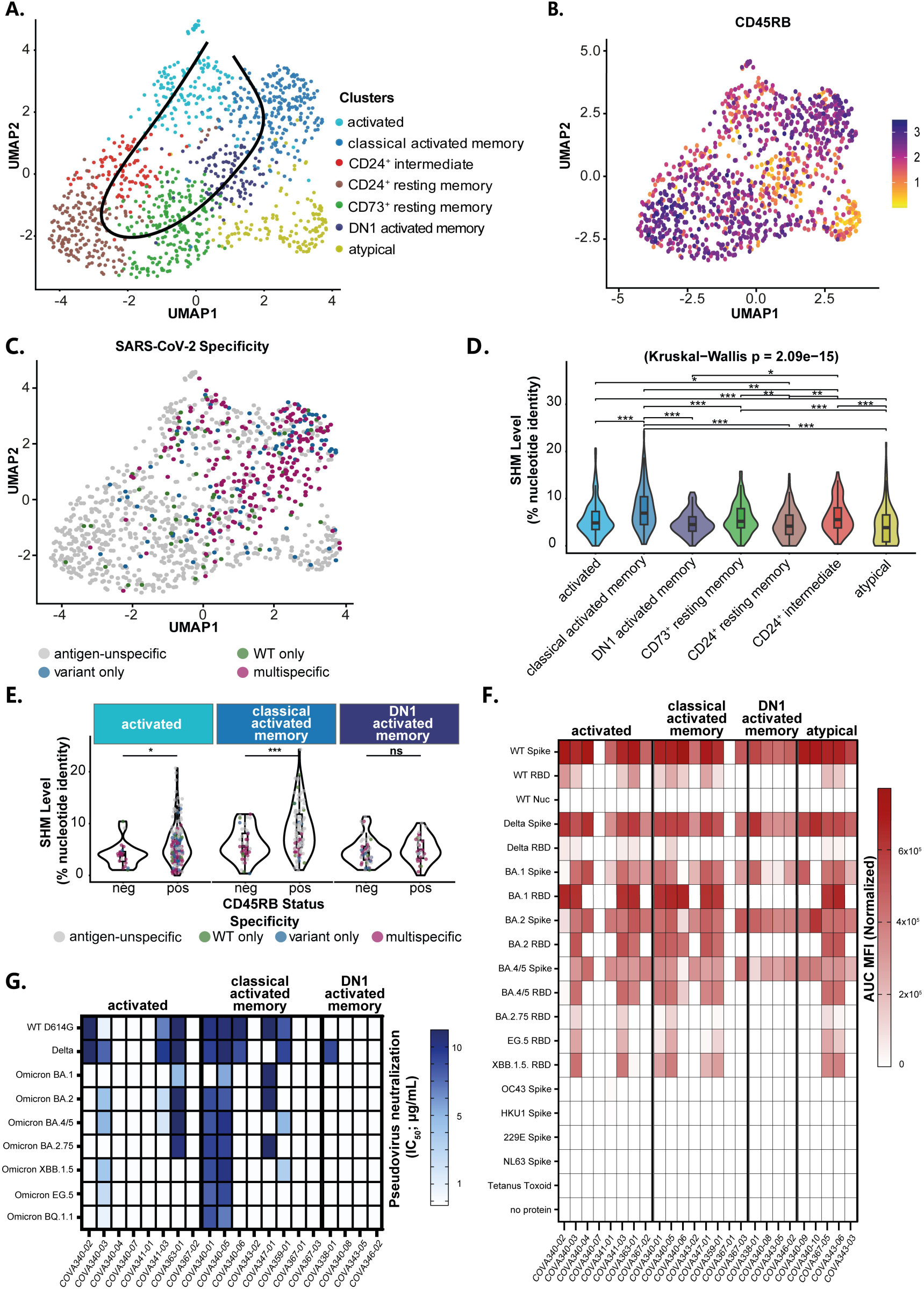
Trajectorial, specificity, and SHM analysis of the MBC compartment and mAb binding and neutralization validation. **(A)** UMAP plot of the MBC compartment showing the lineage trajectorial relationships across the different subpopulations. **(B)** scaled CD45RB expression within the MBC compartment as defined by CITE-seq analysis. **(C)** UMAP plot of the MBC compartment visualizing B cell antigen specificity across the MBC clusters. **(D)** SHM frequency within the IGHV gene separated by B cell subset. Pairwise comparison between B cell subsets within the CD45RB positive and negative compartment can be found in Table S7. Significance is indicated as * <0.05, **<0.01, ***<0.001, **** <0.0001. **(E)** SHM frequency within the IGHV gene separated by B cell subset and CD45RB expression. Cells were separated based on CD45RB expression within every B cell subset and a Wilxocon test was performed to detect any differences in SHM levels between CD45RB positive and negative B cells. Pairwise comparison between B cell subsets within the CD45RB positive and negative compartment can be found in Table S6. Significance is indicated as * <0.05, **<0.01, ***<0.001, **** <0.0001. **(F)** Binding activity of the produced mAbs, separated by MBC cluster, as assessed by our in-house Luminex assay^34^. The color scale shows the normalized area under the curve (AUC) values of measured mean fluorescence intensities (MFI). Beads coupled to the tetanus toxoid protein and uncoupled beads were included as negative controls **(G)** Pseudovirus neutralization potencies (IC50; μg/mL) of all produced mAbs, divided per MBC cluster.

CD45RB expression, a marker associated with previous GC participation^55^, further revealed parallel GC-associated and non-GC-associated differentiation profiles. Although classical Act-Mem and ActB were enriched for CD45RB⁺ cells, CD45RB⁺ and CD45RB⁻ populations were detected throughout the ActB, CD24⁺ Int, and RestMem compartments, whereas DN1 Act-Mem consisted predominantly of CD45RB⁻ B cells (Figure 3B, Supplementary Figure S7A). These findings indicate that GC-associated and extrafollicular differentiation profiles are maintained in parallel across multiple stages of IgG⁺ B cell differentiation rather than defining discrete memory subsets.

Analysis of the antigen-specific B cells using barcoded SARS-CoV-2 spike dextramers supported the proposed trajectory. Antigen-specific B cells were enriched within ActB, classical ActMem, and DN1 ActMem populations, reflecting recently activated recall and *de novo* responses following SARS-CoV-2 breakthrough infection (Figure 3C, Supplementary Figure S7B)^56^. In contrast, antigen-specific B cells remained scarce within the resting memory compartments at this early stage of the response, one month post breakthrough infection, supporting the model that activated populations progressively transition toward resting memory.

SHM analysis further supported the inferred developmental relationships. ActB and classical ActMem exhibited the highest SHM frequencies, consistent with recall responses and preferential GC origin. SHM levels were lower within the RestMem compartment and CD24^+^ Int subset, with CD73⁺ RestMem and CD24⁺ Int displaying intermediate mutation frequencies, both exceeding those observed in CD24⁺ RestMem cells (Figure 3D). In contrast, DN1 Act-Mem and AtypB showed substantially lower SHM levels, consistent with a predominantly extrafollicular origin. Of note, within subset enriched for spike-specific B cells, the ActB and classical ActMem carried significantly higher SHM in their CD45RB+ than CD45RB⁻ B cells, further supporting the association between CD45RB and previous GC participations (Figure 3E, Supplementary Figure S7C, Supplementary Table S7).

Together, these analyses provide insights in the developmental relationships between the major IgG⁺ MBC populations, positioning ActB as the principal activated state, ActMem as a transitional compartment, CD73⁺ and CD24⁺ RestMem as quiescent endpoints, and CD24⁺ Int as a potential re-entry state linking resting memory to activation, while revealing parallel GC-associated and extrafollicular differentiation profiles across this continuum.

To determine whether these developmental states also differed functionally, we characterized monoclonal antibodies (mAbs) derived from SARS-CoV-2-specific B cells sorted from populations enriched for antigen-specific cells — namely ActB, classical ActMem, and DN1 Act-Mem. (Figure 3F-G). Spike and RBD proteins from WT as well as breakthrough variants (Omicron and Delta) were used to determine primary vaccine responses (WT only), recall responses (WT and breakthrough cross-reactive), and *de novo* responses (breakthrough only).

mAbs derived from ActB and classical ActMem showed the broadest binding across WT and Omicron-lineage spike antigens, preferentially targeted the RBD as confirmed by ACE2 competition Supplementary Figure S8), and exhibited the greatest virus neutralization potency, consistent with their high SHM levels and enriched origin from CD45RB⁺ B cells. In contrast, DN1 ActMem-derived mAbs were restricted to non-RBD, spike epitopes and largely lacked neutralizing activity, consistent with their low SHM frequencies and predominantly suspected extrafollicular origin.

Together, these findings demonstrate that the developmental states and inferred developmental origins defined by single-cell analyses are associated with distinct epitope specificities, and neutralization capacities within the SARS-CoV-2-specific IgG⁺ B cell response.

### Longitudinal spectral flow cytometry analysis validates detailed B cell clusters identified by single cell RNAseq

To validate the proposed developmental relationships between IgG⁺ B cell subsets, we longitudinally tracked SARS-CoV-2 spike-specific B cells by spectral flow cytometry. Samples were derived from a different cohort, the RECoVERED cohort, in which participants acquired natural infection with the ancestral (WT) strain followed by two mRNA vaccine doses. Multiple blood samples were collected from 10 individuals at three intervals: (i) 1 week to 8–10 months post-infection, (ii) 1 week to 1 month after the first vaccine dose, and (iii) 1 week to 8 months after the second (booster) dose (Figure 4A). SARS-CoV-2 spike-specific IgG⁺ B cells expanded following the first vaccine dose, administered approximately 10 months after infection, and gradually contracted between 1 and 8 months after booster vaccination, providing a longitudinal framework to track transitions from recently activated to resting memory B cell states (Figure 4B, Supplementary Table S8).

**Figure 4.**
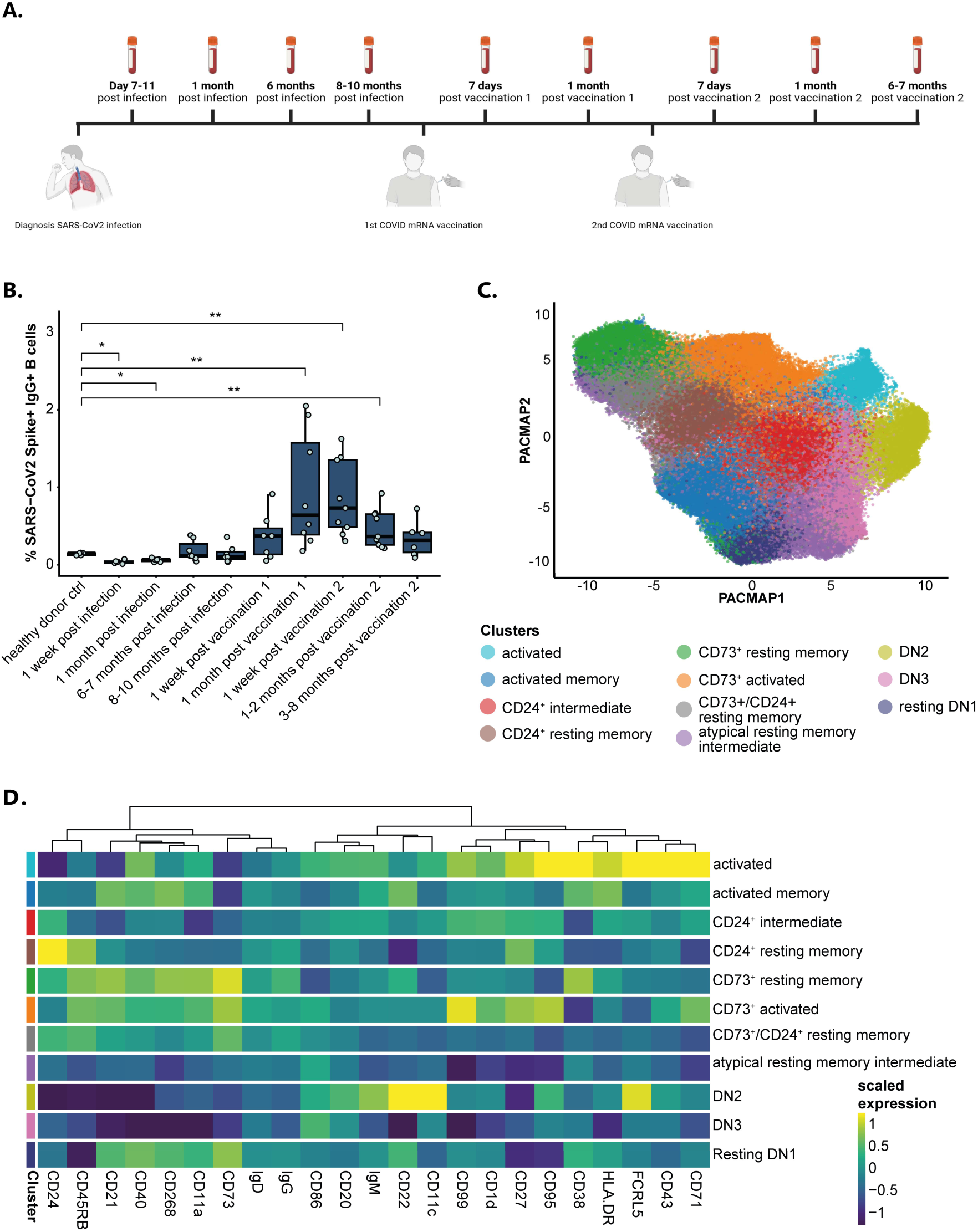
Longitudinal Spectral Flow Cytometry analysis of IgG⁺ MBC after infection and vaccination. **(A)** Schematic overview of available samples from the RECoVERED study including individuals infected with SARS-CoV-2 followed by two vaccinations. Created in BioRender. Van Gils, M. (2026) https://BioRender.com/si5f0wt **(B)** Boxplot representing frequency of IgG⁺ SARS-CoV-2⁺ IgG⁺ B cells out of total B cells. Wilcoxon test was performed comparing healthy donor ctrl with all timepoints individually. Pairwise comparisons between timepoints can be found in Table S8. Significance is indicated as * <0.05, **<0.01, ***<0.001, **** <0.0001. **(C)** PACMAP showing cluster annotation within the IgG⁺ memory compartment. Dimensionality reduction was performed with PACMAP and cell clustering was done on the PCAs using the Leiden algorithm. Clusters were annotated based all cell surface protein markers excluding Ig isotype markers. **(D)** Heatmap of scaled cell surface marker expression per cluster. Markers are ordered based on expression pattern similarities. Differential gene expression analysis can be found in Table S9

Guided by our multimodal single-cell RNAseq (ScRNAseq) analysis, we expanded our previously published spectral flow cytometry panel^33^ to 29 surface markers, including CD99, CD11a, CD1d, CD40, CD22, and BAFF-R. PACMAP dimensionality reduction and unsupervised clustering were applied to a composite dataset of antigen-specific IgG⁺ B cells supplemented with a fixed number of antigen-unspecific IgG⁺ B cells per sample to provide biological context, enabling reconstruction of the phenotypic landscape of the IgG⁺ B cell response (Figure 4C, Supplementary Figure S9-10, Supplementary Table S9). This analysis recapitulated all major IgG⁺ B cell populations identified by multimodal CITE-seq analysis while providing additional phenotypic resolution. As a result, B cell compartments were resolved into additional phenotypically distinct B cell subsets, resulting in the previously established four major compartments comprising 11 subsets (Figure 4C-D). These included a RestMem compartment (CD73⁺ RestMem, CD24⁺ RestMem, CD73⁺CD24⁺ RestMem, and resting DN1), an intermediate activated compartment (ActMem, CD73⁺ activated (CD73⁺ ActB), and CD24⁺ Int), an activated compartment (ActB), and an AtypB compartment (AtypB resting intermediate, DN2, and DN3).

Subset identities were confirmed by conserved expression of the defining markers identified by both the literature and multimodal CITE-seq analysis (Figure 4D, Supplementary Figure S7). As a result, IgG⁺ B cell clusters were characterized by their surface marker expression patterns (Figure 4C–D, Supplementary Figure S9-10, Supplementary Table S9).

Accordingly, ActB were defined by high CD27 and CD38 expression, loss of CD21, and upregulation of CD43, CD71, CD95, and CD99. Differential expression of CD45RB, CD11c, FcRL5, and CD86 within this compartment indicated heterogeneity in origin and state (Supplementary Figure S9-10)^57^. The RestMem compartment was characterized by CD21 expression and absence of activation markers CD71, CD86, and CD43, and resolved into four subsets. CD73⁺ RestMem, CD24⁺ RestMem, and CD73⁺CD24⁺ RestMem were subdivided based on CD73 and/or CD24 expression, with CD73⁺ RestMem showing higher CD38, CD40, BAFF-R, and CD11a than CD24⁺ resting DN1 shared this resting memory phenotype (CD21⁺, CD38⁺, CD40⁺, BAFF-R⁺) but lacked CD27 and CD45RB, consistent with a non-classical resting memory identity. The three transitional populations bridged the activated and resting compartments. ActMem showed similar expression as classical ActMem expressing RestMem and activation markers (CD21, CD22, CD11a, CD71, CD43, CD38). CD73⁺ ActB expressed CD73 alongside activation markers (CD86, CD95, CD43, CD99) and CD21, and was uniformly CD45RB^+^. CD24⁺ Int were defined by CD24 and CD1d with intermediate activation marker levels (CD71, FcRL5, CD95) and reduced resting memory markers (CD21, BAFF-R, CD40, CD11a). Similar to our scRNAseq data, CD45RB heterogeneity was also observed within the ActMem and CD24⁺ Int populations, consistent with coexistence of GC-dependent and extrafollicular cells within these transitional states. The AtypB compartment lacked CD27, CD45RB, and CD21 and comprised three subsets: atypical resting intermediate (CD21⁺ ^/-^, CD40⁺), displaying intermediate levels of resting memory markers; DN2 (FcRL5⁺, CD11c⁺, CD22⁺), exhibiting the highest expression of activation markers (CD95, CD86, and CD43); and DN3 (FcRL5⁻, CD11c⁻, CD40⁻), displaying reduced activation marker expression, defining phenotypically distinct states within the AtypB compartment.

Together, these findings demonstrate that the phenotypic organization of the IgG⁺ memory B cell compartment is reproducibly resolved by spectral flow cytometry, independently validating the populations identified by multimodal scRNAseq. Notably, the increased resolution enabled more detailed characterization of the RestMem compartment into classical RestMem and DN1 RestMem, and of the AtypB compartment into DN2 and DN3 subclusters. This panel further enables longitudinal tracking of the dynamics of all IgG⁺ memory B cell subsets following antigen re-exposure.

### Longitudinal analysis identifies multiple phenotypic B cell stages present at different timepoints after vaccination revealing B cell dynamics and fate

Next, we analyzed longitudinal dynamics of spike-specific B cell populations following infection and vaccination to validate and extend the differentiation trajectory inferred from our scRNAseq analysis. We focused our analysis on the first and second vaccine doses, where we observed pronounced expansion and contraction of IgG⁺ spike-specific B cells, providing well-defined recall responses to interrogate B cell re-activation and memory formation dynamics (Figure 4B).

Initial comparisons across chronological timepoints revealed substantial inter-individual heterogeneity in spike-specific B cell population frequencies, reflecting differences in response timing across donors (Supplementary Figure S11). To account for this, samples were grouped by similarity of B cell cluster frequency profiles using a correlation-based clustering approach, aligning individuals by biological activation state rather than chronological time points. This yielded five phenotypically distinct stages defined by their dominant cluster composition and preferential association with vaccination timepoints: resting, early intermediate 1, early intermediate 2, activated, and late intermediate (Figure 5A).

**Figure 5.**
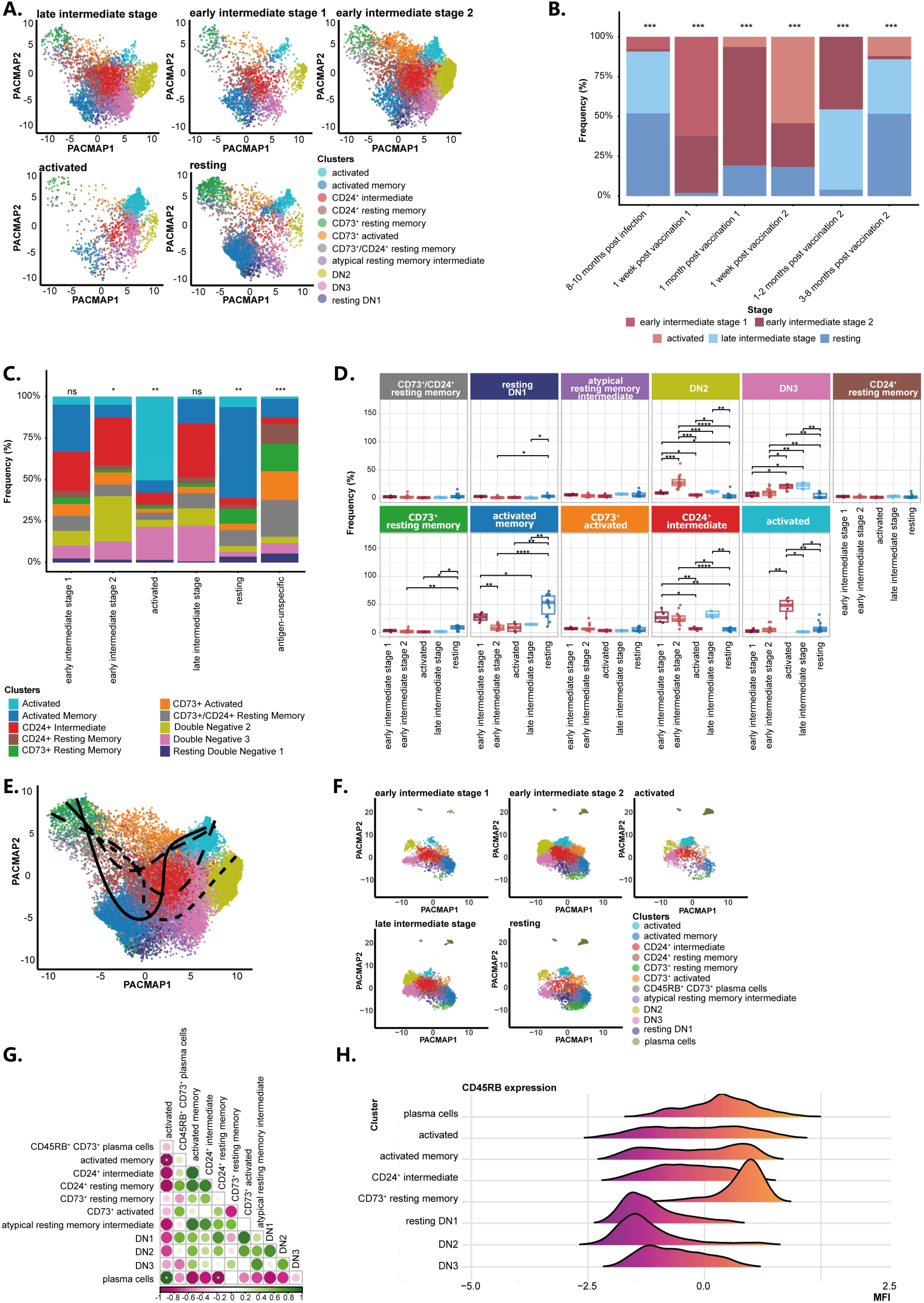
Phenotypic stage and B cell trajectory analysis of the IgG^+^ MBC compartment. **(A)** PACMAP of 5 distinct phenotypic stages found in the IgG+ memory compartment determined by correlation analysis of cluster frequencies between timepoints and individuals. **(B)** Phenotypic stage frequency per timepoint. A mixed model analysis was performed to compare stage frequencies per timepoint. Pairwise comparisons can be found in Table S10. Significance was indicated as * <0.05, **<0.01, ***<0.001, **** <0.0001. **(C)** Composition of B cell phenotypes found per phenotypic stage. Pairwise comparisons can be found in Table S11. Significance was indicated as * <0.05, **<0.01, ***<0.001, **** <0.0001. **(D)** Frequency of B cell clusters found per stage. A pairwise wilcoxon test was performed to determine significant differences in frequency between stages. P-values were adjusted for multiple testing using the Benjamini-Hochberg method. Significance was indicated as * <0.05, **<0.01, ***<0.001, **** <0.0001. **(E)** PACMAP with trajectory commonly found between donors. Trajectorial lineages were inferred using the Slingshot algorithm with longitudinal data embedded in the pseudotime. (**F)** PACMAP of IgG+ memory B cells including ASCs per phenotypic stage. Dimensionality reduction was performed with PACMAP and clustering was done on the PCAs using the Leiden algorithm. Cluster annotation was done based on expression of cell surface markers. The 5 phenotypic stages were previously determined using the IgG^+^ MBC dataset and assigned to the ASCs based on their sample origin. **(G)** Pearson correlation of cluster frequencies in the activated stage within the IgG^+^ memory compartment with plasma cells Significance was indicated as * <0.05, **<0.01, ***<0.001, **** <0.0001. **(H)** Histogram of CD45RB expression in antigen-specific associated clusters.

These stages showed distinct distributions across vaccination time points (Figure 5B, Supplementary Table S10), revealing a coordinated progression of B cell states following antigen exposure. The resting and late intermediate stages predominated at 8 months post-infection and 3-8 months post second vaccination, with late intermediate represented at 1–2 months post second vaccination. early intermediate 1 was dominant at 1 week post first vaccination, early intermediate stage 2 at 1 month post first vaccination, and the activated stage at 1 week post second vaccination, consistent with a faster and stronger recall response following the booster dose.

We next examined the cluster composition of each stage to understand the cellular basis underlying these dynamic responses (Figure 5C-D, Supplementary Table S11). CD73⁺ Rest-Mem, resting DN1, as well as ActMem subsets, were more abundant in the resting stage, in line with their identity as memory populations persisting between antigen exposures^31^. The early intermediate 1 stage was characterized by the disappearance of CD73⁺ RestMem and resting DN1 and a decline of ActMem subsets, coinciding with increased frequency of the CD24⁺ Int population. This validates our multi-omics analysis that identified the CD24⁺ Int subset as an early recall MBC phenotype. At early intermediate 2 stage ActMem clusters fully disappear while CD24⁺ Int remained stable and DN2 B cells reach peak frequency. The activated stage was characterized by an increase in ActB and DN3 cells and a decline of DN2 B cells and CD24⁺ Int. This was followed by a sharp decline in ActB cells and resurgence of re-activated populations CD24⁺ Int and DN2 B cells in the late intermediate stage, potentially reflecting a second wave of reactivation following the booster dose. The changes in cluster frequency observed within spike-specific B cells were mirrored when clusters were expressed as a proportion of total B cells (Figure 5D, Supplementary Figure S12), indicating that the stage-associated dynamics reflect true expansion and contraction of antigen-specific populations rather than redistribution within the IgG⁺ compartment.

To distinguish antigen-driven dynamics from steady-state memory composition, we compared the frequencies of spike-specific B cell clusters across activation stages with those of the unspecific IgG⁺ memory B cell compartment (Figure 5C-D, Supplementary Figure S12, Supplementary Table S11). spike-specific B cells were significantly enriched within the ActB, Act-Mem, CD24⁺ Int, and DN2 subsets, depending of the activation stages, whereas the antigen-unspecific B cell compartment was predominantly composed of CD73⁺ RestMem, CD24⁺ RestMem, CD73⁺CD24⁺ RestMem, resting DN1, AtypB resting intermediate, and CD73⁺ ActB. CD24⁺ RestMem and CD73⁺CD24⁺ RestMem showed little to no spike-specific enrichment at any stage, indicating that these populations contribute minimally to early antigen-driven recall response. In contrast, CD73⁺ RestMem remained consistently detectable among spike-specific B cells, particularly during the resting stage, consistent with their proposed identity as long-lived memory cells^33^. Surprisingly, CD73⁺ ActB represented a substantial fraction of unspecific memory compartment despite their activated phenotype, yet displayed only low persistent representation within the spike-specific compartment across all stages.

To further examine the relationships between the temporally defined B cell states, we performed trajectory analysis incorporating the longitudinal data (Figure 5E). The CD73⁺ RestMem was selected as the trajectory root, as this subset was identified as the most persistent antigen-specific memory population. Trajectory reconstruction identified multiple potential activation paths linking the major spike-specific B cell populations. The pathway encompassing CD24⁺ Int and ActB was consistent with their sequential appearance during the longitudinal recall response. Additional paths involving ActMem, resting DN1 and AtypB populations were also identified.

Next, we investigated the relationship between B cell populations and ASC differentiation by incorporating ASCs into the longitudinal dataset (Figure 5F-G, Supplementary Figure S13-14). ASCs formed a cluster immediately adjacent to the ActB compartment, consistent with a close phenotypic relationship between these populations. ASCs peaked during the activated stage and positively correlated with ActB frequency, whereas ActB negatively correlated with Act-Mem. Together, these findings support ActB as the principal activated effector population immediately preceding ASC differentiation during the recall response.

In line with our scRNAseq data, all populations along the differentiation path, ActB, ActMem, CD24⁺Int, and resting memory compartments resolved into CD45RB⁺ and CD45RB⁻ subsets, as did the ASC population (Figure 5H), suggesting that both GC-associated and extrafollicular developmental origins are maintained throughout differentiation toward memory and antibody-secreting cell fates.

Collectively, these longitudinal analyses using an independent cohort refine the differentiation model inferred from multimodal single-cell analysis by describing the temporal dynamics of the major IgG⁺ memory B cell populations during human recall responses. Whereas the persistent CD73⁺ RestMem and the transient ActMem populations both participated in antigenspecific recall, CD24⁺ RestMem and CD73⁺CD24⁺ RestMem remained largely excluded from the response, identifying functionally distinct resting memory compartments. Longitudinal dynamics further positioned CD24⁺ Int as an early recall-associated state preceding expansion of the ActB compartment, which represented the principal activated population immediately before ASC differentiation. The persistence of both CD45RB⁺ and CD45RB⁻ populations throughout the recall response further indicates that GC-associated and extrafollicular-associated profiles are maintained across activation and differentiation. Finally, early DN2 and late DN3 populations exhibited coordinated temporal dynamics during recall, indicating that they actively participate in the antigen-specific response, although their developmental relationship to the classical memory B cell pathway remains unresolved. Together, these findings provide a temporally resolved framework for human memory B cell re-activation following infection and vaccination.

## Discussion

To understand how the human IgG⁺ memory B cell compartment is organized during recall responses, we combined multimodal single-cell profiling with longitudinal tracking of SARS-CoV-2-specific memory B cells following infection and vaccination. Our findings assign biological functions and developmental relationships to previously described and newly identified B cell populations, transforming descriptive subsets into a coherent model of human memory B cell recall.

Within this framework, ActB represents the activated effector state; ActMem, the transitional population linking effector B cells to long-lived resting memory; CD73⁺ RestMem, the principal long-lived recall reservoir; and CD24⁺ Int, the transitional population through which memory re-enters the recall response. Together, trajectory inference and longitudinal analyses define the directional transitions connecting these functional states, identifying ActMem and CD24⁺ Int as the two principal intermediates coordinating memory formation and recall.

In parallel, our data show that these functional states are traversed by two conserved developmental profiles. classical (CD45RB⁺) and non-classical (CD45RB⁻) memory B cells, previously proposed to arise predominantly through GC and extrafollicular responses, respectively, progress through analogous functional states while maintaining distinct developmental histories^31,32,55,58^. In contrast, AtypB follow a related but separate differentiation pathway. This three-pathway resolution extends prior B cell activation frameworks by showing follicular and extrafollicular fates co-exist across shared activation phenotypes. The application of extensive cell-surface marker expression allowed us to define RestMem, activated, as well as transitional memory B cell subsets at high resolution and assign functionality of B cell subsets throughout re-activation.

ActB have been described as transient populations that emerge following infection or vaccination and have been implicated in recall responses, ASC differentiation and ongoing germinal center reactions^33,37,59–63^. However, their biological role within the memory B cell response has remained unclear. Our data identify ActB as a transient activated B cell subset that represents the effector state of the IgG⁺ memory B cell response. In contrast to resting memory B cells, ActB undergo a marked functional shift from a homeostatic survival profile sustained by BAFF-mediated signals and CD40-dependent T cell interactions toward an inflammatory profile. They acquire inflammatory responsiveness together with the molecular machinery required to migrate to inflamed tissues and secondary lymphoid organs, interact with T cells and differentiate into ASCs. Notably, several components of this inflammatory trafficking profile, including CXCR3 and integrins, have previously been associated with pathogenic B cell populations in autoimmune and chronic inflammatory diseases, suggesting that the physiological effector profile defined here may provide a framework for interpreting disease-associated B cell responses^64–66^. Our data further indicate that this transient effector state can arise through multiple developmental routes. During recall responses, contraction of the resting memory compartment, together with lineage inference and the high levels of SHM observed in ActB, supports their derivation from re-activated memory B cells, consistent with the model proposed by Sokal et al^38,67^. However, the persistence of a small CD45RB⁺ ActB population after immune contraction is also consistent with continued GC output described previously^68,69^. Thus, rather than representing a lineage restricted to a single developmental pathway, ActB defines a convergent effector state generated by both memory recall and ongoing GC responses. Importantly, our data indicate that the ActB state is not a terminal differentiation endpoint. Together with lineage inference and the emergence of persistent ActMem populations, our findings reconcile previous models proposing ActB as either ASC precursors or memory B cell precursors by demonstrating that ActB cells contribute to both antibody-secreting cell differentiation and memory regeneration^70,71^.

Our data position ActMem as a persistent effector-memory population that preserves rapid recall potential beyond the acute response. ActMem expand as ActB contract following the acute response, persists for at least eight months after antigen exposure, and are selectively depleted upon antigen re-encounter, indicating that it represents a durable antigen-experienced reservoir poised for rapid reactivation. Together with our previous observation that Act-Mem are largely absent from long-established antigen-specific memory B cells despite being enriched among recently generated antigen-specific memory B cells and scarce within the antigen-unspecific compartment^33^, these findings indicate that the ActMem compartment represents a persistent, yet ultimately transitional, stage of the recall response rather than prolonged activation. ActMem retain inflammatory responsiveness and migratory capacity while progressively reacquiring follicular homing and homeostatic survival profiles. Together with lineage inference, positioning ActMem between ActB and RestMem, these findings support a model in which recently activated effector B cells transition through ActMem before re-establishing long-lived resting memory while preserving rapid recall competence. Thus, whereas CD73⁺ RestMem maintain durable humoral immunity, ActMem preserve effector competence during the transition back to memory, resembling the functional organization of effectormemory T cells.

Our previous work established CD73⁺ and CD24⁺ RestMem as the principal resting IgG⁺ memory B cell compartments^33^. Both subsets share the molecular hallmarks of durable memory, including follicular recirculation, homeostatic survival and translational readiness, supporting their role in the long-term maintenance of humoral immunity. However, they differ markedly in their contribution to antigen-specific memory, recall responses and immune regulation.

The CD73⁺ RestMem subset represents the principal recall-competent compartment of longlived humoral memory. Previous studies demonstrated the remarkable persistence of CD73⁺ RestMem decades after smallpox vaccination in the spleen^31,72^, while our previous work showed that long-established antigen-specific memory B cells against pathogens such as, influenza virus and RSV and Tetanus toxoid vaccines are predominantly composed of CD73⁺ B cells. Here, we extend these observations by demonstrating that CD73⁺ RestMem are already enriched within the antigen-specific RestMem compartment three to eight months after infection or vaccination. Moreover, CD73⁺ RestMem are mobilized early following antigen reencounter, identifying this population as one of the principal long-lived reservoirs sustaining secondary humoral immune responses^56^. Consistent with this role, CD73⁺ RestMem retain a homeostatic phenotype characterized by stronger follicular maintenance and survival profiles while remaining readily poised for rapid recall.

In contrast, CD24⁺ RestMem contribute minimally to the SARS-CoV-2 recall response despite remaining abundant within the total RestMem compartment. Phenotypically, CD24⁺ RestMem display a distinct resting memory profile characterized by enhanced inhibitory and environmentally responsive pathways, several of which are shared with AtypB, while acquiring inflammatory responsiveness and activation-associated characteristics, consistent with progression toward the ActB effector state. Expression of CD1d, CD1c, HLA-E and multiple inhibitory receptors further suggest overlap with innate-like or regulatory B cell profiles^73,74^, although their precise functional identity remains to be established. These findings suggest that CD24⁺ Rest-Mem represent a biologically distinct resting memory profile, either constituting a deeply quiescent memory compartment with a higher threshold for reactivation or a specialized regulatory memory population.

The identification of the CD24⁺ Int subset represents one of the principal advances of this study. Although transient CD24-expressing ActB populations have previously been described following vaccination, their biological significance has remained unclear^37^. By integrating trajectory inference with longitudinal analyses, we identify the CD24⁺ Int subset as an early transitional state that emerges as RestMem and ActMem populations contract, preceding the expansion of ActB and ASCs. These findings support a model in which CD24⁺ Int represent the principal entry point into the recall response through which RestMem are mobilized following antigen re-encounter. Whether CD24⁺ Int are restricted to secondary immune responses or also contributes to primary responses remains unknown. Likewise, although its temporal dynamics and developmental relationships strongly support derivation from RestMem, definitive lineage relationships will require future clonal lineage-tracing studies.

AtypB closely resembled the well-described atypical memory B cell compartment reported across infection, vaccination and autoimmune diseases^75,76^. Longitudinal analyses further refined the temporal organization of this compartment. DN2 B cells expanded early during recall responses, following the emergence of CD24⁺ Int but preceding the expansion of ActB, and displayed an activated yet inhibitory phenotype. In contrast, DN3 B cells accumulated later during immune contraction, suggesting that these populations represent temporally distinct stages of the AtypB response. Nevertheless, trajectory inference positioned AtypB outside the principal ActB–ActMem–RestMem pathway, supporting the view that AtypB follow a differentiation profile largely independent of the conventional memory B cell response^77^.

Previous studies demonstrated that CD45RB expression distinguishes classical and non-classical memory B cell lineages and remains stable after activation, allowing activated progeny to be traced back to their developmental origin^55,58,78^. Here, we show that these developmental profiles are maintained throughout the entire recall trajectory. GC- and extrafollicular-derived B cells progress through analogous functional states—including resting memory, activation, activated memory and plasmablast differentiation—while preserving distinct developmental identities^79,80^. SHM analyses independently supported these developmental histories. Classical ActMem consistently displayed higher mutation levels than DN1 ActMem, whereas the small CD45RB⁻ fraction within the ActB compartment similarly exhibited lower mutation levels than their CD45RB⁺ counterparts. This is in line with previous reports showing the activation of B cells through different pathways, identifying DN1 B cells as non-GC derived with lower SHM levels and ActB as GC-derived affinity matured responses^25,57,67^.

Consistent with these distinct developmental histories, the two ActMem populations also retained different functional characteristics. classical ActMem preferentially targeted RBD epitopes and generated the most potent neutralizing antibodies, whereas DN1 ActMem were enriched for non-RBD specificities^81^. This extends our previous observation that non-classical ActMem and ActB preferentially recognized SARS-CoV-2 nucleocapsid over spike, suggesting that developmental origin influences both antibody maturation and epitope selection, as expected^33,82^. In addition, classical ActMem displayed a more activated and migratory profile, whereas DN1 ActMem exhibited a more environmentally responsive and regulatory phenotype, including preferential expression of type 2 profile (IL4R, IL23R) and multiple regulatory genes. Whether these distinct profiles also confer specialized migratory behavior, tissue localization, or recall functions, including potential roles in diseases such as allergy responses mediated type 2 immunity^54^, remains an important question for future studies.

In summary, we establish a unified model of the human IgG⁺ memory B cell response that illustrates how humoral memory is generated, maintained and reactivated. CD24⁺ Int define the principal entry point through which RestMem initiate recall responses, whereas the Act-Mem population links the effector response to the regeneration of long-lived memory while remaining poised for rapid reactivation. Together, these previously unrecognized transitional populations connect activation, memory maintenance and recall, transforming descriptive B cell subsets into a coherent framework for human memory B cell biology.

Beyond defining the functional and developmental organization of human memory B cells, our integrated transcriptomic and surface protein atlas provides a high-resolution reference for identifying and comparing IgG⁺ memory B cell populations across physiological and pathological immune responses. Together, this framework provides both a conceptual model of human memory B cell recall and a community resource for studying B cell responses in vaccination, chronic infection, autoimmunity, allergy and other immune-mediated diseases.

## Materials and methods

### Study population

For the analysis of single B cells by 10x Genomics, samples were obtained from 12 individuals included in the COVID-19-specific antibodies (COSCA) cohort^35^, which was conducted at the Amsterdam University Medical Center, location AMC, the Netherlands, and was approved by the local ethical committee (NL73281.018.20). Written informed consent was provided by all participants before being enrolled in the study. Peripheral blood mononuclear cells (PBMCs) were collected from the individuals 4 to 7 weeks after symptom onset. The individuals received one or multiple doses of a COVID-19 mRNA vaccine (Comirnaty from Pfizer/BioNTech or Spikevax from Moderna). Only one donor, COSCA-365, was first vaccinated twice with the viral vector-based Vaxzevria vaccine from AstraZeneca, followed by a third vaccination with Spikevax. Following vaccination, these individuals experienced a breakthrough infection with one SARS-CoV-2 variant (6 infected with Delta, 3 infected with Omicron BA.1/2 and 3 infected with Omicron BA.4/5), as shown by a PCR-confirmed infection or determined by a high likelihood based on the dominant strain circulating in the Netherlands at the time of infection.

Flow cytometry analysis was performed using 10 individuals included in the RECoVERED cohort conducted at the Amsterdam University Medical Center, location AMC, the Netherlands, and the Public Heatlh Service of Amsterdam (GGD Amsterdam)^83^ and was approved by the medical ethical review board of the Amsterdam University Medical Centers (NL73759.018.20). All participants provided written informed consent. Individuals were infected with PCR-confirmed SARS-CoV-2 WT followed by a 2-dose COVID-19 mRNA vaccine regimen (Comirnaty from Pfizer/BioNTech or Spikevax from Moderna). PBMCs were collected 1 week to 10 months following SARS-CoV-2 WT infection, 1-4 weeks after the first vaccination, and 1 week to 10 months after boost vaccination. participant characteristics are reported in Supplementary Tables S1 and S2.

### Soluble SARS-CoV-2 protein design, production and purification

Soluble avidin/hexahistidine-tagged SARS-CoV-2 S proteins were stabilized in the prefusion conformation by introducing two prolines (S-2P) and a T4 trimerization domain, as described previously^34,35^. The wild-type (WT) SARS-CoV-2 S sequence (Wuhan Hu-1, GenBank: MN908947.3) was used as a template sequence to introduce mutations and produce S proteins of the other variants. Soluble RBD protein constructs of the corresponding variants (residues 319-541) containing the same mutations were ordered as gBlock gene fragments (Integrated DNA Technologies) and cloned into a pPPI4 vector by PstI/BamHI digestion and ligation, using a Gibson Assembly protocol optimized in-house (Thermo Fisher Scientific). Sanger sequencing was employed to check the sequences of all constructs.

Soluble proteins were then produced in human embryonic kidney (HEK)293F cells (Thermo Fisher Scientific), maintained in FreeStyle medium (Gibco). HEK293F cells were transfected with a 1:3 ratio of expression DNA plasmids and polyethyleneimine hydrochloride (PEI)max (1 µg/mL) in OptiMEM. Transfected HEK293F cells were cultured at 37°C for 6 days, after which supernatants containing the produced soluble SARS-CoV-2 proteins were harvested, centrifuged at 4000 rpm for 30 min and filtered using 0.22 µm SteriTop filters (Merck Millipore). Affinity chromatography with Ni-NTA agarose beads (Qiagen) was used for protein purification. Proteins were eluted, concentrated, and buffer exchanged to phosphate-buffered saline (PBS) or TN75 buffer (75 mM NaCl and 20 mM Tris HCl, pH 8.0) using VivaSpin20 filters (Sartorius). The concentration of the proteins produced was determined by measurement at the Nanodrop

### Probe labeling methods for SARS-CoV-2-specific B cell selection

All SARS-CoV-2 proteins (WT S, Delta S, Omicron BA.1 RBD, Omicron BA.2 RBD, Omicron BA4/5 S, H1N1 trimer) were prepared in two different colors to detect double positive SARS-CoV-2-specific B cells and eliminate any fluorochrome-specific B cells. SARS-CoV-2 dextramer probes were prepared using the (dCode) Klickmer products from Immudex. First, proteins were incubated with either PE dCode Klickmer or APC Klickmer (Immudex) in a 5:1 protein to dextramer ratio for 1 h in the dark at 4°C. Next, 10 µM biotin solution was added. Probes were incubated with biotin for 30 min in the dark at 4°C. After this step, the different colour probes were mixed and used for cell staining.

Probes were tested by flow cytometry bead assay or PBMC staining for their binding abilities. For the bead assay, anti-mouse Ig/negative control CompBeads (BD Biosciences) were stained with 0.125 µg mouse anti-human Ig primary antibody, followed by 0.5 µg of the antibody to test. For PBMC staining, isolated PBMCs from healthy donor buffy coats were thawed briefly in the water bath at 37°C and placed in 40 mL RPMI supplemented with 20% FCS. Next, SARS-CoV-2 probes were added to either the beads or PBMCs and incubated for 30 min. Beads or cells were washed with FACS buffer. Next, cells were stained with a B cell phenotype antibody panel for FACS detection of B cell populations (CD4 eF780, CD3 eF780, CD14 eF780, CD16 eF780, viability dye eF780, CD19 AF700, IgD BV785, IgG BV605, CD27 BV421) for 30 min at 4°C in the dark. After incubation, cells were washed with FACS buffer. Cells and beads were analyzed using flow cytometry analysis on the BD FACS Aria IIu SORP 4 laser sorter or BD Fortessa machine.

### Single-cell RNA sequencing of B cells by 10x Genomics

First, PBMC samples were thawed in separate Falcon tubes containing 40 mL thaw medium (RPMI medium supplemented with 20% fetal calf serum (FCS)) by placing them in the water bath at 37°C until they were partially defrosted and later transferred into the pre-warmed thaw medium. Cells were centrifuged at 1600 rpm for 5 min, after which the supernatant was carefully removed by decanting without disturbing the cell pellet. Cell pellets were resuspended in FACS buffer (2% FCS and 1 mM EDTA), adjusted to approximately 2x10^6^ cells/mL and centrifuged at 800 rcf for 5 min. After the centrifugation step, the supernatant was discarded and 37.5 µL of 1:10 diluted TruStain FcX solution (BioLegend) was added to each sample. Cells were incubated in the dark at 4°C for 10 min to allow Fc receptor blocking. In the meantime, TotalSeqC hashtag antibody (BioLegend) mixes (Supplementary Table S6) were prepared according to the protocol from the manufacturer, and then incubated with the PBMC samples for 30 min, in the dark at 4°C. After incubation, 1 mL of FACS buffer was added to the samples, which were then centrifuged at 800 rcf for 5 min. Cells were then washed at least 3 times with 1 mL of FACS buffer. After hashtag staining, samples were combined into three groups based on the viral variant of the breakthrough infection (Delta, BA.1/BA.2, or BA.4/BA.5).

Next, samples were enriched for B cells using the human REAlease CD19 MicroBead kit (Miltenyi) according to the manufacturer’s protocol and the B cell fraction was taken for further processing. B cells were then stained with barcoded SARS-CoV-2 S and RBD proteins for 30 min in the dark at 4°C. A mix of probes with two different fluorophores were used to allow detection of double positive SARS-CoV-2 binding B cells and remove any background signal against fluorophores. After staining, cells were washed with FACS buffer. Subsequently, cells were enriched for probe specificity using PE/APC MicroBeads for positive selection (Miltenyi) according to the manufacturer’s protocol. SARS-CoV-2-specific B cells were stained with the TotalSeqC Human Universal Cocktail inside the column for 30 min according to the manufacturer’s protocol. After incubation, cells were washed 3x with MACS buffer (0.5% Bovine serum albumin (BSA) and 2 mM EDTA) and eluted from the column. The positive fraction was bulk sorted on the BD FACS Aria IIU 4 laser sorter. Library preparation was performed using the 10X Genomics platform.

### Spectral flow cytometry analysis

PBMC samples were thawed in separate Falcon tubes containing 40 mL thaw medium by placing them in the water bath at 37°C until they were partially defrosted and later transferred into the pre-warmed thaw medium. Cells were centrifuged at 1600 rpm for 5 min, after which the supernatant was carefully removed by decanting without disturbing the cell pellet. Cell pellets were resuspended in FACS buffer (2% FCS and 1 mM EDTA), adjusted to approximately 2x10^6^ cells/mL and centrifuged at 800 rcf for 5 min. CD3⁺ T cells were depleted from the cell suspension using the CD3⁺ microbeads by Stem Cell Technologies according to the manufacturer’s instruction. Next, SARS-CoV-2 WT spike or Tetanus toxoid protein was loaded onto streptavidin (BB515, BV421, BUV615) in a 2:1 protein to streptavidin ratio and incubated for 1h at 4°C. Next, cells were stained with SARS-CoV-2 WT spike and Tetanus probes for 30 min in the dark at 4°C. After incubation 10 µM biotin was added to occupy any unbound spaces on the streptavidin molecules and prevent false positive signal. Cells were incubated with probes for 30 min in the dark at 4°C and washed with FACS buffer to remove any unbound probe. Finally, cells were stained with the cell surface marker panel as shown in Supplementary table S12 in 100 uL total volume and incubated for 30 min in the dark at 4°C. Cells were washed using FACS buffer and resuspended in a final volume of 150 uL. Cells were measured using the Cytek Aurora spectral flow cytometer.

### Raw data processing

Single-cell RNAseq raw data was processed by the sequencing company (Single Cell Discoveries) using the Cellranger pipeline (10x Genomics). Cells were annotated with the human GRCh38-2020-A reference genome and V(D)J sequences were annotated using the vdj_GRCH38_alts_ensembl-7.1.0 database. Sample demultiplexing was performed using the demux Cellranger pipeline. Data was received as gene expression matrix for transcriptomics and CITE-seq data and a csv file of annotated V(D)J genes for BCR sequencing.

Spectral flow cytometry data was unmixed using the Cytek flow cytometry acquisition software SpectroFlo. Monostain beads or healthy individual PBMCs were used to define marker spectra and used to unmix data without autofluorescence extraction. Data was further compensated using FlowJo v.10 software prior to data normalization and scaling in R.

### Data pre-processing

Cell quality, with regards to gene expression, was assessed by visualizing the distribution of detected features (genes) per cell. Genes that were detected in less than 3 cells were removed. Cells with less than 200 or more than 3500 detected genes were removed, considered as low-quality cells, or doublets, respectively. Cells with a percentage of reads, mapping to mitochondrial genes, greater than 7.5% were also removed, representing highly stressed and/or dead cells. Cells with very low ribosomal protein expression have also been removed, since they were considered low quality. No additional thresholds have been imposed, with regards to surface protein, and probe count data. The data pre-processing was implemented using the Seurat R Package (version 4.4.0)^36^.

Quality control of spectral flow cytometry data was performed using the PeacoQC R package^84^. First, data was transformed using arcsinh transformation with co-factors between 5000 and 8000. The PeacoQC algorithm was run on transformed data with a median absolute diviation of 40 and isolation tree limit 10.

### Data Normalization

The gene expression counts were normalized using the “SCTransform” modelling framework^85,86^. The normalization was implemented with the SCTransform function in Seurat. The “vst.flavor” parameter was set to “v2”, with the rest to the defaults. The cell surface protein counts were normalized with the Centered Log Ratio (CLR) transformation method across cells (“margin” parameter set to 2) using the NormalizeData function in Seurat.

Spectral flow cytometry data was normalized using CytoNorm 2.0^87^. 10,000 cells per sample were randomly sampled to provide a training set for normalization. Normalization was, subsequently, applied to all samples.

### Cell Clustering

The cell clustering was performed using the “Weighted Nearest Neighbor” (WNN) analysis^36^, available through the Seurat R package. In brief, the method provides a framework for the unsupervised learning of the relative contribution of each data modality in every cell, thus enabling the integrative analysis of multimodal data. For more details, we refer to the original publication. In this study, we used three following modalities to perform the WNN clustering: (i) normalized gene expression data, (ii) and normalized cell surface data. The dimensionality reduction of the RNA assay was performed, excluding genes that code for immunoglobulins. The final clusters were identified using the Leiden algorithm, with the resolution value equal to 2. Spectral flow cytometry data was visualized using PacMAP dimensionality reduction. PCAs were established based on previously established markers excluding SARS-CoV-2 spike probes and the novel markers identified by RNAseq (CD99, CD11a, CD1d, CD40, CD22, and BAFF-R). The Leiden algorithm with resolution 0.9 was used to cluster data and annotation was based on cluster marker expression as established by our previous work and scRNAseq results^33^.

### Cluster Marker Identification

The positive cluster markers, for the RNA and surface protein modalities, were identified using the FindAllMarkers Seurat function, as well as the graph_test function from the Monocle3 (Cao et al., 2019) R package (version 1.3.1). The neighbor_graph, and expression_family parameters were set to “knn”, and “negbinomial”, respectively. Seurat uses the Wilcoxon Rank Sum test for marker identification, whereas Monocle3 uses the Moran’s I test. For further details, we refer to the original publications^88,89^. For both methods, we retained markers with adjusted p value lower or equal to 0.05.

### Trajectory Analysis

The lineage inference analysis was performed, on the RNA module, using the Slingshot R package (version 2.10.0)^90^. In brief, Slingshot first constructs a Minimum Spanning Tree (MST) based on the cluster labels to identify the basic shape and the number of the cell lineages. Secondly, the so-called method of “simultaneous principal curves” is applied to generate a smoother depiction of the lineages. Lastly, pseudotime values are acquired by orthogonal projection onto the curves. For more details, we refer to the original publication. In our analysis, the ActB cell cluster or CD73⁺ was used as the starting point of the trajectory for RNAseq and Flow cytometry data, respectively. The rest of the parameters were set to default. Longitudinal data was embedded in pseudotime for analysis of flow cytometry data. The obtained pseudotime values were used for the trajectory visualization.

### Pathway Activity Inference

Pathway activity scores were computed using PROGENy (R package v.2.12.0) with the following parameters: organism=’Human’ top=500, perm=1, scale=FALSE^91^. PROGENy identifies 14 canonical signaling pathways: Androgen, EGFR, Estrogen, Hypoxia, JAK-STAT, MAPK, NFKB, p53, PI3K, TGFβ, Trail, TNFα, VEGF, and WNT For each cell, the method calculates pathway activity scores based on the weighted expression of pathway-responsive genes determined from perturbation experiments. Pathway activity scores were scaled (zscore normalization) for cross-sample and cross-pathway comparison.

### Cell Specificity Definition based on scRNAseq probes

The cell specificity thresholds for the different SARS-CoV-2 proteins were set after manual assessment of the scatterplots of the raw probe count data. Explicitly, specificity was defined as follows: (i) cells with 4 or more probe counts against SARS-CoV-2 WT were considered WT-specific; (ii) cells with 2 or more probe counts against SARS-CoV-2 Delta spike were considered Delta-specific; (iii) cells with 5 or more probe counts against SARS-CoV-2 Omicron BA.1 RBD were considered Omicron BA.1-specific; (iv) cells with 3 or more probe counts against SARS-CoV-2 Omicron BA.2 RBD were considered Omicron BA.2-specific; (v) cells with 4 or more probe counts against SARS-CoV-2 Omicron BA.4/BA.5 spike were considered Omicron BA.4/BA.5-specific. The specificity of the cells was also validated to match the breakthrough infection types of the donor, each particular cell was derived from^91^.

### Cluster correlation analysis for B cell stage definition

Due to large inter-individual differences in activation pathways and speed, B cell stages of activation were defined based on cluster frequency of samples overcoming individual and time-based bias. Cluster Frequency of SARS-CoV-2 spike specific B cells was calculated per sample. Correlations between samples based on cluster frequency was calculated using Pearson correlations. Next, distance between samples was calculated based on correlation with a complete negative correlation of -1 defined as a distance of 0, and a complete positive correlation as the biggest distance. Hierarchical clustering based on the average linkage algorithm was used to cluster samples together based on their distance. Stages were annotated based on cluster abundance and timepoint of enrichment.

### Production of recombinant monoclonal antibodies

Sequences of the HC and LC of SARS-CoV-2 specific B cells were obtained from the single-cell V(D)J enriched library. Plasmids encoding the selected sequences were ordered from GenScript gene service. Suspension HEK293F cells (Invitrogen) were maintained in FreeStyle medium (Gibco) and co-transfected with a 1:3 ratio of the two HC/LC DNA plasmids and PEI-max (1 µg/mL). Six days after transfection, the supernatants were centrifuged at 4000 rpm for 30 min, followed by filtration through 0.22 µm SteriTop filters (Millipore). The filtered supernatants containing the produced recombinant mAbs were incubated overnight with protein G beads and run the next day over a 10 mL column (Pierce). The mAbs were eluted with elution buffer (0.1 M glycine pH 2.5) into neutralization buffer (1 M Tris pH 8.7), concentrated and buffer exchanged to PBS using 50 kDa VivaSpin20 columns (Sartorius). The concentration of the mAbs was determined by the Nanodrop.

### Luminex binding assay

SARS-CoV-2 spike proteins were coupled to Magplex beads (Luminex Corporation) in a 75:12.5 ratio of spike protein to million beads. Equimolar amounts compared to the spike proteins were used to couple the other proteins included in the panel. To avoid aspecific binding, beads coupled to the proteins were blocked with a 30 min incubation step with phosphate-buffered saline (PBS) that contained 2% bovine serum albumin (BSA), 3% fetal calf serum (FCS) and 0.02% Tween-20 at pH 7.0. Subsequently, coupled beads were stored in the dark at 4°C in PBS with 0.05% sodium azide and employed within one year after coupling.

To measure the binding of mAbs to the different proteins coupled to the beads, 50 μL of a bead mixture in a concentration of 20 beads per μL were added to 50 μL of diluted mAbs. Following an incubation O/N at 4°C on a rotator, plates were washed twice with Tris-buffered saline (TBS) containing 0.05% Tween-20 (TBST) and resuspended with 50 μL (1.3 μg/mL) of goat-anti-human IgG-PE (Southern Biotech). Next, plates were incubated for 2 h on a plate shaker at room temperature and then washed twice with TBST before the addition of 70 μL of Magpix drive fluid (Luminex). Read-out was performed on a Magpix (Luminex). Antibody binding was expressed as the median fluorescence intensity (MFI) of approximately 50 to 150 beads per well and corrected for background signals by subtracting the MFI of wells containing only buffer and beads.

### Pseudovirus design and neutralization assay

The WT D614G, Delta, Omicron BA.1, BA.2, BA.4/5, BA.2.75, BQ.1.1, XBB.1.5, and EG.5 pseudovirus constructs were ordered as gBlock gene fragments (Integrated DNA Technologies) and cloned using an optimized Gibson Assembly protocol, as described before^92^. All constructs were produced by co-transfecting HEK293F cells with a plasmid encoding the SARS-CoV-2 spike with the pHIV-1NL43-ΔEnv-NanoLuc reporter virus plasmid. Cell supernatants containing the pseudoviruses were harvested after 48 h from transfection and stored in the -80°C freezer. To test the neutralizing activity of the produced mAbs, these were serially diluted in 3-fold steps starting at a concentration of 10 µg/mL (which was also defined as our cut-off value). They were then mixed with the pseudoviruses and incubated for 1 h at 37°C. The mAb/virus mixes were then added to HEK293-ACE2 cells, which were seeded the day before in poly-L-lysine-coated 96well plates at a density of 20.000 cells/well. After 48 h, the mAb/virus mixes were removed, cells were lysed and transferred to half-area 96-well microplates (Greiner Bio-One). Luciferase activity of cell lysate was measured using the NanoGlo luciferase assay System (Promega) with a Glomax late reader (Turner BioSystems).

### Biolayer interferometry (BLI) assay

To test the competition of the most potent neutralizing mAbs with the ACE2 receptor, we employed Ni-NTA sensors (Sartorius) which were loaded with 10 μg/mL of His-tagged SARS-CoV-2 WT RBD for 300 s, followed by a first association step with 10 μg/mL of the produced mAb for 300 s and a second association step of 10 μg/mL of ACE2 receptor for 300 s. Sensors were regenerated between measurements with a solution of 10 mM glycine in PBS. All BLI experiments were performed using an Octet K2 instrument (ForteBio).

### Data Visualization and manipulation

The single-cell RNAseq and flow cytometry data have been visualized and manipulated using the tidyverse collection of R packages (version 2.0.0)^93^. Data regarding mAbs have been investigated using GraphPad Prism 9.

## List of supplementary materials

- Supplementary Figure S1. QC analysis of multi-omics scRNAseq experiment.
- Supplementary Figure S2. SARS-CoV-2 probe labeling strategies for simultaneous FACS and scRNAseq antigen-specificity analysis.
- Supplementary Figure S3. Total B cell and donor composition analysis.
- Supplementary Figure S4. Characterization of distinct B cell populations through multimodal data analysis.
- Supplementary Figure S5. Gene expression UMAPs of AtypB markers.
- Supplementary Figure S6. UMAPs of cell surface markers expressed by AtypB.
- Supplementary Figure S7. Cluster composition based on specificity and CD45RB expression per cluster.
- Supplementary Figure S8. Competition assay of expressed mAbs.
- Supplementary Figure S9. Cell surface marker expression in the IgG⁺ memory composite dataset.
- Supplementary Figure S10. Histograms of cell surface marker expression in the IgG⁺ memory composite dataset.
- Supplementary Figure S11. PACMAP of SARS-CoV-2 IgG⁺ MBCs per sample.
- Supplementary Figure S12. Cluster frequency analysis per phenotypic stage corrected for total B cells.
- Supplementary Figure S13. Phenotypic stage analysis of unspecific B cells.
- Supplementary Figure S14. Cell surface marker expression in the IgG⁺ memory composite dataset including ASCs.
- Table S1. Participant characteristics from the COSCA study
- Table S2. Participant characteristics from the ReCoVERED study
- Table S3. 1 vs all cluster comparison Wilcoxon test ADT markers
- Table S4. 1 vs all cluster comparison Wilcoxon test unsupervised gene analysis
- Table S5. 1 vs all cluster comparison Wilcoxon test supervised gene analysis
- Table S6. 1 vs all cluster comparison Wilcoxon test PROGENy pathway analysis
- Table S7. Pairwise Wilcoxon Test of SHM levels between B cell subsets
- Table S8. Linear mixed effects model of SARS-CoV-2 spike⁺ IgG⁺ B cells
- Table S9. Pairwise Wilcoxon Test of cell surface marker expression per B cell subset
- Table S10. Mixed effects model of phenotypic stages across timepoints
- Table S11. Pairwise Wilcoxon Test of B cell subset composition per phenotypic stage
- Table S12. Spectral Flow Cytometry phenotypic panel
- Supplemental Materials and Methods

## Acknowledgments

We thank Dirk Eggink from RIVM, the Netherlands, for providing the plasmids encoding the SARS-CoV-2 Omicron BA.4/5, BA.2.75 and BQ.1.1 S proteins, and for patient recruitment in the COSCA cohort. We also thank all participants of the COSCA and RECoVERED cohorts for their participation and the researchers involved in patient recruitment and sample collection.

## Author contributions

Conceptualization: L.T.M.G., D.G., A.C., T.B., M.J.v.G., M.C.

Methodology: L.T.M.G, D.G., A.C., M.J.v.G., M.C.

Investigation: L.T.M.G, D.G., A.C., G.K., M.C.

Resources: L.T.M.G, D.G., K.vdS., M.P, M.D.dJ, G.J.dB., R.W.S., M.J.v.G., M.C.

Data curation and visualization: L.T.M.G., D.G., A.C., M.C.

Supervision: T.B., M.J.v.G., M.C.

Funding acquisition: R.W.S., M.J.v.G., M.C.

Writing-Original draft: L.T.M.G., D.G., A.C., M.J.v.G., M.C.

Writing-Review and editing: all authors.

## Data and materials availability

scRNAseq data will become available on the European Nucleotide Archive upon publication.

