## Supplementary data for "A dynamic activation–memory cycle organizes the human IgG⁺ memory B cell compartment"

### Supplementary figures and tables

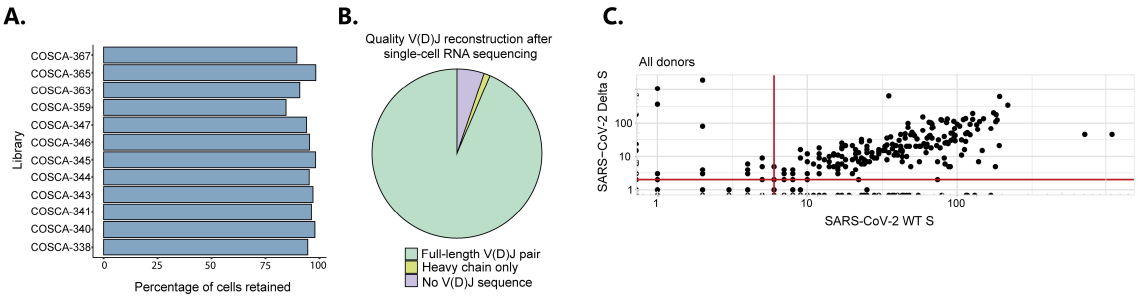


**Supplementary Figure S1**. **QC analysis of multi-omics scRNAseq experiment.**

Plots showing QC analysis of the obtained single-cell libraries. **(A)** Percentage of retained B cells divided per donor. **(B)** Pie chart showing the quality of V(D)J reconstruction after single-cell RNA sequencing. **(C)** Scatterplot showing an example of threshold definition for antigen specificity analysis. Threshold values for each SARS-CoV-2 protein are reported in the Method section.

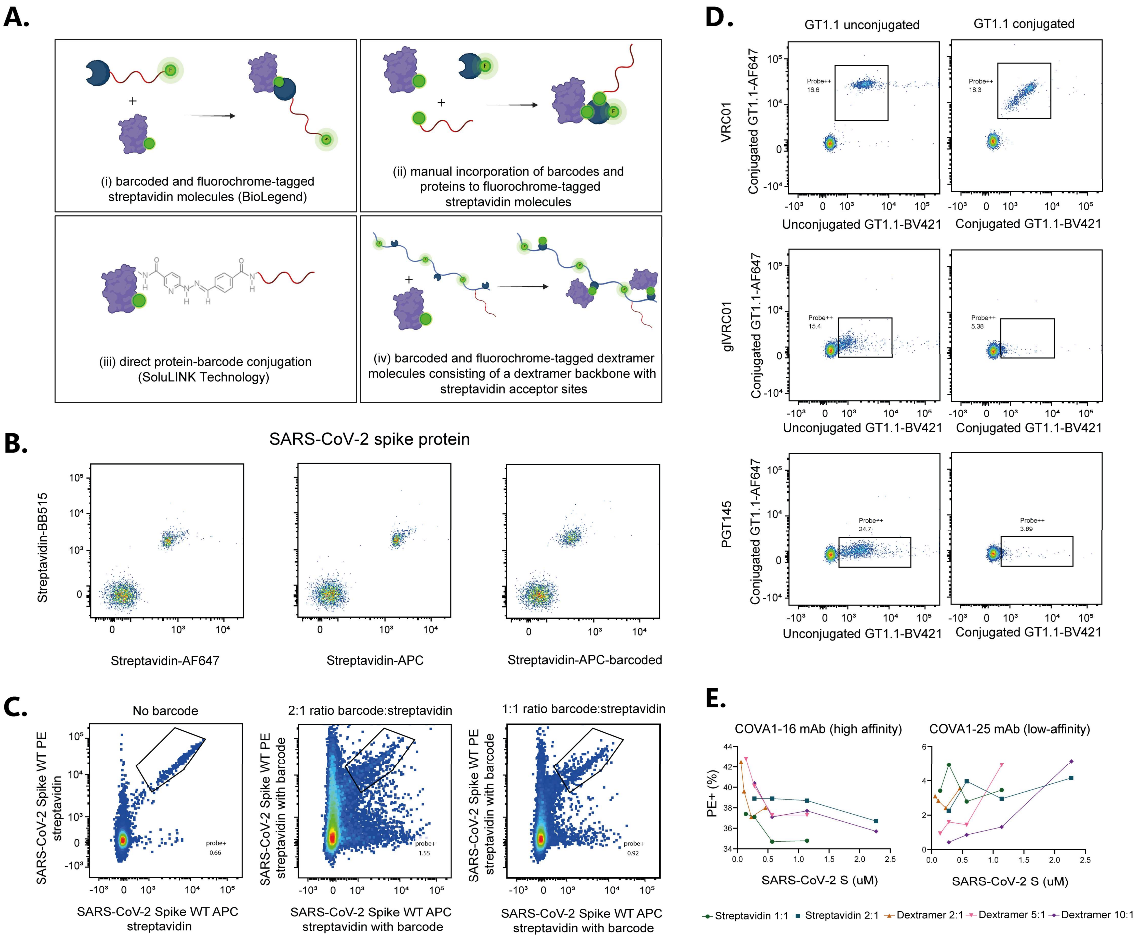


**Supplementary Figure S2. SARS-CoV-2 probe labeling strategies for simultaneous FACS and scRNAseq antigen-specificity analysis.**

Diverse methods to label SARS-CoV-2 proteins. **(A)** Schematic representation of the four different labeling methods tested for SARS-CoV-2 protein labeling for flow cytometry and single-cell RNA sequencing detection. Created in BioRender. Van Gils, M. (2026) https://BioRender.com/jht8f0d. **(B)** Use of streptavidin molecules with (APC barcode) or without (APC or AF647) oligonucleotide barcode for single-cell RNA sequencing (BioLegend). **(C)** Frequency of SARS-CoV-2-specific CD19$^{+}$ B cells stained with manually barcoded SARS-CoV-2 WT S streptavidin constructs with either a 2:1 or 1:1 oligonucleotide. **(D)** Bead assay with directly labelled single-stranded oligonucleotide barcodes to an HIV-1 envelope protein. Binding of antibodies known to target different epitopes on the HIV-1 envelope protein with low and high affinity is shown. **(E)** Bead assay of comparison of SARS-CoV-2 dextramer and spike probes in different protein to construct ratio. The percentage of SARS-CoV-2 probe positive beads for different protein concentrations is shown for two known mAbs with low (COVA1-25) and high (COVA1-16) affinity.

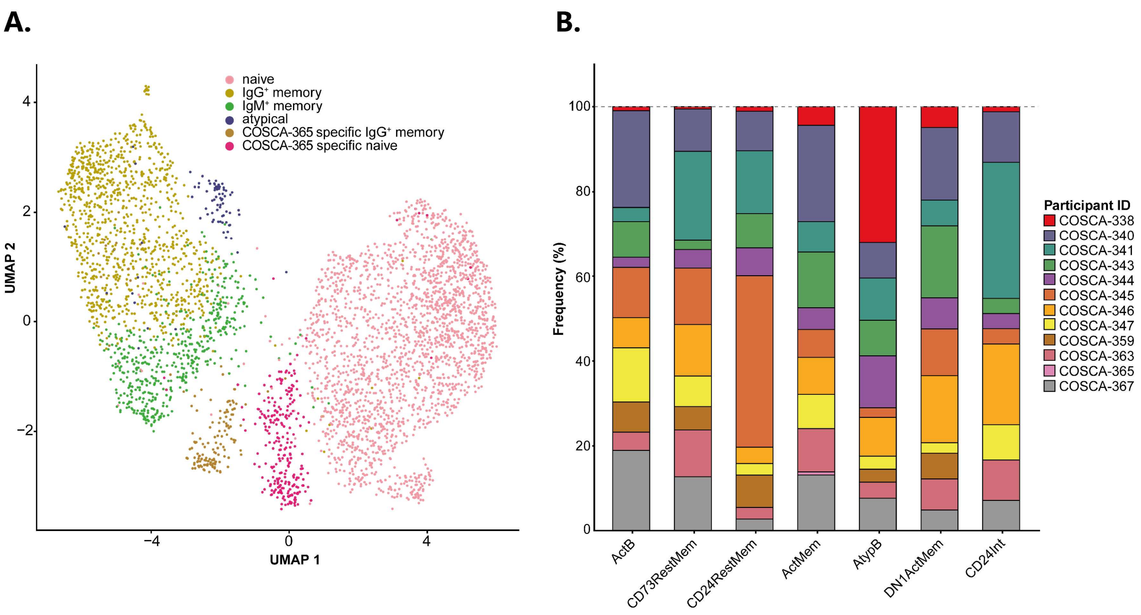


**Supplementary Figure S3. Total B cell and donor composition analysis.**

**(A)** UMAP showing the entire B cell population divided into naive, memory and atypical b cell clusters. B cells from the COSCA-365 donor are shown as separate participant-specific clusters. The annotation was conducted based on known cell surface protein markers. **(B)** Stacked bar plot of participant frequency within every B cell subset.

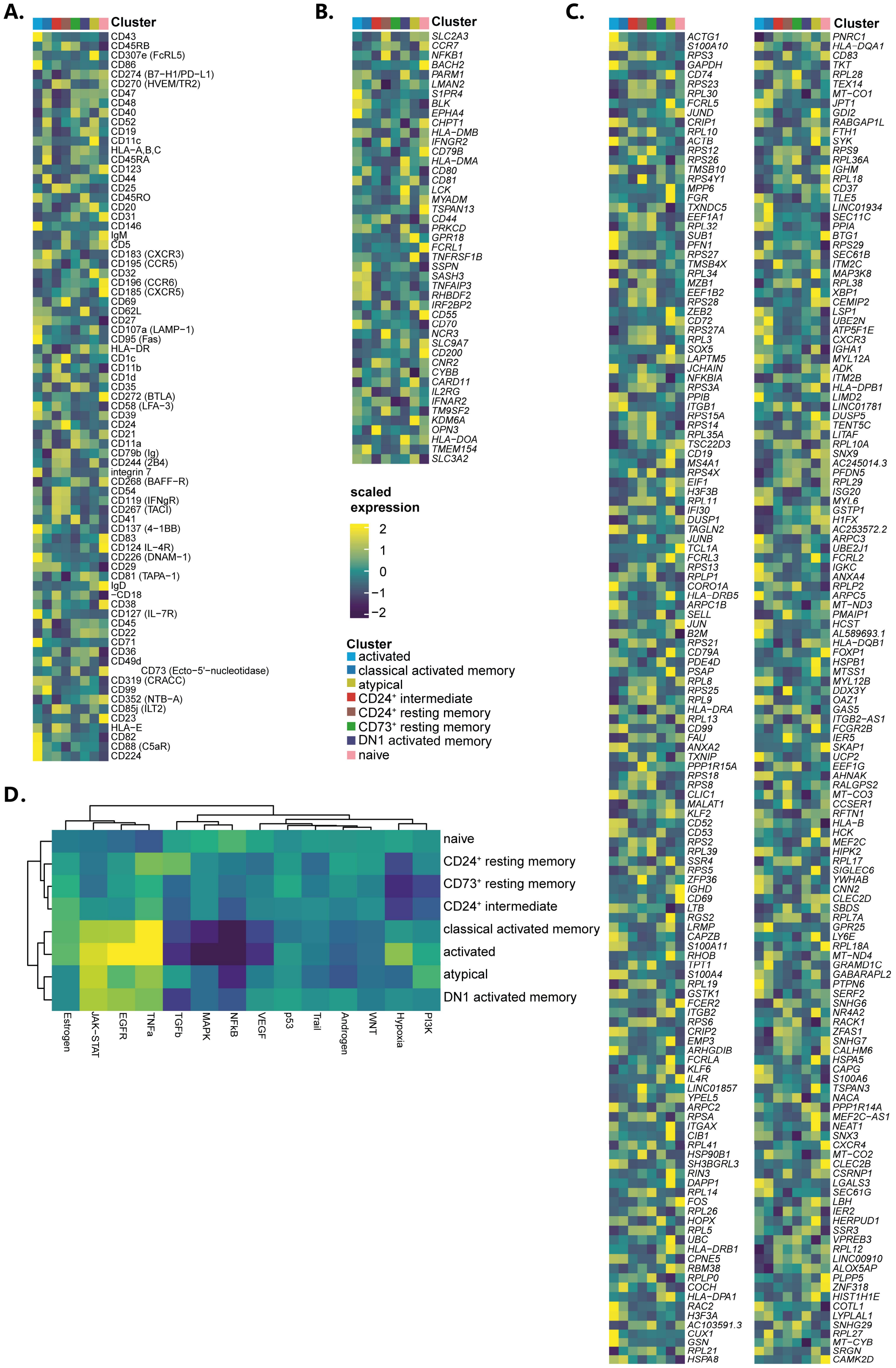


**Supplementary Figure S4. Characterization of distinct B cell populations through multimodal data analysis.**

**(A)** Surface protein ADT expression per MBC cluster. **(B)** mRNA markers per MBC cluster as defined supervised analysis. The markers were identified based on biological relevance and verified through spatial autocorrelation analysis using Moran's I test. **(C)** mRNA markers per MBC cluster as defined by unsupervised analysis. The markers were identified through spatial autocorrelation analysis using Moran's I test. **(D)** Pathway activity analysis per MBC cluster as defined by PROGENy[91].

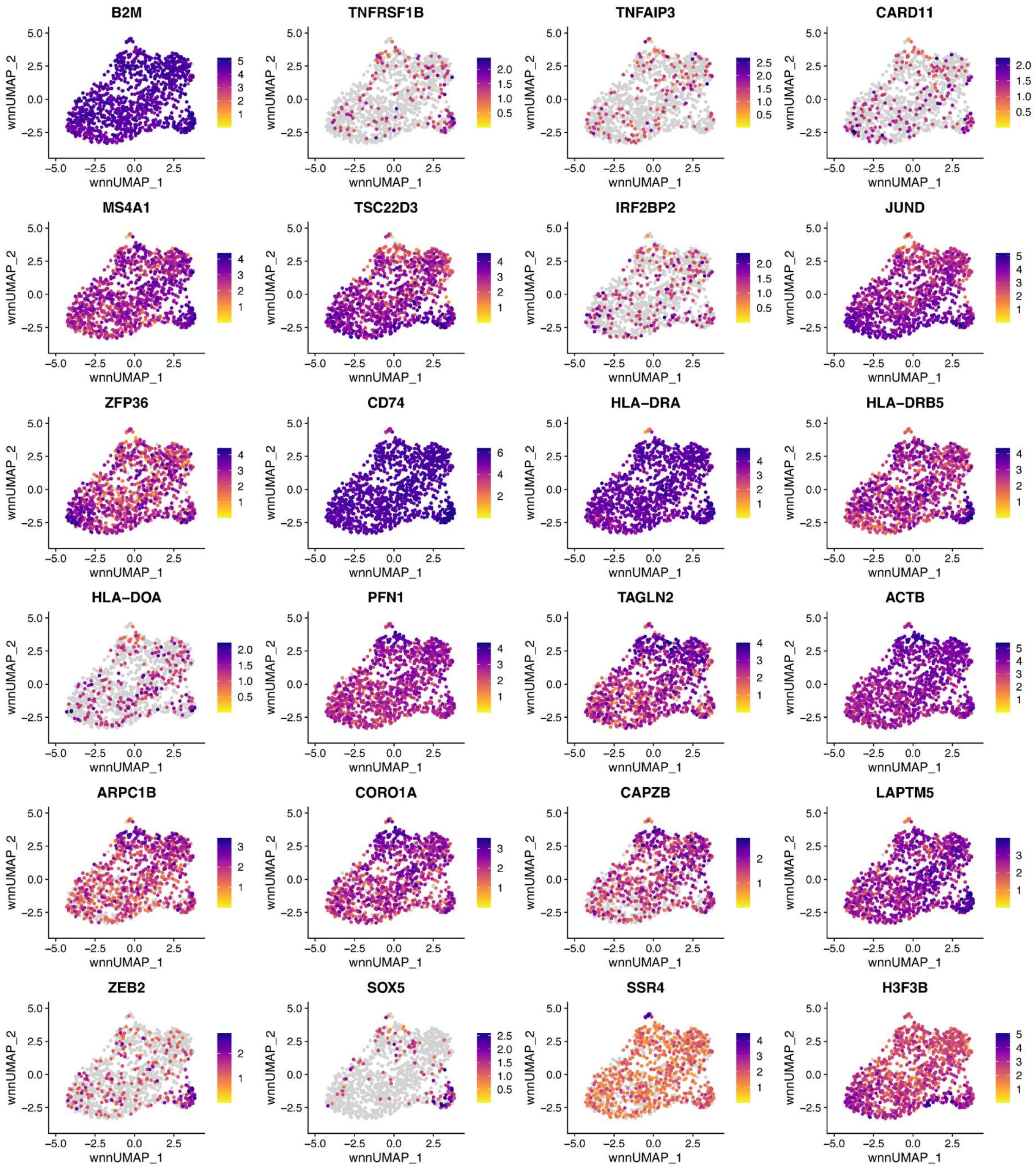


**Supplementary Figure S5. Gene expression UMAPs of AtypB markers.**

UMAPs showing gene expression levels of genes that were differentially expressed in the AtypB compartment as described in the main text.

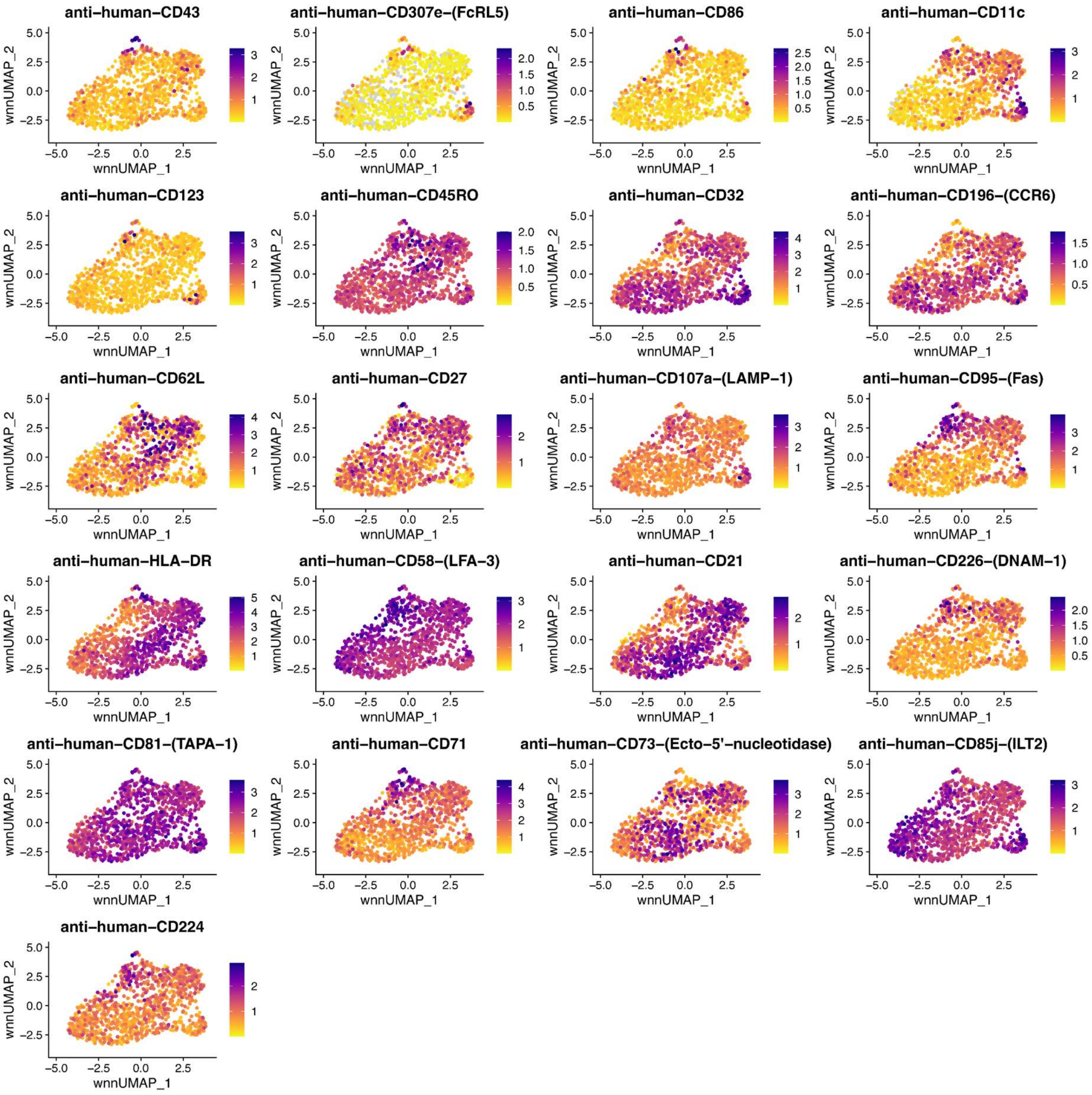


**Supplementary Figure S6. UMAPs of cell surface markers expressed by AtypB.**

UMAPs showing cell surface marker expression levels of proteins described in the main text that were differentially expressed in the AtypB compartment as detected by CITE-seq analysis.

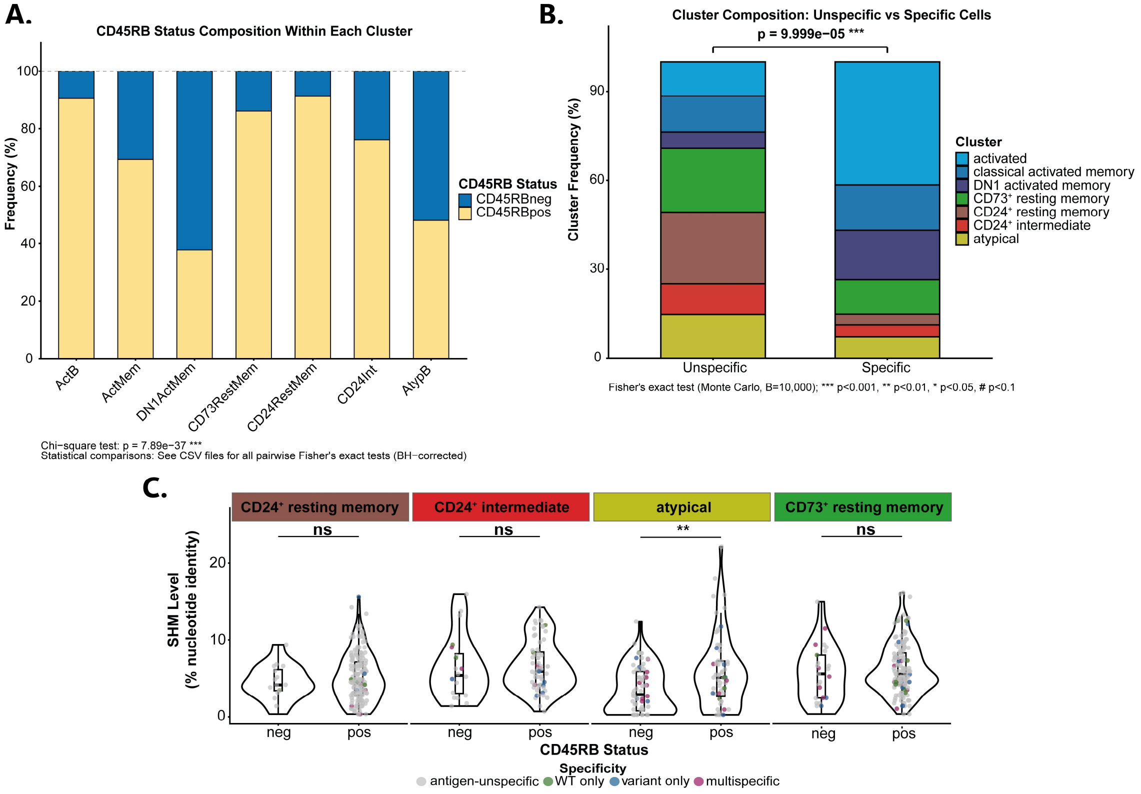


**Supplementary Figure S7. Cluster composition based on specificity and CD45RB expression per cluster.**

**(A)** Barplot of CD45RB expression per B cell subset. Cells were labeled as CD45RB$^-$ or CD45RB$^+$ based on a set threshold of CD45RB expression. The frequency of CD45RB$^+$ and CD45RB$^-$ B cells per subsets were depicted as frequency of total B cells within the B cell subset. **(B)** Barplot showing cluster composition of unspecific and SARS-CoV-2 specific B cells. **(C)** SHM frequency analysis within the CD24$^+$ RestMem, CD24$^+$ Int, AtypB, and CD83$^+$ RestMem subsets. Cells were separated based on their CD45RB status to allow comparison between CD45RB$^-$ and CD45RB$^+$ B cells. Pairwise comparison can be found in Table S7.

A.

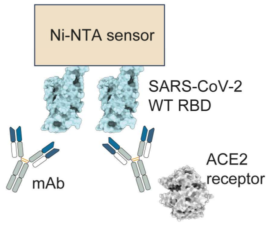


B.

| | | % residual ACE2 binding to WT RBD |
|---|---|---|
| activated | COVA340-02 | 1 |
| | COVA341-03 | 39 |
| | COVA363-01 | 2 |
| classical activated memory | COVA340-01 | 48 |
| | COVA340-05 | 45 |
| | COVA340-06 | 10 |
| | COVA347-01 | 55 |
| | COVA359-01 | 42 |
| | COVA2-15 | 2 |
| | mock | 100 |

**Supplementary Figure S8. Competition assay of expressed mAbs.**

**(A)** Schematic representation of the competition assay between the studied mAbs and the ACE2 receptor for binding to the SARS-CoV-2 WT RBD protein. **(B)** Values corresponding to the percentage of residual binding of the receptor in the presence of the competitor mAb, as assessed by biolayer interferometry (bottom panel). mAb COVA2-15 and no competitor/mock were included as positive and negative controls, respectively.

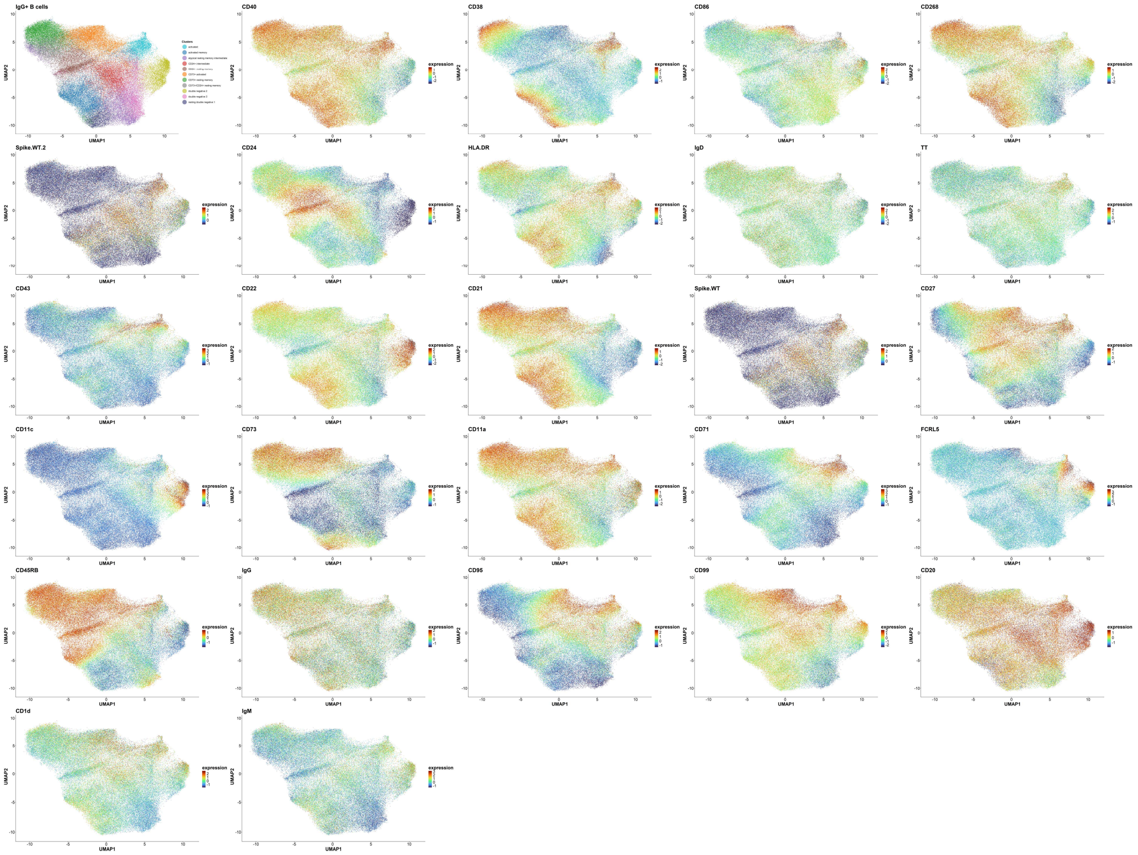


**Supplementary Figure S9**. **Cell surface marker expression in the IgG$^{+}$ memory composite dataset**.

PACMAPs of IgG$^{+}$ MBC displaying scaled cell surface marker expression of all markers included in the spectral flow cytometry B cell panel. Dimensionality reduction was performed with PACMAP and clustering was done on the PCAs using the Leiden algorithm. Cluster annotation was done based on expression of cell surface markers.

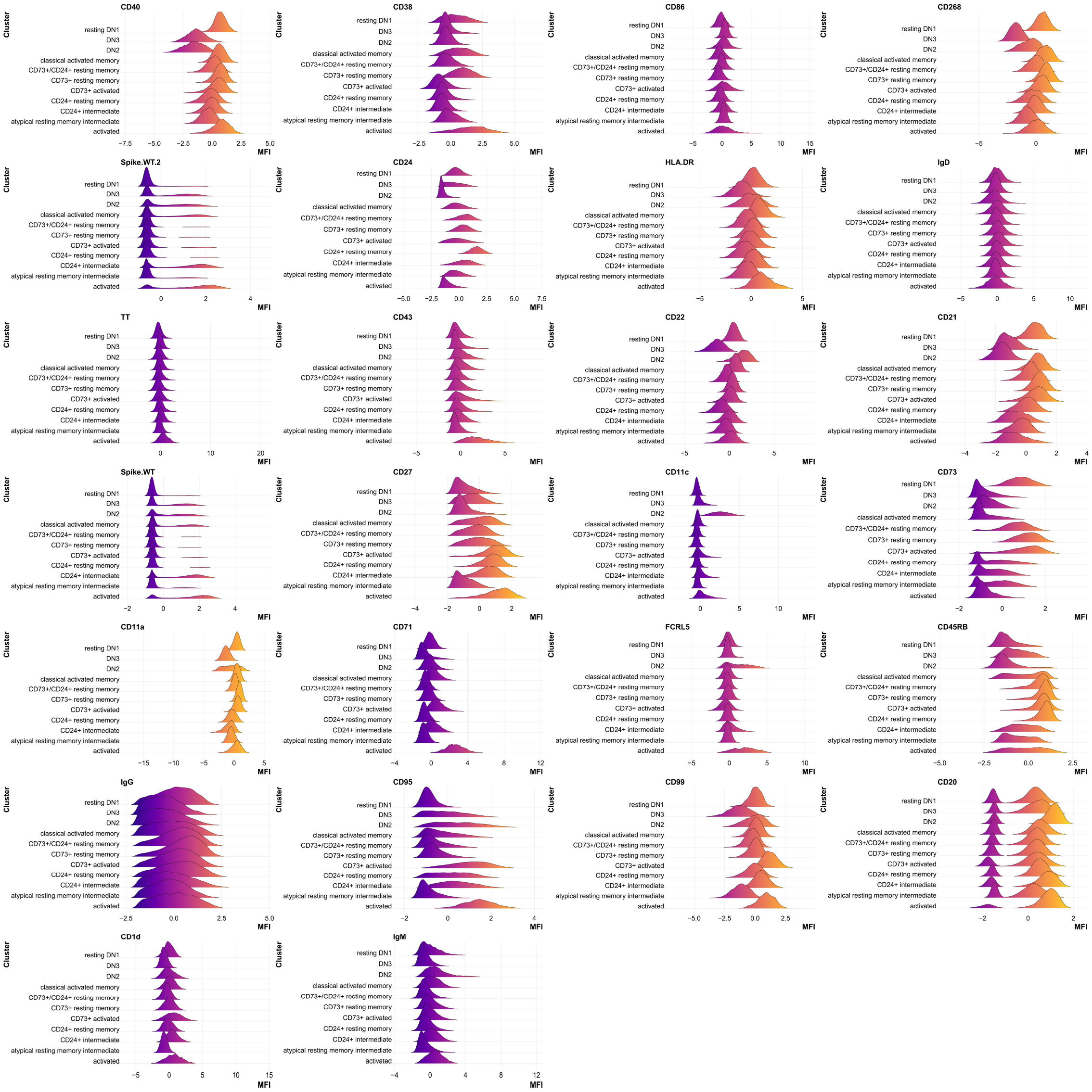


**Supplementary Figure S10. Histograms of cell surface marker expression in the IgG⁺ memory composite dataset.**

Histograms of MFI expression per cell surface marker incorporated in the flow cytometry phenotyping. MFI expression is shown per B cell subset as defined by dimensionality reduction and Leiden clustering performed on the IgG⁺ MBC compartment.

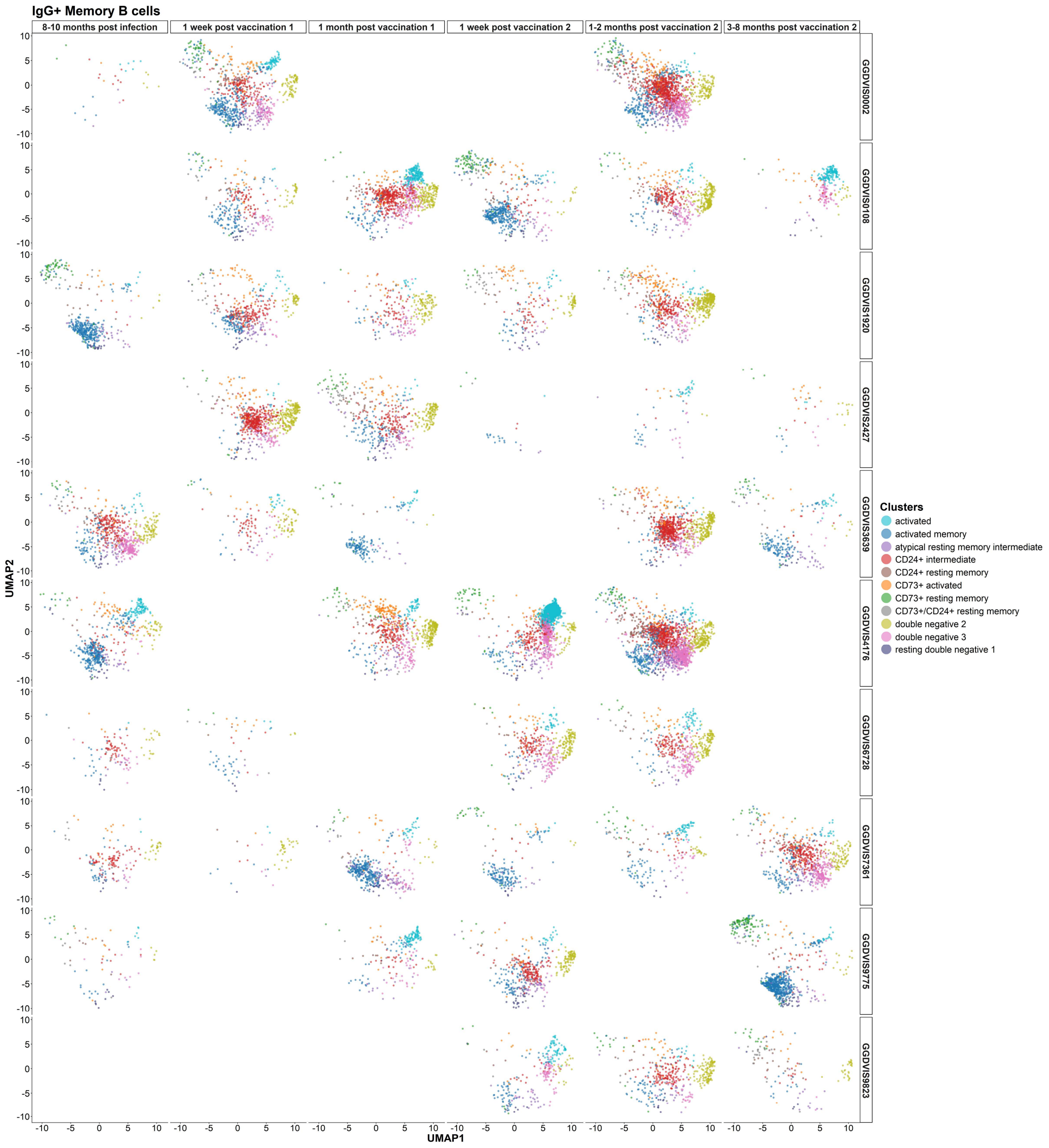


**Supplementary Figure S11**. **PACMAP of SARS-CoV-2 IgG⁺ MBCs per sample.**

PACMAPs of SARS-CoV-2 IgG⁺ MBCs divided per individual and timepoint. Dimensionality reduction was performed with PACMAP and clustering was done on the PCAs using the Leiden algorithm. Cluster annotation was done based on expression of cell surface markers.

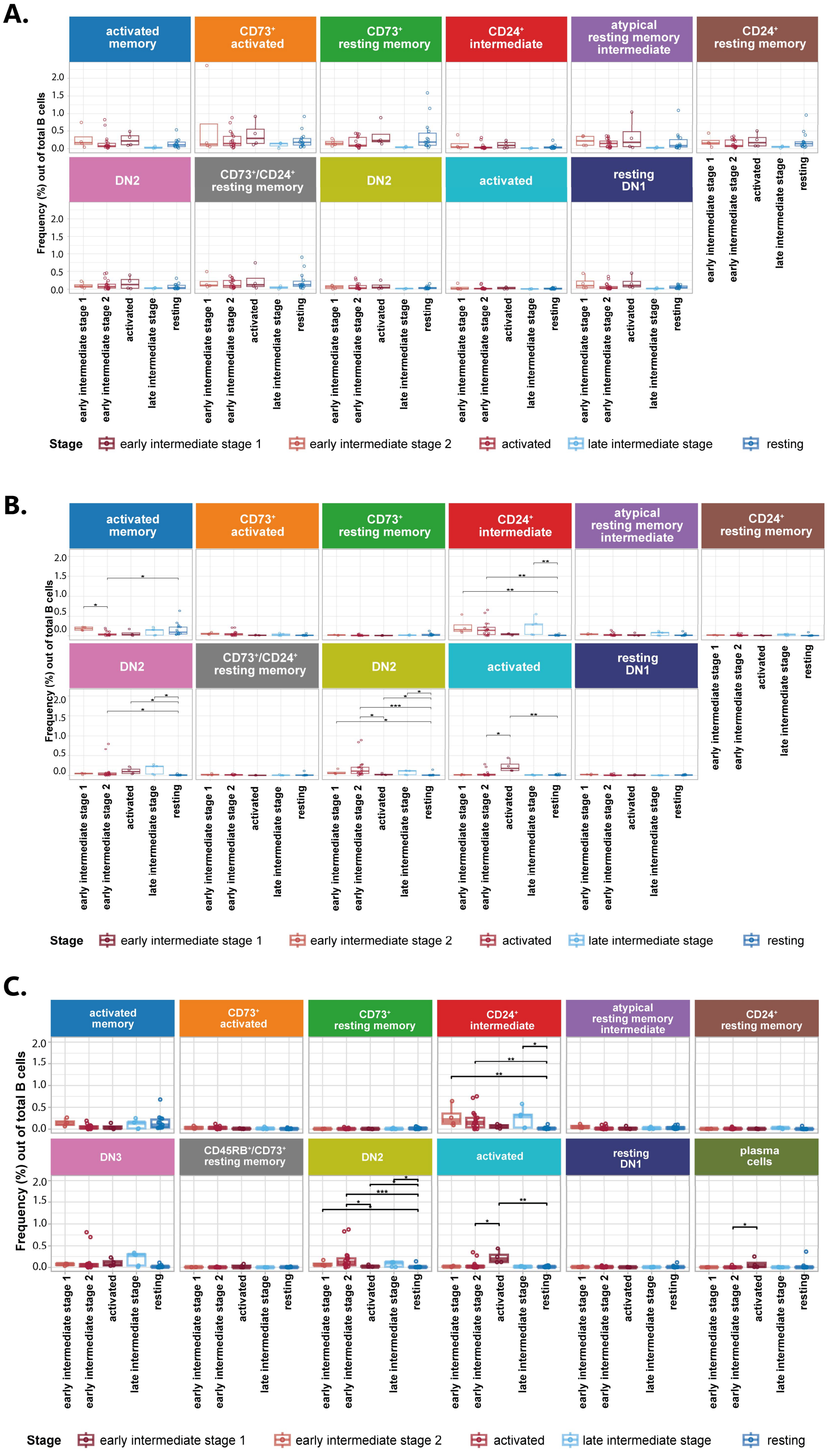


**Supplementary Figure S12**. **Cluster frequency analysis per phenotypic stage corrected for total B cells**

**(A)** Frequency of B cell clusters found per stage in the unspecific B cells included in the IgG⁺ MBC composite dataset. **(B)** Frequency of B cell clusters found per stage in the SARS-CoV-2 specific B cells included in the IgG⁺ MBC composite dataset. **(C)** Frequency of B cell clusters found per stage in the SARS-CoV-2 specific B cells included in the IgG⁺ MBC composite dataset including ASCs. A pairwise Wilcoxon test was performed to determine significant differences in frequency between stages. P-values were adjusted for multiple testing using the Benjamini-Hochberg method. Significance was indicated as * <0.05, **<0.01, ***<0.001, **** <0.0001.

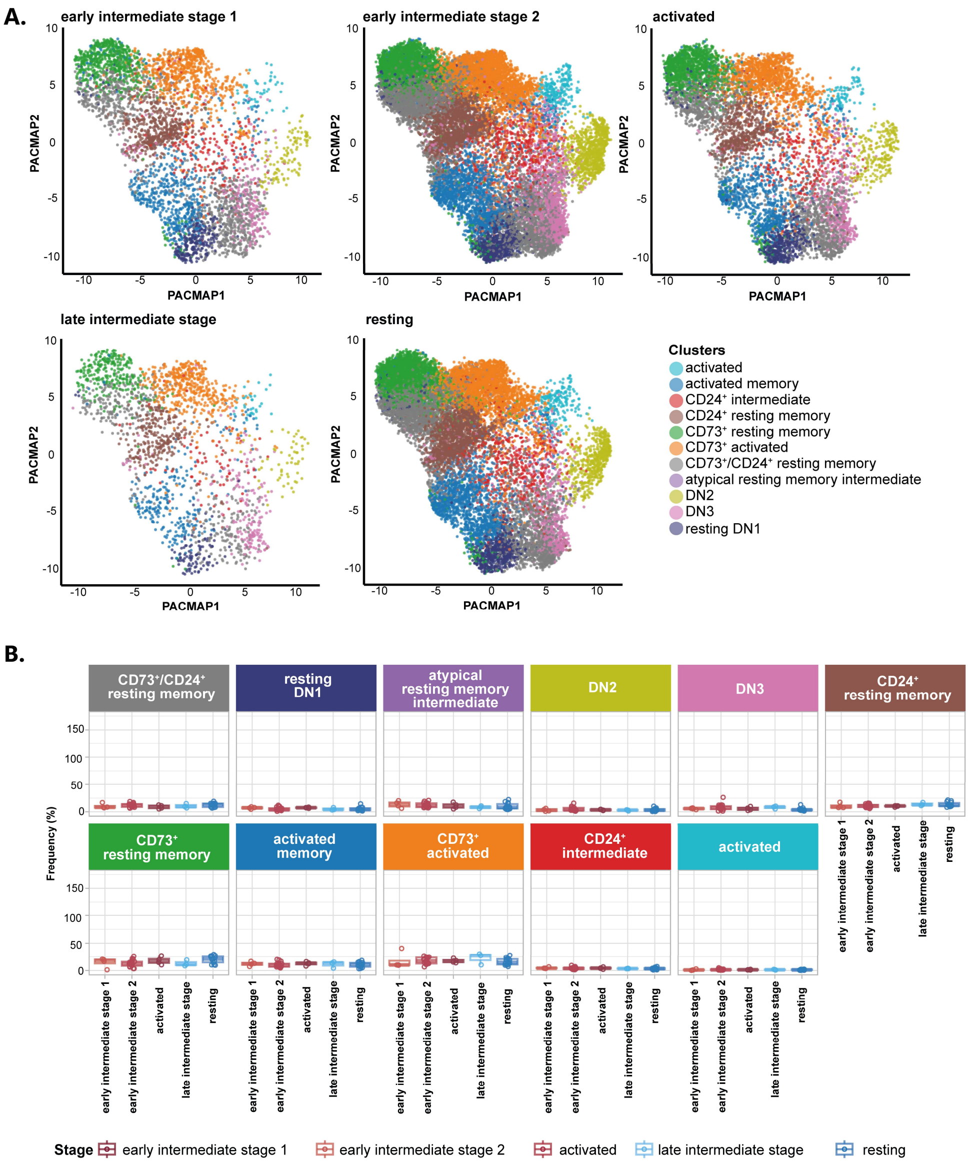


**Supplementary Figure S13**. **Phenotypic stage analysis of unspecific B cells.**

**(A)** PACMAP of unspecific IgG$^+$ MBCs per phenotypic stage. Dimensionality reduction was performed with PACMAP and clustering was done on the PCAs using the Leiden algorithm. Cluster annotation was done based on expression of cell surface markers. The 5 phenotypic stages were previously determined using the SARS-CoV-2 specific IgG$^+$ MBCs from the composite dataset and assigned to the unspecific B cells based on their sample origin. **(B)** Frequency of B cell clusters found per stage in the unspecific IgG$^+$ MBCs. A pairwise Wilcoxon test was performed to determine significant differences in frequency between stages. P-values were adjusted for multiple testing using the Benjamini-Hochberg method. Significance was indicated as * <0.05, **<0.01, ***<0.001, **** <0.0001.

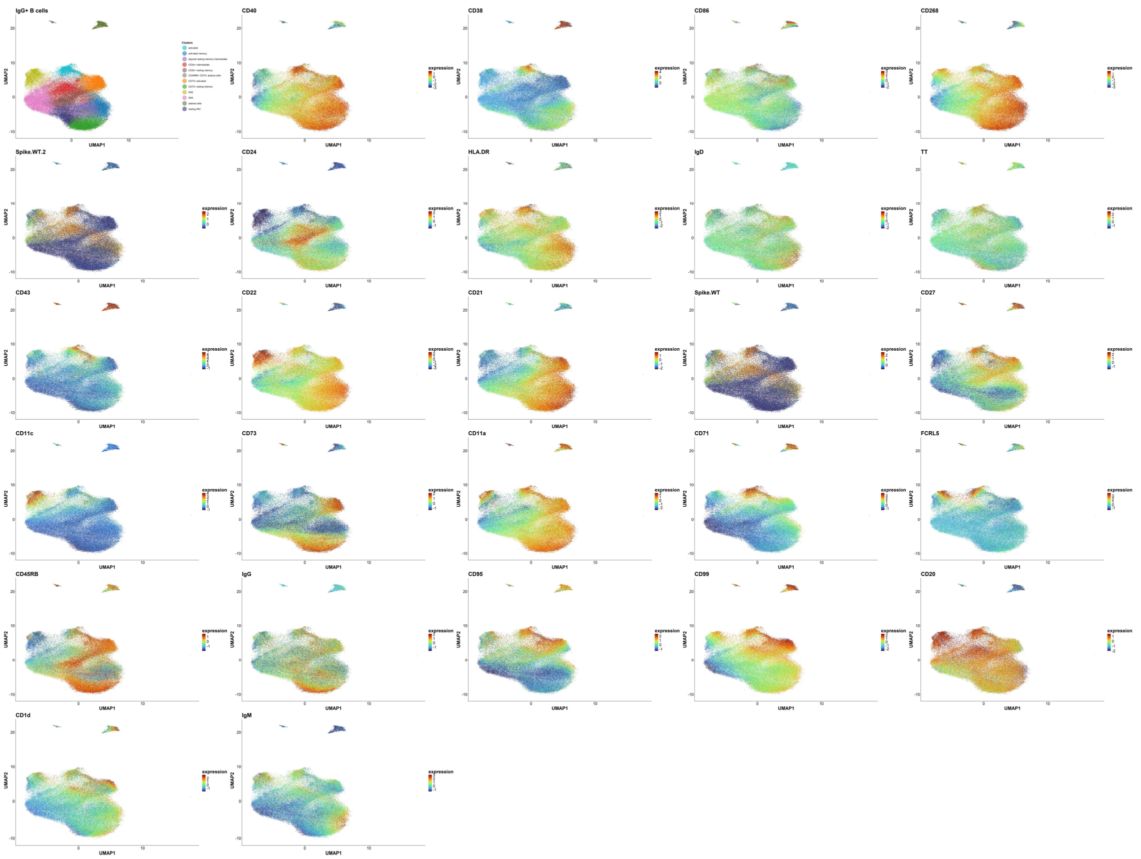


**Supplementary Figure S14**. **Cell surface marker expression in the IgG$^+$ memory composite dataset including ASCs.** PACMAPs of IgG$^+$ MBCs including ASCs displaying scaled cell surface marker expression of all markers included in the spectral flow cytometry B cell panel. Dimensionality reduction was performed with PACMAP and clustering was done on the PCAs using the Leiden algorithm. Cluster annotation was done based on expression of cell surface markers.

#### Table S1. Participant characteristics from the COSCA study

| | COSCA-338 | COSCA-340 | COSCA-341 | COSCA-343 | COSCA-344 | COSCA-345 | COSCA-346 | COSCA-347 | COSCA-359 | COSCA-363 | COSCA-365 | COSCA-367 |
|---|---|---|---|---|---|---|---|---|---|---|---|---|
| **Age** | 23 | 24 | 32 | 59 | 28 | 25 | 27 | 29 | 25 | 51 | 62 | 46 |
| **Sex** | Female | Female | Female | Male | Female | Male | Female | Female | Male | Female | Female | Female |
| **Hospital admission** | No | No | No | No | No | No | No | No | No | No | No | No |
| **Fever** | Yes | Yes | No | No | Yes | Yes | Yes | No | No | Yes | Yes | Yes |
| **Fever duration (days)** | 4 | 1 | - | - | 4 | 3 | 2 | - | - | 5 | 3 | 4 |
| **PCR confirmed infection** | Yes | Yes | Yes | Yes | Yes | Yes | No | Yes | No | Yes | Yes | Yes |
| **Strain causing infection** | Delta | Delta | Delta | Delta | Delta | Delta | Omicron BA.2 | Omicron BA.1 | Omicron BA.2 | Omicron BA5.1 | Omicron BA.4 | Omicron BA.5 |
| **Sequence confirmed strain** | Likely | Likely | Likely | Likely | Likely | Likely | Likely | Likely | Likely | PCR proven | PCR proven | PCR proven |
| **Previous infection** | No | No | No | No | No | No | Yes | No | No | No | No | No |
| **1st vaccination** | Moderna | Moderna | Pfizer | Pfizer | Pfizer | Moderna | Moderna | Moderna | Moderna | Moderna | Astra Zeneca | Pfizer |
| **Date 1st vaccination** | 01-06-2021 | 31-05-2021 | 09-02-2021 | 07-05-2021 | 27-06-2021 | 15-06-2021 | 17-05-2021 | 21-05-2021 | 07-07-2021 | 14-04-2021 | 14-04-2021 | 12-06-2021 |
| **2nd vaccination** | Moderna | Moderna | Pfizer | Pfizer | Pfizer | Moderna | Moderna | Moderna | Moderna | Moderna | Astra Zeneca | Pfizer |
| **Date 2nd vaccination** | 28-06-2021 | 29-06-2021 | 17-03-2021 | 11-06-2021 | 01-08-2021 | 20-07-2021 | 17-06-2021 | 22-06-2021 | 11-08-2021 | 12-05-2021 | 23-06-2021 | 20-07-2021 |
| **3rd vaccination** | - | - | - | - | - | - | Pfizer | Pfizer | Pfizer | Pfizer | Moderna | Pfizer |
| **Date 3rd vaccination** | - | - | - | - | - | - | 08-12-2021 | 08-12-2021 | 11-01-2022 | 29-11-2021 | 20-12-2021 | 29-01-2022 |
| **Start symptoms post infection** | 16-11-2021 | 17-11-2021 | 21-11-2021 | 19-11-2021 | 20-11-2021 | 15-11-2021 | 01-03-2022 | 05-02-2022 | 08-05-2022 | 13-06-2022 | 19-06-2022 | 21-06-2022 |
| **Date blood withdrawal** | 13-12-2021 | 21-12-2021 | 11-01-2022 | 23-12-2021 | 23-12-2021 | 23-12-2021 | 29-03-2022 | 15-03-2022 | 14-06-2022 | 12-07-2022 | 28-07-2022 | 27-07-2022 |

#### Table S2. Participant characteristics from the ReCoVERED study

| | ReCoVered-GGDVIS0002 | ReCoVered-GGDVIS0108 | ReCoVered-GGDVIS1920 | ReCoVered-GGDVIS2427 | ReCoVered-GGDVIS3639 | ReCoVered-GGDVIS4176 | ReCoVered-GGDVIS6728 | ReCoVered-GGDVIS7361 | ReCoVered-GGDVIS9775 | ReCoVered-GGDVIS9823 |
|---|---|---|---|---|---|---|---|---|---|---|
| **Age** | 39 | 27 | 34 | 28 | 62 | 61 | 64 | 42 | 32 | 79 |
| **Sex** | Male | Female | Male | Male | Female | Male | Male | Female | Female | Male |
| **Hospital admission** | No | No | No | No | No | No | No | No | No | No |
| **1st vaccination** | Pfizer | Pfizer | Pfizer | Pfizer | Pfizer | Pfizer | Pfizer | Pfizer | Pfizer | Pfizer |
| **Date 1st vaccination** | 12-04-2021 | 13-04-2021 | 13-04-2021 | 11-04-2021 | 15-04-2021 | 13-04-2021 | 11-04-2021 | 11-04-2021 | 11-04-2021 | 19-03-2021 |
| **2nd vaccination** | Pfizer | Pfizer | Pfizer | Pfizer | Pfizer | Pfizer | Pfizer | Pfizer | Pfizer | Pfizer |
| **Date 2nd vaccination** | 10-05-2021 | 11-05-2021 | 11-05-2021 | 09-05-2021 | 16-05-2021 | 11-05-2021 | 09-05-2021 | 10-05-2021 | 09-05-2021 | 24-04-2021 |
| **3rd vaccination** | Pfizer | Pfizer | Pfizer | Pfizer | Pfizer | Pfizer | Pfizer | Pfizer | Pfizer | Pfizer |
| **Date 3rd vaccination** | | 06-01-2022 | | | 22-12-2021 | | 27-12-2021 | 26-12-2021 | | 17-12-2021 |

**Table S3. 1 vs all cluster comparison Wilcoxon test ADT markers**

| Cluster | Marker | logFC | Adjusted P-value (BH) | Significance |
|---|---|---|---|---|
| activated | CD95 (Fas) | 1.273849691 | 2.23E-60 | *** |
| | CD71 | 0.963390715 | 3.27E-47 | *** |
| | CD86 | 1.391485665 | 3.80E-32 | *** |
| | CD58 (LFA-3) | 0.302572327 | 3.86E-31 | *** |
| | CD226 (DNAM-1) | 0.756530911 | 5.60E-25 | *** |
| | CD224 | 0.624959191 | 1.81E-20 | *** |
| | HLA-E | 0.297741685 | 2.34E-18 | *** |
| | CD107a (LAMP-1) | 0.408063413 | 1.70E-14 | *** |
| | CD137 (4-1BB) | 0.181520114 | 4.13E-14 | *** |
| | CD82 | 0.16338323 | 5.64E-12 | *** |
| | CD43 | 0.864753735 | 1.88E-09 | *** |
| | CD2 | 0.472340608 | 1.29E-08 | *** |
| | CD88 (C5aR) | 0.13708966 | 1.65E-08 | *** |
| | CD11c | 0.532077883 | 2.18E-08 | *** |
| | CD307e (FcRL5) | 1.029607244 | 2.76E-08 | *** |
| | CD39 | 0.295050015 | 4.13E-08 | *** |
| | CD123 | 0.505593481 | 2.01E-07 | *** |
| | CD335 (NKp46) | 0.280210539 | 2.30E-07 | *** |
| | CD54 | 0.359867555 | 2.72E-07 | *** |
| | CD124 (IL-4R-) | 0.179381369 | 6.17E-07 | *** |
| | CD27 | 0.446653543 | 3.13E-06 | *** |
| | CD11b | 0.306304193 | 4.12E-06 | *** |
| | CD45RO | 0.19069663 | 1.81E-05 | *** |
| | CD319 (CRACC) | 0.250167503 | 3.39E-05 | *** |
| | CD38 | 0.371380303 | 4.98E-05 | *** |
| | CD3 | 0.328065647 | 0.000195517 | *** |
| | CD119 (IFNgR) | 0.146437531 | 0.000438851 | *** |
| | CD127 (IL-7R) | 0.147152172 | 0.000494886 | *** |
| | CD101 (BB27) | 0.2051976 | 0.000765653 | *** |
| | CD99 | 0.226359764 | 0.001798237 | ** |
| | CD49b | 0.244276053 | 0.00328659 | ** |
| | CD8 | 0.479514479 | 0.004291644 | ** |
| | CD26 | 0.122893383 | 0.01043603 | * |
| | integrin 7 | 0.288852523 | 0.026592874 | * |
| | CLEC12A | 0.425191338 | 0.041631666 | * |
| | CD158e1 (KIR3DL1/NKB1) | 0.234096306 | 0.044758566 | * |
| | CD99 | 0.624069426 | 4.80E-62 | *** |

| | | | | |
|---|---|---|---|---|
| **Classical activated memory** | CD44 | 0.796090694 | 5.12E-61 | *** |
| | CD45RA | 0.44288628 | 3.21E-60 | *** |
| | CD45 | 0.523837589 | 2.25E-44 | *** |
| | CD48 | 0.628692522 | 2.83E-40 | *** |
| | HLA-A,B,C | 0.51600728 | 2.01E-39 | *** |
| | CD11c | 0.691767359 | 2.08E-35 | *** |
| | CD141 (Thrombomodulin) | 0.290661042 | 1.73E-27 | *** |
| | CD47 | 0.483268947 | 3.16E-27 | *** |
| | CD35 | 0.373775294 | 1.43E-23 | *** |
| | CD183 (CXCR3) | 0.336767125 | 6.17E-22 | *** |
| | HLA-DR | 0.327424733 | 2.98E-21 | *** |
| | CD95 (Fas) | 0.366896258 | 2.76E-20 | *** |
| | CD52 | 0.186359538 | 5.26E-19 | *** |
| | CD49d | 0.199413153 | 1.53E-18 | *** |
| | CD195 (CCR5) | 0.229425317 | 1.01E-17 | *** |
| | CD27 | 0.474694099 | 1.05E-15 | *** |
| | CD71 | 0.244687461 | 4.04E-15 | *** |
| | CD319 (CRACC) | 0.113673353 | 3.70E-14 | *** |
| | CD11a | 0.281708816 | 8.01E-14 | *** |
| | CD226 (DNAM-1) | 0.428765016 | 1.21E-13 | *** |
| | CD21 | 0.34686204 | 1.58E-13 | *** |
| | CD22 | 0.314041585 | 1.31E-10 | *** |
| | CD20 | 0.189360951 | 8.55E-10 | *** |
| | CD39 | 0.189082979 | 1.27E-09 | *** |
| | CD62L | 0.538307038 | 2.85E-08 | *** |
| | CD29 | 0.135398923 | 3.40E-08 | *** |
| | CD224 | 0.175614966 | 1.51E-06 | *** |
| | CD274 (B7-H1/PD-L1) | 0.135168391 | 9.21E-06 | *** |
| | CD158b (KIR2DL2-L3/-NKAT2) | 0.159317217 | 9.16E-05 | *** |
| | CD54RB | 0.198869767 | 0.000992383 | *** |
| | CX3CR1 | 0.106798961 | 0.01028359 | * |
| | CD57-Recombinant | 0.171373483 | 0.017258634 | * |
| | CD40 | 0.115416293 | 0.019287469 | * |
| | CD32 | 0.511476062 | 4.45E-25 | *** |
| | CD20 | 0.333899283 | 1.01E-22 | *** |
| | CD11c | 1.135619997 | 3.42E-21 | *** |
| | CD352 (NTB-A) | 0.198403164 | 3.27E-16 | *** |
| | CD307e (FcRL5) | 1.932217872 | 2.03E-12 | *** |

| | | | | |
|---|---|---|---|---|
| atypical | IgD | 0.748439821 | 2.33E-10 | *** |
| | CD5 | 0.547425991 | 1.46E-09 | *** |
| | CD22 | 0.416146747 | 9.96E-09 | *** |
| | CD274 (B7-H1/PD-L1) | 0.211570359 | 1.55E-06 | *** |
| | CD47 | 0.300654478 | 5.49E-06 | *** |
| | CD52 | 0.124003482 | 0.000126643 | *** |
| | CD7 | 0.766717519 | 0.000141657 | *** |
| | CD19 | 0.145239198 | 0.00027838 | *** |
| | IgM | 0.274956209 | 0.000375508 | *** |
| | CD28 | 0.159581045 | 0.000425196 | *** |
| | CD81 (TAPA-1) | 0.122430387 | 0.000497303 | *** |
| | CD4 | 0.328637747 | 0.000689122 | *** |
| | CD107a (LAMP-1) | 0.278950128 | 0.000959468 | *** |
| | CD45 | 0.212558164 | 0.001243184 | ** |
| | CD48 | 0.274900943 | 0.001319349 | ** |
| | CD36 | 0.445819212 | 0.00184711 | ** |
| | CD41 | 0.328287357 | 0.002055481 | ** |
| | CD134 (OX40) | 0.111389764 | 0.016948983 | * |
| | CD101 (BB27) | 0.159138066 | 0.030379496 | * |
| | CD141 (Thrombomodulin) | 0.112828902 | 0.045450382 | * |
| CD24+ intermediate | CD270 (HVEM/TR2) | 0.421455939 | 2.90E-27 | *** |
| | CD1d | 0.650623647 | 9.03E-24 | *** |
| | CD119 (IFN---R---chain) | 0.40266611 | 3.83E-18 | *** |
| | HLA-E | 0.364043099 | 5.71E-18 | *** |
| | CD25 | 0.827141108 | 8.60E-17 | *** |
| | CD85j (ILT2) | 0.28161981 | 2.39E-15 | *** |
| | CD267 (TACI) | 0.229692086 | 3.47E-13 | *** |
| | CD58 (LFA-3) | 0.194082821 | 9.38E-12 | *** |
| | CD54 | 0.513971222 | 1.82E-11 | *** |
| | CD3 | 0.564750373 | 3.42E-11 | *** |
| | CD4 | 0.539430823 | 4.09E-11 | *** |
| | CD272 (BTLA) | 0.200096378 | 7.68E-11 | *** |
| | CD11b | 0.450845486 | 9.24E-11 | *** |
| | CD24 | 0.368347315 | 4.00E-09 | *** |
| | CD38 | 0.342521508 | 1.32E-07 | *** |
| | integrin--7 | 0.304027708 | 4.30E-07 | *** |
| | CD79b (Ig-) | 0.251363399 | 1.49E-06 | *** |
| | CD18 | 0.220436201 | 2.05E-06 | *** |

| | | | | |
|---|---|---|---|---|
|  | TCR---- | 0.548366103 | 1.04E-05 | *** |
| | CLEC12A | 0.585130932 | 3.89E-05 | *** |
| | CD1c | 0.388701076 | 6.51E-05 | *** |
| | CD49f | 0.471932558 | 0.000138165 | *** |
| | CD29 | 0.133656841 | 0.000588977 | *** |
| | CD2 | 0.349100621 | 0.000744138 | *** |
| | CD328 (Siglec-7) | 0.839034477 | 0.007162678 | ** |
| | CD5 | 0.205442991 | 0.017403074 | * |
| | CD8 | 0.632928751 | 0.029908987 | * |
| **CD24+ resting memory** | CD24 | 0.751078351 | 1.09E-61 | *** |
| | CD1d | 0.763930802 | 3.63E-52 | *** |
| | CD270 (HVEM/TR2) | 0.396373958 | 1.11E-45 | *** |
| | CD1c | 0.881117786 | 1.04E-43 | *** |
| | CD25 | 0.837155512 | 5.91E-33 | *** |
| | CD119 (IFN---R---chain) | 0.40411261 | 1.78E-31 | *** |
| | CD267 (TACI) | 0.222699879 | 1.06E-30 | *** |
| | CD85j (ILT2) | 0.265666955 | 8.95E-28 | *** |
| | CD54 | 0.549813884 | 5.80E-25 | *** |
| | integrin--7 | 0.343339377 | 2.04E-19 | *** |
| | CD272 (BTLA) | 0.199194273 | 2.88E-19 | *** |
| | HLA-E | 0.257642586 | 1.55E-18 | *** |
| | CD185 (CXCR5) | 0.309612245 | 1.89E-18 | *** |
| | CD79b (Ig-) | 0.280667922 | 7.22E-16 | *** |
| | CD54RB | 0.31535001 | 8.13E-16 | *** |
| | IgM | 0.515924994 | 1.26E-15 | *** |
| | CD31 | 0.372422156 | 9.11E-09 | *** |
| | CD69 | 0.339285713 | 0.000254629 | *** |
| | CD38 | 0.149145677 | 0.003133934 | ** |
| | CD196 (CCR6) | 0.205514743 | 0.007278606 | ** |
| **CD73+ resting memory** | CD31 | 0.860774397 | 1.93E-49 | *** |
| | CD35 | 0.46368066 | 5.11E-39 | *** |
| | CD11a | 0.410037916 | 7.28E-36 | *** |
| | CD21 | 0.529272895 | 6.48E-34 | *** |
| | CD185 (CXCR5) | 0.39870153 | 1.83E-32 | *** |
| | CD40 | 0.312912839 | 5.61E-30 | *** |
| | CD22 | 0.479133384 | 1.29E-24 | *** |
| | CD268 (BAFF-R) | 0.223358617 | 2.77E-23 | *** |
| | CD73 (Ecto-5'-nucleotidase) | 0.545573365 | 4.08E-19 | *** |
| | CD47 | 0.415992301 | 3.22E-17 | *** |

| | | | | |
|---|---|---|---|---|
| | CD44 | 0.475223866 | 1.01E-16 | *** |
| | CD48 | 0.441092422 | 9.85E-16 | *** |
| | HLA-A,B,C | 0.36596005 | 4.71E-15 | *** |
| | HLA-DR | 0.275903588 | 1.60E-13 | *** |
| | CD32 | 0.279733594 | 2.51E-12 | *** |
| | CD45RA | 0.244396349 | 2.10E-11 | *** |
| | CD274 (B7-H1/PD-L1) | 0.216130444 | 2.60E-11 | *** |
| | CD196 (CCR6) | 0.304175781 | 3.51E-11 | *** |
| | CD45 | 0.31473459 | 4.73E-11 | *** |
| | CD20 | 0.171627694 | 8.91E-08 | *** |
| | CD28 | 0.127970039 | 4.25E-06 | *** |
| | CD141 (Thrombomodulin) | 0.147101992 | 3.65E-05 | *** |
| | CD25 | 0.206893215 | 5.90E-05 | *** |
| | CD54RB | 0.201660552 | 7.67E-05 | *** |
| | IgD | 0.186560685 | 0.000201104 | *** |
| | CD24 | 0.127516466 | 0.00021915 | *** |
| **DN1 activated memory** | CD22 | 0.587757223 | 1.28E-21 | *** |
| | HLA-DR | 0.416041555 | 1.63E-17 | *** |
| | CD62L | 0.786959436 | 1.54E-13 | *** |
| | CD23 | 0.442303351 | 3.78E-13 | *** |
| | CD35 | 0.318423876 | 5.89E-09 | *** |
| | CD81 (TAPA-1) | 0.181792968 | 2.04E-08 | *** |
| | HLA-A,B,C | 0.332521793 | 4.09E-07 | *** |
| | CD19 | 0.184165396 | 1.08E-06 | *** |
| | CD20 | 0.210910129 | 1.49E-06 | *** |
| | CD124 (IL-4R-) | 0.169618041 | 4.62E-05 | *** |
| | CD47 | 0.315489629 | 8.59E-05 | *** |
| | CD352 (NTB-A) | 0.134092829 | 0.000154016 | *** |
| | CD44 | 0.347334289 | 0.000215566 | *** |
| | CD268 (BAFF-R) | 0.149115806 | 0.000386482 | *** |
| | CD11b | 0.230985852 | 0.000503589 | *** |
| | CD45RO | 0.197197208 | 0.00061582 | *** |
| | CD21 | 0.257468837 | 0.006310421 | ** |
| | CD11a | 0.215134998 | 0.006930855 | ** |
| | IgD | 0.189565505 | 0.020795544 | * |
| | CD28 | 0.123860919 | 0.021197273 | * |
| | CD141 (Thrombomodulin) | 0.152791036 | 0.039376044 | * |

**Table S4. 1 vs all cluster comparison Wilcoxon test unsupervised gene analysis**

| Cluster | Gene | logFC | Adjusted P-value (BH) | Significance |
|---|---|---|---|---|
| activated | DHRS9 | 3.157330757 | 0.000857724 | *** |
| | CUX1 | 2.259102855 | 5.45E-11 | *** |
| | GSN | 2.184128765 | 3.03E-17 | *** |
| | AL589693.1 | 1.954247488 | 1.46E-09 | *** |
| | CHAD | 1.707619311 | 0.059863992 | # |
| | ITGAX | 1.503523653 | 0.002970704 | ** |
| | HCST | 1.366975971 | 0.000170114 | *** |
| | ACTG1 | 1.357000921 | 5.95E-28 | *** |
| | S100A10 | 1.323714511 | 2.03E-22 | *** |
| | S100A11 | 1.286844578 | 9.02E-06 | *** |
| | ARPC5 | 1.276259207 | 2.03E-06 | *** |
| | CRIP2 | 1.2585434 | 2.78E-06 | *** |
| | SUB1 | 1.244834513 | 7.93E-18 | *** |
| | ANXA4 | 1.240496347 | 5.12E-08 | *** |
| | JPT1 | 1.240174458 | 3.35E-10 | *** |
| | CRIP1 | 1.239070132 | 4.50E-23 | *** |
| | GAPDH | 1.064121103 | 5.32E-23 | *** |
| | ANXA2 | 1.01485216 | 4.39E-08 | *** |
| | S100A4 | 0.997772761 | 2.83E-07 | *** |
| | ARPC1B | 0.938073524 | 2.10E-13 | *** |
| | CORO1A | 0.930522698 | 2.11E-25 | *** |
| | RFTN1 | 0.922505148 | 0.058393975 | # |
| | CAPZB | 0.910443887 | 2.18E-10 | *** |
| | CAPG | 0.879664117 | 5.32E-08 | *** |
| | TKT | 0.852097193 | 0.004120474 | ** |
| | UCP2 | 0.8430958 | 2.65E-07 | *** |
| | ACTB | 0.841711784 | 1.51E-24 | *** |
| | IFI30 | 0.820819098 | 0.002249571 | ** |
| | PTPN6 | 0.796187816 | 3.89E-07 | *** |
| | TAGLN2 | 0.791460228 | 3.17E-12 | *** |
| | RAC2 | 0.779740055 | 5.57E-15 | *** |
| | RABGAP1L | 0.760405869 | 5.69E-05 | *** |
| | PLPP5 | 0.759616187 | 0.08025768 | # |
| | CD53 | 0.753639861 | 4.67E-06 | *** |
| | CPNE5 | 0.750352022 | 0.003748699 | ** |
| | PFN1 | 0.743071566 | 7.92E-17 | *** |
| | LIMD2 | 0.721062939 | 1.65E-12 | *** |
| | LITAF | 0.720753039 | 0.000177977 | *** |
| | PDE4D | 0.720234297 | 4.56E-07 | *** |
| | CLIC1 | 0.709758106 | 3.26E-11 | *** |

| | | | | |
|---|---|---|---|---|
| | ARPC2 | 0.706228197 | 8.64E-10 | *** |
| | TLE5 | 0.694901928 | 6.37E-05 | *** |
| | MYL12A | 0.671062012 | 2.17E-07 | *** |
| | ARHGDIB | 0.664177723 | 1.35E-12 | *** |
| | ADK | 0.647627648 | 0.01124075 | * |
| | TMSB10 | 0.629133068 | 3.45E-20 | *** |
| | GABARAPL2 | 0.616748964 | 0.0072694 | ** |
| | GSTK1 | 0.612400569 | 0.009305831 | ** |
| | CNN2 | 0.603119249 | 0.020054429 | * |
| | S100A6 | 0.598408902 | 0.0461751 | * |
| | YWHAB | 0.58614421 | 0.004487005 | ** |
| | COTL1 | 0.580456001 | 5.08E-07 | *** |
| | ARPC3 | 0.567328148 | 3.70E-10 | *** |
| | H3F3A | 0.558662521 | 3.66E-07 | *** |
| | LSP1 | 0.544585405 | 4.00E-06 | *** |
| | MYL12B | 0.542973352 | 0.003523987 | ** |
| | LYPLAL1 | 0.538196896 | 4.89E-07 | *** |
| | SH3BGRL3 | 0.529997772 | 2.58E-08 | *** |
| | SEC61B | 0.515839004 | 0.022170733 | * |
| | TMSB4X | 0.515093705 | 5.13E-11 | *** |
| | PPIA | 0.431744101 | 3.00E-05 | *** |
| | CD52 | 0.42058291 | 0.000532836 | *** |
| | ITGB2-AS1 | 0.409458644 | 0.073177561 | # |
| | ATP5F1E | 0.404657411 | 1.75E-06 | *** |
| | LTB | 0.388336325 | 0.071854957 | # |
| | OAZ1 | 0.373962321 | 0.004615634 | ** |
| | SERF2 | 0.343539123 | 0.0123084 | * |
| | HERPUD1 | 0.291219678 | 0.019338306 | * |
| | ITGB2 | 0.257582563 | 0.047053356 | * |
| | SYK | 0.177785718 | 0.05378353 | # |
| | CAMK2D | 0.111317237 | 0.025585908 | * |
| | LINC01934 | 2.876162505 | 1.40E-08 | *** |
| | CHAD | 2.585946558 | 2.05E-05 | *** |
| | CXCR3 | 2.489524733 | 8.72E-17 | *** |
| | GPR25 | 2.449430671 | 3.09E-08 | *** |
| | LGALS3 | 1.837605923 | 0.068654793 | |
| | GRAMD1C | 1.770912973 | 1.22E-11 | *** |
| | ITGB1 | 1.714348371 | 9.05E-31 | *** |
| | COCH | 1.522357255 | 3.97E-16 | *** |
| | S100A10 | 1.494297363 | 1.34E-55 | *** |
| | S100A11 | 1.457045924 | 2.76E-10 | *** |

| | | | | |
|---|---|---|---|---|
| **classical activated memory** | HCST | 1.40238291 | 0.001353218 | ** |
| | ANXA2 | 1.370337223 | 2.66E-21 | *** |
| | JPT1 | 1.322025085 | 2.10E-08 | *** |
| | CRIP2 | 1.320778864 | 3.68E-10 | *** |
| | UBE2N | 1.213556101 | 2.71E-16 | *** |
| | S100A4 | 1.153974251 | 2.44E-21 | *** |
| | ANXA4 | 1.070237614 | 1.21E-06 | *** |
| | LYPLAL1 | 1.062705364 | 6.73E-08 | *** |
| | HOPX | 1.061381692 | 3.53E-06 | *** |
| | CRIP1 | 1.051345671 | 4.13E-31 | *** |
| | HIPK2 | 1.025183352 | 5.86E-05 | *** |
| | CD99 | 0.975749371 | 2.31E-20 | *** |
| | PDE4D | 0.910508212 | 1.62E-09 | *** |
| | GSTK1 | 0.888956625 | 6.91E-14 | *** |
| | TKT | 0.862805683 | 3.96E-10 | *** |
| | AHNAK | 0.830687141 | 1.34E-10 | *** |
| | CPNE5 | 0.808168323 | 4.54E-12 | *** |
| | SELL | 0.759194237 | 7.87E-09 | *** |
| | TAGLN2 | 0.701990417 | 1.74E-15 | *** |
| | ARPC1B | 0.647840615 | 3.55E-12 | *** |
| | MYL12B | 0.647672778 | 0.000131835 | *** |
| | CAPZB | 0.643363923 | 1.11E-06 | *** |
| | HSPA8 | 0.633048346 | 8.21E-12 | *** |
| | S100A6 | 0.632759488 | 5.85E-07 | *** |
| | SUB1 | 0.629588262 | 1.88E-05 | *** |
| | CD53 | 0.586626308 | 2.19E-08 | *** |
| | ISG20 | 0.534664901 | 0.04268945 | * |
| | IFI30 | 0.515504995 | 1.81E-08 | *** |
| | MYL12A | 0.467916868 | 2.73E-05 | *** |
| | CD52 | 0.459239048 | 3.65E-14 | *** |
| | CLIC1 | 0.456233275 | 1.36E-05 | *** |
| | UBE2J1 | 0.450495203 | 0.052738311 | # |
| | GAPDH | 0.446843682 | 9.40E-09 | *** |
| | SH3BGRL3 | 0.407163845 | 4.20E-07 | *** |
| | YWHAB | 0.403429462 | 0.001960681 | ** |
| | RABGAP1L | 0.40271111 | 0.008837478 | ** |
| | TMSB10 | 0.381148826 | 8.59E-12 | *** |
| | TMSB4X | 0.368620991 | 1.34E-09 | *** |
| | PPIA | 0.350360411 | 2.07E-05 | *** |
| | H3F3A | 0.328804424 | 0.007917711 | ** |
| | LTB | 0.327235721 | 1.78E-05 | *** |

| | | | | |
|---|---|---|---|---|
| | ACTB | 0.317925321 | 1.36E-08 | *** |
| | OAZ1 | 0.305828018 | 0.000267757 | *** |
| | PTPN6 | 0.302959486 | 0.017027081 | * |
| | ARHGDIB | 0.296840625 | 0.000440857 | *** |
| | ATP5F1E | 0.294955021 | 0.025678339 | * |
| | PFN1 | 0.249182445 | 0.000913801 | *** |
| | MT-CO3 | 0.242268088 | 0.08621154 | # |
| | ACTG1 | 0.226786551 | 0.003025025 | ** |
| | B2M | 0.189191667 | 0.001412938 | ** |
| | HLA-B | 0.186715015 | 0.031866971 | * |
| | RPLP1 | 0.159982 | 0.019212759 | * |
| atypical | ENC1 | 5.210705983 | 2.18E-06 | *** |
| | SLC11A1 | 5.173365035 | 9.68E-10 | *** |
| | TCL1A | 4.483174166 | 1.25E-05 | *** |
| | DTX1 | 4.085319262 | 3.38E-05 | *** |
| | PCDH9 | 4.031058604 | 1.98E-06 | *** |
| | TAGLN | 4.019482667 | 0.050650523 | # |
| | SOX5 | 3.432696414 | 9.40E-24 | *** |
| | ZNF385A | 3.299132507 | 0.011859054 | * |
| | CD72 | 3.14660625 | 5.16E-19 | *** |
| | FCRL5 | 3.138054341 | 2.66E-21 | *** |
| | MPP6 | 3.079225783 | 1.18E-20 | *** |
| | IGHD | 2.961026672 | 4.37E-07 | *** |
| | FGR | 2.580630114 | 1.79E-19 | *** |
| | ITGAX | 2.549389362 | 0.00369562 | ** |
| | SKAP1 | 2.538762742 | 0.011783171 | * |
| | RIN3 | 2.468723833 | 1.84E-08 | *** |
| | HCK | 2.399740863 | 4.25E-05 | *** |
| | RGS2 | 2.341066193 | 1.74E-11 | *** |
| | FCRL3 | 2.308879502 | 1.14E-11 | *** |
| | ZEB2 | 2.302445863 | 8.59E-23 | *** |
| | CEMIP2 | 2.179004852 | 1.29E-06 | *** |
| | SIGLEC6 | 2.115359364 | 0.004571243 | ** |
| | RHOB | 2.059292324 | 6.09E-23 | *** |
| | MEF2C-AS1 | 1.987505311 | 0.002033765 | ** |
| | HSPB1 | 1.980527157 | 9.93E-10 | *** |
| | ZNF318 | 1.963496157 | 0.067526729 | # |
| | CLEC2B | 1.936260392 | 0.068793097 | # |
| | FCRL2 | 1.879582828 | 3.29E-06 | *** |
| | DUSP5 | 1.835495981 | 0.000942841 | *** |
| | SNX9 | 1.68369777 | 1.75E-05 | *** |

| | | | | |
|---|---|---|---|---|
| | DAPP1 | 1.600912296 | 4.49E-14 | *** |
| | RBM38 | 1.576119375 | 6.73E-11 | *** |
| | CD19 | 1.519172135 | 2.55E-18 | *** |
| | PSAP | 1.515246919 | 1.19E-22 | *** |
| | MAP3K8 | 1.511991544 | 9.59E-14 | *** |
| | CAMK2D | 1.491853543 | 0.000207924 | *** |
| | H1FX | 1.488754723 | 0.027972188 | * |
| | SYK | 1.424539521 | 2.67E-12 | *** |
| | FCRLA | 1.329949242 | 7.36E-10 | *** |
| | NEAT1 | 1.281358284 | 0.001065644 | ** |
| | TENT5C | 1.234261285 | 0.005007672 | ** |
| | FCGR2B | 1.149840017 | 0.006302117 | ** |
| | CIB1 | 1.137961954 | 1.97E-09 | *** |
| | NR4A2 | 1.126012359 | 7.13E-05 | *** |
| | CLEC2D | 1.125837837 | 0.052032231 | # |
| | HSPA5 | 1.117383907 | 1.76E-08 | *** |
| | GSTP1 | 1.107334098 | 0.000839674 | *** |
| | ITGB2 | 1.101328977 | 1.36E-06 | *** |
| | LRMP | 1.099695816 | 0.000118416 | *** |
| | TSPAN3 | 1.089163981 | 1.41E-07 | *** |
| | MTSS1 | 1.015436001 | 0.000241812 | *** |
| | HERPUD1 | 0.979154914 | 0.000992246 | *** |
| | H3F3B | 0.972754524 | 5.35E-11 | *** |
| | EMP3 | 0.965484974 | 3.18E-13 | *** |
| | IFI30 | 0.894699687 | 0.001870317 | ** |
| | LY6E | 0.833135283 | 1.54E-05 | *** |
| | GDI2 | 0.822004148 | 2.32E-05 | *** |
| | LBH | 0.82175571 | 0.000355964 | *** |
| | LITAF | 0.806465487 | 0.01242655 | * |
| | KLF2 | 0.805972332 | 7.28E-09 | *** |
| | HLA-DRB5 | 0.800997679 | 9.37E-08 | *** |
| | LAPTM5 | 0.688198825 | 1.66E-11 | *** |
| | TSC22D3 | 0.66056505 | 4.45E-05 | *** |
| | MS4A1 | 0.627492432 | 4.51E-07 | *** |
| | GABARAPL2 | 0.620106978 | 0.018014101 | * |
| | JUND | 0.585451635 | 0.00012396 | *** |
| | UBC | 0.567374707 | 9.74E-08 | *** |
| | CD79A | 0.557552496 | 0.000201865 | *** |
| | SSR4 | 0.551498817 | 0.050362397 | * |
| | HLA-DRB1 | 0.545218012 | 9.02E-06 | *** |
| | RABGAP1L | 0.51545775 | 0.021371046 | * |

| | | | | |
|---|---|---|---|---|
| | EIF1 | 0.444849843 | 3.66E-05 | *** |
| | FTH1 | 0.438305151 | 1.46E-06 | *** |
| | CD74 | 0.423393297 | 7.24E-25 | *** |
| | HLA-DPA1 | 0.373643666 | 0.000957214 | *** |
| **CD24+ intermediate** | ITM2C | 1.315648661 | 0.035610534 | * |
| | ITM2B | 0.739278909 | 0.023580872 | * |
| | SNHG6 | 0.626014064 | 0.092283137 | # |
| | TXNIP | 0.557470995 | 0.016444407 | * |
| | RPS3 | 0.268786879 | 0.043546593 | * |
| | RPS23 | 0.259517171 | 0.000348522 | *** |
| | RPL30 | 0.222414353 | 0.003420037 | ** |
| | RPS28 | 0.212686221 | 0.002460972 | ** |
| **CD24+ resting memory** | CCSER1 | 2.636422772 | 7.24E-10 | *** |
| | LINC01857 | 2.515999562 | 4.72E-27 | *** |
| | RPS4Y1 | 1.719779315 | 3.14E-17 | *** |
| | DDX3Y | 1.607796155 | 0.00775117 | ** |
| | FOXP1 | 1.221296136 | 5.36E-12 | *** |
| | TENT5C | 1.180301179 | 0.001009801 | ** |
| | CALHM6 | 1.16491469 | 6.97E-07 | *** |
| | CD69 | 1.07468792 | 1.80E-15 | *** |
| | JUNB | 1.01583139 | 3.27E-25 | *** |
| | IGHM | 0.95686152 | 0.001241629 | ** |
| | JCHAIN | 0.952195171 | 1.36E-07 | *** |
| | CD83 | 0.948895147 | 9.66E-09 | *** |
| | ZFP36 | 0.948292666 | 5.43E-22 | *** |
| | PMAIP1 | 0.897199615 | 0.000452399 | *** |
| | YPEL5 | 0.889431258 | 1.61E-06 | *** |
| | NR4A2 | 0.84704183 | 0.005707613 | ** |
| | IER5 | 0.783937663 | 0.001155576 | ** |
| | CXCR4 | 0.72182098 | 0.000698706 | *** |
| | RPS26 | 0.708347 | 1.23E-10 | *** |
| | SNHG7 | 0.639721119 | 0.089733822 | # |
| | DUSP1 | 0.623792159 | 4.64E-11 | *** |
| | NFKBIA | 0.616688174 | 1.38E-06 | *** |
| | PPP1R15A | 0.604743794 | 1.04E-07 | *** |
| | H3F3B | 0.598160136 | 2.05E-10 | *** |
| | JUND | 0.560618381 | 2.49E-13 | *** |
| | FOS | 0.540964183 | 0.004339134 | ** |
| | RALGPS2 | 0.532487695 | 0.00112688 | ** |
| | JUN | 0.460292049 | 0.000521038 | *** |
| | TSC22D3 | 0.393336107 | 9.60E-05 | *** |

| | | | | |
|---|---|---|---|---|
| | RPL3 | 0.28927848 | 3.55E-06 | *** |
| | FTH1 | 0.274970423 | 0.020429704 | * |
| | EIF1 | 0.272806088 | 0.002644909 | ** |
| | RPL9 | 0.268694599 | 5.96E-05 | *** |
| | RPS3 | 0.252408858 | 3.10E-06 | *** |
| | RPL30 | 0.220656176 | 2.97E-09 | *** |
| | RPS9 | 0.213471389 | 0.025463854 | * |
| | RPS27A | 0.198874089 | 9.86E-06 | *** |
| | RPS3A | 0.194104478 | 0.011186814 | * |
| | RPS23 | 0.190535063 | 3.98E-05 | *** |
| | RPL11 | 0.163354921 | 0.000695984 | *** |
| | RPS27 | 0.145620288 | 0.010573341 | * |
| **CD73+ resting memory** | TEX14 | 1.24134571 | 0.00077603 | *** |
| | AC103591.3 | 1.194438631 | 0.001553628 | ** |
| | AC253572.2 | 1.089058732 | 0.000250191 | *** |
| | LINC01781 | 0.924539823 | 0.000492627 | *** |
| | VPREB3 | 0.792650457 | 0.003753358 | *** |
| | TXNIP | 0.717410659 | 1.57E-09 | *** |
| | EEF1B2 | 0.55643415 | 8.35E-22 | *** |
| | GAS5 | 0.555986589 | 2.62E-07 | *** |
| | RPS3 | 0.495936821 | 4.91E-33 | *** |
| | SNHG29 | 0.47791865 | 3.73E-05 | *** |
| | RPS12 | 0.393945338 | 1.44E-28 | *** |
| | RPL3 | 0.375085118 | 9.70E-16 | *** |
| | RPL10 | 0.368025599 | 4.22E-25 | *** |
| | RPS23 | 0.350209221 | 6.31E-23 | *** |
| | TPT1 | 0.337117273 | 2.20E-09 | *** |
| | RPL36A | 0.32984766 | 1.23E-06 | *** |
| | RPL38 | 0.326595226 | 3.00E-07 | *** |
| | RPL35A | 0.326568453 | 5.63E-15 | *** |
| | RPLP0 | 0.320331603 | 1.85E-08 | *** |
| | PFDN5 | 0.315599355 | 0.002258896 | ** |
| | RPS4X | 0.309207371 | 4.68E-08 | *** |
| | RPS2 | 0.303184397 | 1.93E-12 | *** |
| | RPSA | 0.299365095 | 3.64E-07 | *** |
| | RPS5 | 0.293157928 | 2.64E-07 | *** |
| | EEF1G | 0.291476354 | 0.022451448 | * |
| | RPL9 | 0.290668919 | 1.14E-08 | *** |
| | RPS14 | 0.290480034 | 1.38E-14 | *** |
| | RPS13 | 0.284953676 | 1.95E-12 | *** |
| | RPL14 | 0.278872955 | 6.45E-10 | *** |

| | | | | |
|---|---|---|---|---|
| | RPS28 | 0.276319851 | 2.67E-18 | *** |
| | RPS25 | 0.273708368 | 1.52E-10 | *** |
| | RPS9 | 0.265882328 | 2.54E-05 | *** |
| | RPS8 | 0.265519477 | 4.75E-13 | *** |
| | RPS27A | 0.265261403 | 6.23E-13 | *** |
| | RPS18 | 0.263160131 | 6.65E-08 | *** |
| | RPS3A | 0.263029457 | 5.21E-09 | *** |
| | RPL8 | 0.261856654 | 2.27E-09 | *** |
| | RPL34 | 0.256344192 | 2.47E-15 | *** |
| | RPS15A | 0.255381391 | 1.04E-13 | *** |
| | RPL32 | 0.254793146 | 2.65E-15 | *** |
| | RPL30 | 0.251063111 | 1.59E-13 | *** |
| | EEF1A1 | 0.245770562 | 5.45E-12 | *** |
| | RPL18 | 0.238427088 | 2.69E-09 | *** |
| | RPL26 | 0.237489896 | 5.04E-08 | *** |
| | RPL17 | 0.234956983 | 0.003479579 | ** |
| | RPS6 | 0.230688841 | 3.99E-05 | *** |
| | RPLP1 | 0.229876559 | 2.00E-06 | *** |
| | RPS21 | 0.223631712 | 1.62E-07 | *** |
| | RPL5 | 0.222123155 | 0.007526786 | ** |
| | FAU | 0.218387989 | 1.91E-07 | *** |
| | NACA | 0.215931339 | 0.010976011 | * |
| | RPL11 | 0.204430781 | 4.16E-08 | *** |
| | RPL10A | 0.20412185 | 0.022120852 | * |
| | RPL28 | 0.202425746 | 7.80E-08 | *** |
| | RACK1 | 0.19802712 | 0.011389479 | * |
| | RPL21 | 0.191648625 | 0.024547954 | * |
| | RPL13 | 0.191297329 | 3.69E-07 | *** |
| | RPL39 | 0.184123442 | 8.49E-07 | *** |
| | RPLP2 | 0.177261276 | 0.09804584 | # |
| | RPS27 | 0.17274067 | 0.000123953 | *** |
| | RPL19 | 0.15474756 | 0.000535151 | *** |
| DN1 activated memory | IL4R | 3.441090704 | 6.37E-19 | *** |
| | FCER2 | 2.179251153 | 1.02E-25 | *** |
| | HOPX | 1.421442019 | 2.60E-05 | *** |
| | PLPP5 | 1.244236047 | 0.00486292 | ** |
| | PDE4D | 1.053636134 | 1.17E-06 | *** |
| | PPP1R14A | 1.018358244 | 0.077273119 | # |
| | SELL | 1.012469736 | 1.23E-12 | *** |
| | LYPLAL1 | 0.969910727 | 0.003887349 | ** |
| | MEF2C | 0.897884198 | 3.02E-07 | *** |

| | | | | |
|---|---|---|---|---|
| | ADK | 0.872356515 | 0.000364931 | *** |
| | CD53 | 0.631342364 | 0.015220325 | * |
| | LAPTM5 | 0.509987111 | 5.73E-06 | *** |
| | TMSB10 | 0.30898028 | 0.025132409 | * |

**Table S5. 1 vs all cluster comparison Wilcoxon test supervised gene analysis**

| Cluster | Gene | logFC | Adjusted P-value (BH) | Significance |
|---|---|---|---|---|
| activated | EPHA4 | 1.889282296 | 7.06E-05 | *** |
| | S1PR4 | 1.140587898 | 1.55E-05 | *** |
| | CD70 | 1.090209358 | 2.87E-05 | *** |
| | SASH3 | 0.706935972 | 0.021510657 | * |
| | PRKCD | 0.565398089 | 0.006516181 | ** |
| | BLK | 0.540483175 | 0.006031498 | ** |
| | RHBDF2 | 0.516765891 | 0.00500964 | ** |
| | CYBB | 0.378547505 | 0.002113359 | ** |
| | SSPN | 0.308297624 | 5.15E-05 | *** |
| classical activated memory | SSPN | 1.114920531 | 0.045821731 | * |
| | PARM1 | 1.024422994 | 0.002161381 | ** |
| | TMEM154 | 0.754064624 | 0.001518188 | ** |
| | BLK | 0.5309684 | 0.000760119 | *** |
| | HLA-DMB | 0.385381333 | 0.028307804 | * |
| atypical | GPR18 | 1.972454732 | 0.028271744 | * |
| | TNFRSF1B | 1.454674396 | 0.043300074 | * |
| CD24+ resting memory | SLC2A3 | 1.000556027 | 8.37E-05 | *** |
| DN1 activated memory | HLA-DMA | 0.746593876 | 5.28E-06 | *** |

**Table S6. 1 vs all cluster comparison Wilcoxon test PROGENy pathway analysis**

| Cluster | Pathway | Mean cluster | Mean other | Adjusted P-value (BH) | Significance |
|---|---|---|---|---|---|
| activated | EGFR | 1.5407 | -0.0763 | 0 | ns |
| | JAK-STAT | 0.8714 | -0.0433 | 0 | ns |
| | TNFa | 1.2113 | -0.06 | 0 | ns |
| | NFkB | -1.198 | 0.0593 | 0 | ns |
| | MAPK | -1.3721 | 0.0679 | 0 | ns |
| | TGFb | -0.7322 | 0.0362 | 0 | ns |
| | VEGF | -0.7171 | 0.0355 | 0 | ns |
| | Hypoxia | 0.6685 | -0.0331 | 0 | ns |
| | Estrogen | 0.4741 | -0.0235 | 0 | ns |
| | Androgen | -0.3909 | 0.0193 | 0 | ns |
| | WNT | -0.2805 | 0.0139 | 0.0004 | *** |
| | PI3K | 0.2098 | -0.0104 | 0.0104 | ** |
| | Trail | -0.1479 | 0.0073 | 0.2854 | ns |
| | p53 | 0.0742 | -0.0037 | 0.6668 | ns |
| classical activated memory | EGFR | 0.7625 | -0.0687 | 0 | ns |
| | JAK-STAT | 0.7352 | -0.0664 | 0 | ns |
| | TNFa | 1.1241 | -0.1012 | 0 | ns |
| | NFkB | -1.1034 | 0.0994 | 0 | ns |
| | MAPK | -0.7932 | 0.0714 | 0 | ns |
| | TGFb | -0.5586 | 0.0503 | 0 | ns |
| | VEGF | -0.5687 | 0.0512 | 0 | ns |
| | Estrogen | 0.4609 | -0.0415 | 0 | ns |
| | Androgen | -0.3306 | 0.0298 | 0 | ns |
| | WNT | -0.285 | 0.0257 | 0 | ns |
| | p53 | -0.2284 | 0.0206 | 0.0001 | *** |
| | Trail | -0.1271 | 0.0114 | 0.0328 | * |
| | Hypoxia | 0.0856 | -0.0077 | 0.0704 | # |
| | PI3K | -0.0481 | 0.0043 | 0.3515 | ns |
| atypical | JAK-STAT | 0.8136 | -0.0408 | 0 | ns |
| | TNFa | 0.6571 | -0.0328 | 0 | ns |
| | NFkB | -0.7798 | 0.0389 | 0 | ns |
| | EGFR | 0.44 | -0.0219 | 0 | ns |
| | Androgen | -0.4738 | 0.0236 | 0 | ns |
| | PI3K | 0.3971 | -0.0198 | 0 | ns |
| | Hypoxia | -0.338 | 0.0169 | 0.0001 | *** |
| | Trail | -0.3093 | 0.0154 | 0.0002 | *** |
| | MAPK | -0.2995 | 0.0149 | 0.0006 | *** |
| | WNT | -0.2452 | 0.0122 | 0.0026 | ** |

| | | | | | |
|---|---|---|---|---|---|
| | VEGF | -0.153 | 0.0076 | 0.2075 | ns |
| | TGFb | 0.1265 | -0.0063 | 0.2303 | ns |
| | Estrogen | -0.0491 | 0.0024 | 0.7455 | ns |
| | p53 | 0.0346 | -0.0017 | 0.7737 | ns |
| **CD24+ intermediate** | Hypoxia | -0.6138 | 0.0208 | 0 | ns |
| | PI3K | -0.4177 | 0.0141 | 0 | ns |
| | NFkB | -0.4564 | 0.0154 | 0 | ns |
| | Estrogen | 0.4303 | -0.0145 | 0 | ns |
| | TNFa | 0.3946 | -0.0133 | 0 | ns |
| | TGFb | -0.3945 | 0.0133 | 0 | ns |
| | Androgen | -0.1394 | 0.0047 | 0.1463 | ns |
| | p53 | -0.0773 | 0.0026 | 0.2996 | ns |
| | VEGF | -0.104 | 0.0035 | 0.4042 | ns |
| | JAK-STAT | 0.0535 | -0.002 | 0.6187 | ns |
| | MAPK | -0.0038 | 0.0001 | 0.7737 | ns |
| | EGFR | -0.0143 | 0.0005 | 0.8054 | ns |
| | Trail | -0.0283 | 0.001 | 0.8468 | ns |
| | WNT | -0.0054 | 0.0002 | 0.9995 | ns |
| **CD24+ resting memory** | TNFa | 0.5809 | -0.0423 | 0 | ns |
| | Hypoxia | -0.5585 | 0.0406 | 0 | ns |
| | TGFb | 0.5337 | -0.0388 | 0 | ns |
| | Trail | -0.3009 | 0.0219 | 0 | ns |
| | VEGF | -0.2821 | 0.0205 | 0 | ns |
| | JAK-STAT | -0.226 | 0.0163 | 0.0054 | ** |
| | NFkB | -0.1606 | 0.0117 | 0.007 | ** |
| | Androgen | 0.1735 | -0.0126 | 0.037 | * |
| | WNT | 0.1122 | -0.0082 | 0.0648 | # |
| | PI3K | -0.0723 | 0.0053 | 0.1369 | ns |
| | EGFR | 0.073 | -0.0053 | 0.1517 | ns |
| | Estrogen | 0.1469 | -0.0107 | 0.1727 | ns |
| | p53 | 0.1058 | -0.0077 | 0.3928 | ns |
| | MAPK | -0.0233 | 0.0017 | 0.6187 | ns |
| **CD73+ resting memory** | Hypoxia | -0.7551 | 0.0582 | 0 | ns |
| | PI3K | -0.5806 | 0.0448 | 0 | ns |
| | TNFa | 0.2656 | -0.0205 | 0 | ns |
| | TGFb | -0.3264 | 0.0252 | 0 | ns |
| | NFkB | -0.2705 | 0.0209 | 0 | ns |
| | Estrogen | 0.2797 | -0.0216 | 0 | ns |
| | JAK-STAT | -0.2755 | 0.0211 | 0 | ns |
| | Trail | -0.2027 | 0.0156 | 0.0013 | ** |
| | EGFR | 0.1136 | -0.0088 | 0.0099 | ** |

| | | | | | |
|---|---|---|---|---|---|
| | p53 | 0.2066 | -0.0159 | 0.0183 | * |
| | VEGF | -0.112 | 0.0086 | 0.0482 | * |
| | WNT | -0.1161 | 0.009 | 0.0767 | # |
| | Androgen | 0.064 | -0.0049 | 0.4042 | ns |
| | MAPK | -0.0058 | 0.0005 | 0.8054 | ns |
| **DN1 activated memory** | TGFb | -0.6339 | 0.0233 | 0 | ns |
| | EGFR | 0.6928 | -0.0255 | 0 | ns |
| | JAK-STAT | 0.748 | -0.0277 | 0 | ns |
| | TNFa | 0.5748 | -0.0211 | 0 | ns |
| | NFkB | -0.4749 | 0.0175 | 0 | ns |
| | MAPK | -0.3229 | 0.0119 | 0.0001 | *** |
| | WNT | -0.1697 | 0.0062 | 0.1199 | ns |
| | p53 | 0.1655 | -0.0061 | 0.1862 | ns |
| | Androgen | 0.1164 | -0.0043 | 0.5063 | ns |
| | VEGF | 0.0799 | -0.0029 | 0.6187 | ns |
| | PI3K | -0.0709 | 0.0026 | 0.7832 | ns |
| | Trail | -0.0176 | 0.0006 | 0.9082 | ns |
| | Estrogen | -0.0447 | 0.0016 | 0.9995 | ns |
| | Hypoxia | -0.0286 | 0.001 | 0.9995 | ns |
| **Naive** | Hypoxia | 0.1471 | -0.2351 | 0 | ns |
| | MAPK | 0.2569 | -0.4106 | 0 | ns |
| | NFkB | 0.4011 | -0.6411 | 0 | ns |
| | TGFb | 0.1581 | -0.2526 | 0 | ns |
| | VEGF | 0.1882 | -0.3009 | 0 | ns |
| | TNFa | -0.4436 | 0.7091 | 0 | ns |
| | EGFR | -0.3149 | 0.5034 | 0 | ns |
| | JAK-STAT | -0.2176 | 0.3474 | 0 | ns |
| | Estrogen | -0.1635 | 0.2614 | 0 | ns |
| | Trail | 0.1116 | -0.1784 | 0 | ns |
| | Androgen | 0.0851 | -0.136 | 0 | ns |
| | WNT | 0.0899 | -0.1437 | 0 | ns |
| | PI3K | 0.0616 | -0.0984 | 0 | ns |
| | p53 | -0.0188 | 0.0301 | 0.9995 | ns |

**Table S7. Pairwise Wilcoxon Test of SHM levels between B cell subsets**

| CD45RB status | Cluster 1 | Cluster 2 | N cells cluster 1 | N cells cluster 2 | Adjusted P-value (BH) | Significance |
|---|---|---|---|---|---|---|
| CD45RB+ | activated | CD73+ resting memory | 20 | 25 | 0.491 | ns |
| | | CD24+ resting memory | 20 | 16 | 0.759 | ns |
| | | classical activated memory | 20 | 42 | 0.046 | * |
| | | atypical | 20 | 68 | 0.396 | ns |
| | | DN1 activated memory | 20 | 51 | 0.497 | ns |
| | | $CD24^{+}$ intermediate | 20 | 20 | 0.265 | ns |
| | CD73+ resting memory | CD24+ resting memory | 25 | 16 | 0.505 | ns |
| | | classical activated memory | 25 | 42 | 0.497 | ns |
| | | atypical | 25 | 68 | 0.052 | # |
| | | DN1 activated memory | 25 | 51 | 0.682 | ns |
| | | $CD24^{+}$ intermediate | 25 | 20 | 0.682 | ns |
| | CD24+ resting memory | classical activated memory | 16 | 42 | 0.109 | ns |
| | | atypical | 16 | 68 | 0.396 | ns |
| | | DN1 activated memory | 16 | 51 | 0.682 | ns |
| | | $CD24^{+}$ intermediate | 16 | 20 | 0.396 | ns |
| | classical activated memory | atypical | 42 | 68 | 0.000813 | *** |
| | | DN1 activated memory | 42 | 51 | 0.064 | # |
| | | $CD24^{+}$ intermediate | 42 | 20 | 0.759 | ns |
| | atypical | DN1 activated memory | 68 | 51 | 0.046 | * |
| | | $CD24^{+}$ intermediate | 68 | 20 | 0.046 | * |
| | DN1 activated memory | $CD24^{+}$ intermediate | 51 | 20 | 0.396 | ns |
| CD45RB+ | activated | CD73+ resting memory | 191 | 156 | 0.862 | ns |

| | | | | | | |
|---|---|---|---|---|---|---|
| | | CD24+ resting memory | 191 | 167 | 0.005 | ** |
| | | classical activated memory | 191 | 95 | 3.98E-07 | *** |
| | | atypical | 191 | 63 | 0.405 | ns |
| | | DN1 activated memory | 191 | 31 | 0.48 | ns |
| | | CD24$^+$ intermediate | 191 | 64 | 0.49 | ns |
| | **CD73+ resting memory** | CD24+ resting memory | 156 | 167 | 0.005 | ** |
| | | classical activated memory | 156 | 95 | 1.00E-06 | *** |
| | | atypical | 156 | 63 | 0.38 | ns |
| | | DN1 activated memory | 156 | 31 | 0.405 | ns |
| | | CD24$^+$ intermediate | 156 | 64 | 0.538 | ns |
| | **CD24+ resting memory** | classical activated memory | 167 | 95 | 7.31E-12 | *** |
| | | atypical | 167 | 63 | 0.538 | ns |
| | | DN1 activated memory | 167 | 31 | 0.49 | ns |
| | | CD24$^+$ intermediate | 167 | 64 | 0.005 | ** |
| | **classical activated memory** | atypical | 95 | 63 | 3.56E-05 | *** |
| | | DN1 activated memory | 95 | 31 | 6.47E-05 | *** |
| | | CD24$^+$ intermediate | 95 | 64 | 0.000882 | *** |
| | **atypical** | DN1 activated memory | 63 | 31 | 0.949 | ns |
| | | CD24$^+$ intermediate | 63 | 64 | 0.237 | ns |
| | **DN1 activated memory** | CD24$^+$ intermediate | 31 | 64 | 0.246 | ns |

**Table S8. Linear mixed effects model of SARS-CoV-2 spike$^+$ IgG$^+$ B cells**

| Timepoint 1 | Timepoint 2 | Adjusted P-value (BH) | significance |
|---|---|---|---|
| **1-2 months post vaccination 2** | 1 month post infection | 0.0250438448382725 | * |
| | 1 month post vaccination 1 | 0.00526146204052931 | ** |
| | 1 week post infection | 0.00723063373944098 | ** |

| | | | |
|---|---|---|---|
| | 1 week post vaccination 1 | 0.968228173115455 | ns |
| | 1 week post vaccination 2 | 0.0448729791928401 | * |
| | 3-8 months post vaccination 2 | 0.336670719610713 | ns |
| | 6-7 months post infection | 0.0611806862405175 | ns |
| | 8-10 months post infection | 0.0353611317654291 | * |
| **1 month post infection** | 1 month post vaccination 1 | 1.48E+08 | **** |
| | 1 week post infection | 0.597440997858221 | ns |
| | 1 week post vaccination 1 | 0.0353611317654291 | * |
| | 1 week post vaccination 2 | 0.000145806624307372 | *** |
| | 3-8 months post vaccination 2 | 0.305911057223993 | ns |
| | 6-7 months post infection | 0.639334793913884 | ns |
| | 8-10 months post infection | 0.826482522438372 | ns |
| **1 month post vaccination 1** | 1 week post infection | 1.47E+09 | **** |
| | 1 week post vaccination 1 | 0.00736698378176467 | ** |
| | 1 week post vaccination 2 | 0.350593566322701 | ns |
| | 3-8 months post vaccination 2 | 0.00039913677388354 | *** |
| | 6-7 months post infection | 2.82E+09 | **** |
| | 8-10 months post infection | 1.48E+08 | **** |
| **1 week post infection** | 1 week post vaccination 1 | 0.0105913183946718 | * |
| | 1 week post vaccination 2 | 3.56E+09 | **** |
| | 3-8 months post vaccination 2 | 0.106421957663111 | ns |
| | 6-7 months post infection | 0.340787164410738 | ns |
| | 8-10 months post infection | 0.450451317684514 | ns |
| **1 week post vaccination 1** | 1 week post vaccination 2 | 0.0580151048929591 | ns |
| | 3-8 months post vaccination 2 | 0.350593566322701 | ns |
| | 6-7 months post infection | 0.0843005960914319 | ns |
| | 8-10 months post infection | 0.0477796032913487 | * |
| **1 week post vaccination 2** | 3_8_months_post_vacc_2 | 0.00577463125065915 | ** |
| | 6-7 months post infection | 0.000339770612952505 | *** |
| | 8-10 months post infection | 0.000145806624307372 | *** |
| **3-8 months post vaccination 2** | 6-7 months post infection | 0.488363409312986 | ns |
| | 8-10 months post infection | 0.350593566322701 | ns |

| | | | |
|---|---|---|---|
| **6-7 months post infection** | 8-10 months post infection | 0.800344820388621 | ns |

**Table S9. Pairwise Wilcoxon Test of cell surface marker expression per B cell subset**

| Cluster | Marker | logFC | Adjusted P-value (BH) | Significance |
|---|---|---|---|---|
| activated | CD40 | 0.59338999975 | 0.00161273376144088 | ** |
| | CD38 | 20,416,950,505 | 4.60E+08 | *** |
| | CD86 | 0.051277151 | 0.484924279417111 | ns |
| | CD268 | 0.0681254025 | 0.749000800118647 | ns |
| | CD24 | -0.9954921265 | 0.000125347863267036 | *** |
| | HLA-DR | 1,333,299,418 | 4.28E+08 | *** |
| | IgD | -0.1648888625 | 0.12605741056499 | ns |
| | CD43 | 233,214,630,725 | 3.65E+08 | *** |
| | CD22 | 0.212019759 | 0.604146381163383 | ns |
| | CD21 | -0.765834877 | 0.022562085058195 | * |
| | CD27 | 12,362,984,545 | 3.02E+07 | *** |
| | CD11c | 0.633683036 | 0.000171920990443541 | *** |
| | CD73 | -0.6193909845 | 0.000629427496692098 | *** |
| | CD11a | 0.3828086715 | 0.0695158343556909 | ns |
| | CD71 | 27,885,423,155 | 3.65E+08 | *** |
| | FCRL5 | 18,555,683,705 | 5.92E+08 | *** |
| | CD45RB | 0.11117975925 | 0.991475366491254 | ns |
| | IgG | 0.09166665475 | 0.670831445601111 | ns |
| | CD95 | 198,433,580,925 | 3.65E+08 | *** |
| | CD99 | 0.91009326725 | 0.000125347863267036 | *** |
| | CD20 | 0.400873109 | 0.0305434215617656 | * |
| | CD1d | 0.75449613675 | 6.06E+09 | *** |
| | IgM | 0.53508132 | 0.000516343401098703 | *** |
| activated memory | CD40 | 0.5540718855 | 0.0159664763817339 | * |
| | CD38 | 0.938742681 | 0.0064905713539879 | ** |
| | CD86 | -0.4649389575 | 0.000378329643087851 | *** |
| | CD268 | 0.82403777775 | 9.71E+09 | *** |
| | CD24 | 0.05146213925 | 0.967829757774563 | ns |
| | HLA-DR | 0.8376598155 | 0.00010004627342838 | *** |
| | IgD | 0.10945985925 | 0.042826689053636 | * |
| | CD43 | 0.15838920175 | 0.0159664763817339 | * |
| | CD22 | 0.72548071425 | 0.000166825098727766 | *** |
| | CD21 | 0.83735144425 | 0.00122273615055192 | ** |

| | | | | |
|---|---|---|---|---|
| | CD27 | 0.1751545305 | 0.657447777745408 | ns |
| | CD11c | -0.00234636225000001 | 0.991475366491254 | ns |
| | CD73 | -0.668786396 | 6.45E+08 | *** |
| | CD11a | 0.451786692 | 0.0244543625955012 | * |
| | CD71 | 0.39561654525 | 0.011313264203099 | * |
| | FCRL5 | -0.0825077895 | 0.0194500698451712 | * |
| | CD45RB | 0.07759450825 | 0.923987521727834 | ns |
| | IgG | -0.04258893 | 0.743826712297475 | ns |
| | CD95 | -0.3925834495 | 0.414235965453174 | ns |
| | CD99 | 0.0925134115 | 0.396707155598123 | ns |
| | CD20 | 0.01244766475 | 0.466130994340294 | ns |
| | CD1d | 0.21751703775 | 0.241692594584887 | ns |
| | IgM | 0.414890917 | 0.00924705696678 79 | ** |
| **CD24+ intermediate** | CD40 | -0.38647838225 | 0.337643177108909 | ns |
| | CD38 | -0.4558639005 | 0.000341928916802381 | *** |
| | CD86 | 0.055723763 | 0.59907908281269 | ns |
| | CD268 | -0.10174020525 | 0.504102978672123 | ns |
| | CD24 | 0.7714529675 | 0.0037516686991614 | ** |
| | HLA-DR | 0.29877280775 | 0.0859546552552891 | ns |
| | IgD | -0.093990415 | 0.12605741056499 | ns |
| | CD43 | 0.05790549725 | 0.548238375023249 | ns |
| | CD22 | -0.1748119975 | 0.59907908281269 | ns |
| | CD21 | -0.78693371425 | 0.0168449068606499 | * |
| | CD27 | 0.7200141805 | 0.0597948231684804 | ns |
| | CD11c | 0.31424820075 | 0.00495910404436822 | ** |
| | CD73 | -0.1293330575 | 0.672929381565716 | ns |
| | CD11a | -0.99201634175 | 0.000540154469705394 | *** |
| | CD71 | 0.3505113465 | 0.0282474036015949 | * |
| | FCRL5 | 0.24748041375 | 0.00521475655795886 | ** |
| | CD45RB | -0.1832601735 | 0.543296838775533 | ns |
| | IgG | -0.27757679925 | 0.0669656087010243 | ns |
| | CD95 | 0.74790281675 | 0.0311965639450362 | * |
| | CD99 | 0.51544363825 | 0.00466877383559301 | ** |
| | CD20 | 0.45801504125 | 0.00804301262551147 | ** |
| | CD1d | 0.5813634075 | 0.000794357758305584 | *** |
| | IgM | 0.2964334445 | 0.041967769892663 8 | * |
| | CD40 | -0.538548368 | 0.0721461422468984 | ns |

| | | | | |
|---|---|---|---|---|
| CD24+ resting memory | CD38 | -0.42930754675 | 0.00196177181046124 | ** |
| | CD86 | -0.378952079 | 0.00369520923412828 | ** |
| | CD268 | -0.259750672 | 0.107831105108175 | ns |
| | CD24 | 186,781,094,325 | 3.65E+08 | *** |
| | HLA-DR | -0.6786298515 | 0.00098707486926102 4 | *** |
| | IgD | 0.053938637 | 0.0785721333421502 | ns |
| | CD43 | -0.01222550375 | 0.90954243356585 | ns |
| | CD22 | -102,191,031,075 | 0.00012534786326703 6 | *** |
| | CD21 | 0.064257227 | 0.986568295087484 | ns |
| | CD27 | 0.9803429355 | 0.0024313515512883 | ** |
| | CD11c | -0.15162948975 | 0.0341692489033879 | * |
| | CD73 | -0.115942879 | 0.807383551453535 | ns |
| | CD11a | -0.5816379135 | 0.20025473369077 | ns |
| | CD71 | -0.56737940875 | 0.0017891667198542 | ** |
| | FCRL5 | -0.02312226475 | 0.259069730138414 | ns |
| | CD45RB | 13,192,788,535 | 6.45E+08 | *** |
| | IgG | 0.315416237 | 0.0092470569667879 | ** |
| | CD95 | 0.5102434185 | 0.265917203651108 | ns |
| | CD99 | -0.08698618325 | 0.284445521581369 | ns |
| | CD20 | 0.11074889075 | 0.708160542165301 | ns |
| | CD1d | -0.13186505525 | 0.318530696373466 | ns |
| | IgM | -0.3186594515 | 0.0499040202237808 | * |
| CD73+ resting memory | CD40 | 0.667264735 | 0.00042402281609695 2 | *** |
| | CD38 | 122,931,048,875 | 9.71E+09 | *** |
| | CD86 | -0.673615077 | 4.60E+08 | *** |
| | CD268 | 0.96276555875 | 1.32E+09 | *** |
| | CD24 | 0.6349579485 | 0.0215870143812683 | * |
| | HLA-DR | 0.30569452025 | 0.12605741056499 | ns |
| | IgD | 0.20049562425 | 0.00054015446970539 4 | *** |
| | CD43 | -0.06379906 | 0.713337622277255 | ns |
| | CD22 | 0.3924815965 | 0.0590278675796913 | ns |
| | CD21 | 102,868,311,575 | 3.65E+08 | *** |
| | CD27 | 0.116843352 | 0.824206284672894 | ns |
| | CD11c | -0.2220009215 | 5.92E+08 | *** |
| | CD73 | 168,556,413,325 | 6.21E+08 | *** |
| | CD11a | 0.806573423 | 3.65E+08 | *** |
| | CD71 | -0.1332956755 | 0.506468590872323 | ns |
| | FCRL5 | 0.05715616175 | 0.359298259662874 | ns |

| | | | | |
|---|---|---|---|---|
| | CD45RB | 12,616,276,675 | 9.71E+09 | *** |
| | IgG | 0.6085142215 | 4.54E+08 | *** |
| | CD95 | -0.627303697 | 0.0341692489033879 | * |
| | CD99 | 0.01157122225 | 0.895992976051756 | ns |
| | CD20 | -0.17228588525 | 0.0721461422468984 | ns |
| | CD1d | 0.091234254 | 0.686341778808527 | ns |
| | IgM | 0.0214553405 | 0.860119031861107 | ns |
| **CD73+ activated** | CD40 | 0.4632153235 | 0.194577978807963 | ns |
| | CD38 | -0.67616163625 | 3.65E+08 | *** |
| | CD86 | 0.193723093 | 0.0198905043867905 | * |
| | CD268 | 0.565745912 | 0.0276479659239434 | * |
| | CD24 | 0.13526989375 | 0.914382623680031 | ns |
| | HLA-DR | -0.23534327175 | 0.846475399170639 | ns |
| | IgD | 0.328669892 | 1.29E+09 | *** |
| | CD43 | 0.2635365145 | 0.00693066825809647 | ** |
| | CD22 | 0.05941223275 | 0.969099564462485 | ns |
| | CD21 | 0.84231135975 | 0.00139721548357745 | ** |
| | CD27 | 119,218,094 | 2.94E+09 | *** |
| | CD11c | 0.15638678675 | 0.122703310326845 | ns |
| | CD73 | 15,755,165,325 | 6.06E+09 | *** |
| | CD11a | 0.52495895275 | 0.000367440411638548 | *** |
| | CD71 | 0.833574315 | 0.000125347863267036 | *** |
| | FCRL5 | -0.33184569775 | 3.65E+08 | *** |
| | CD45RB | 1,145,684,417 | 0.00335019843602574 | ** |
| | IgG | 0.233133679 | 0.147383312556268 | ns |
| | CD95 | 14,101,485,955 | 0.000125347863267036 | *** |
| | CD99 | 11,985,656,605 | 3.65E+08 | *** |
| | CD20 | -0.03979012625 | 0.41867403932195 | ns |
| | CD1d | 0.78534381375 | 5.86E+09 | *** |
| | IgM | 0.00771867449999999 | 0.822687647294322 | ns |
| **CD73+/CD24+ resting memory** | CD40 | -0.05194388 | 0.968466381573275 | ns |
| | CD38 | -0.12600294675 | 0.530897759333413 | ns |
| | CD86 | -0.09368889175 | 0.576152844267732 | ns |
| | CD268 | 0.0518402675 | 0.968466381573275 | ns |
| | CD24 | 0.79555648825 | 0.000885832147456208 | *** |
| | HLA-DR | -0.48220717625 | 0.013569217950915 | * |
| | IgD | 0.0430320395 | 0.822687647294322 | ns |

| | | | | |
|---|---|---|---|---|
| | CD43 | -0.30050332975 | 0.00398938181343532 | ** |
| | CD22 | -0.46584352525 | 0.0373827553578212 | * |
| | CD21 | 0.399755933 | 0.414001727290772 | ns |
| | CD27 | -0.181667833 | 0.466130994340294 | ns |
| | CD11c | -0.18904729925 | 0.00105630285480404 | ** |
| | CD73 | 11,933,446,765 | 0.0448818384712203 | * |
| | CD11a | -0.03215810375 | 0.973638680849234 | ns |
| | CD71 | -0.489535574 | 0.0457910002932598 | * |
| | FCRL5 | -0.0817332385 | 0.00804301262551147 | ** |
| | CD45RB | 10,554,884,905 | 0.053609004015601 | ns |
| | IgG | 0.3754693185 | 0.0037516686991614 | ** |
| | CD95 | -0.7293413395 | 0.00978843870868639 | ** |
| | CD99 | -0.26035123075 | 0.00653506203817657 | ** |
| | CD20 | -0.048048136 | 0.379847899183751 | ns |
| | CD1d | -0.3257461285 | 0.00734784895813309 | ** |
| | IgM | -0.4383114645 | 0.00466877383559301 | ** |
| **atypical resting memory intermediate** | CD40 | -0.69054813675 | 0.0062932158008849 | ** |
| | CD38 | 0.24276572325 | 0.414235965453174 | ns |
| | CD86 | 0.2835030775 | 0.00765449516011318 | ** |
| | CD268 | -0.80858675975 | 0.000129213447734692 | *** |
| | CD24 | -0.1739573515 | 0.136015536814727 | ns |
| | HLA-DR | -0.52783622825 | 0.00765449516011318 | ** |
| | IgD | -0.161580845 | 0.0220696385738743 | * |
| | CD43 | -0.370387694 | 2.87E+08 | *** |
| | CD22 | -0.538475959 | 0.0100290405581466 | * |
| | CD21 | -0.51276106325 | 0.298511085605186 | ns |
| | CD27 | -0.99852959475 | 0.00033210791258819 | *** |
| | CD11c | -0.1091228525 | 0.322393836358045 | ns |
| | CD73 | 0.193429819 | 0.506468590872323 | ns |
| | CD11a | -0.7476758845 | 0.00765449516011318 | ** |
| | CD71 | -0.65669139075 | 8.26E+09 | *** |
| | FCRL5 | -0.02065318775 | 0.824206284672894 | ns |
| | CD45RB | -0.5800513735 | 0.0373827553578212 | * |
| | IgG | -0.34579846325 | 0.0499040202237808 | * |
| | CD95 | -0.96726418775 | 4.28E+08 | *** |
| | CD99 | -118,143,566,925 | 6.06E+09 | *** |
| | CD20 | -0.20316815425 | 0.0262914073097457 | * |
| | CD1d | -0.7367178015 | 2.53E+09 | *** |
| | IgM | -0.59887336225 | 6.10E+09 | *** |

| | | | | |
|---|---|---|---|---|
| DN2 | CD40 | -18,863,167,815 | 5.47E+08 | *** |
| | CD38 | -0.15734937725 | 0.265917203651108 | ns |
| | CD86 | 0.25531893 | 0.00639033632965628 | ** |
| | CD268 | -0.545646661 | 0.0064905713539879 | ** |
| | CD24 | -141,547,385,375 | 3.65E+08 | *** |
| | HLA-DR | -0.08971150525 | 0.991475366491254 | ns |
| | IgD | -0.3660708965 | 1.42E+09 | *** |
| | CD43 | 0.25021457825 | 0.00639033632965628 | ** |
| | CD22 | 147,628,816,575 | 3.65E+08 | *** |
| | CD21 | -173,797,754,025 | 3.65E+08 | *** |
| | CD27 | -10,491,046,245 | 3.01E+08 | *** |
| | CD11c | 296,109,802,375 | 3.65E+08 | *** |
| | CD73 | -0.37482302 | 0.0669656087010243 | ns |
| | CD11a | -0.70252017375 | 0.0499040202237808 | * |
| | CD71 | 0.01589592825 | 0.923987521727834 | ns |
| | FCRL5 | 126,770,739,225 | 6.77E+09 | *** |
| | CD45RB | -124,144,079,575 | 6.01E+08 | *** |
| | IgG | -0.61341523825 | 3.02E+07 | *** |
| | CD95 | 0.82239856325 | 0.0257285806324215 | * |
| | CD99 | -0.01363136725 | 0.821171221927859 | ns |
| | CD20 | 0.745331663 | 6.59E+08 | *** |
| | CD1d | -0.04561026825 | 0.526004948153448 | ns |
| | IgM | 0.661470864 | 5.44E+09 | *** |
| DN3 | CD40 | -166,757,855,075 | 7.30E+09 | *** |
| | CD38 | -0.15779983225 | 0.414235965453174 | ns |
| | CD86 | 0.492703054 | 1.26E+09 | *** |
| | CD268 | -1,790,425,717 | 3.65E+08 | *** |
| | CD24 | -0.3389421435 | 0.0143301891051782 | * |
| | HLA-DR | -102,914,706,475 | 4.28E+08 | *** |
| | IgD | -0.195684183 | 0.00824424497006134 | ** |
| | CD43 | -0.29868019425 | 0.00508533579662169 | ** |
| | CD22 | -148,573,866,975 | 3.65E+08 | *** |
| | CD21 | -1,301,563,936 | 0.000211566217257901 | *** |
| | CD27 | -0.500060548 | 0.0597948231684804 | ns |
| | CD11c | 0.12477759475 | 0.20025473369077 | ns |
| | CD73 | -0.6178017175 | 0.00053143032469898 | *** |
| | CD11a | -160,257,201,375 | 3.65E+08 | *** |
| | CD71 | -0.718346208 | 2.48E+09 | *** |
| | FCRL5 | -0.058575101 | 0.149706553347655 | ns |

| | | | | |
|---|---|---|---|---|
| | CD45RB | -0.49523020225 | 0.035867418685684 | * |
| | IgG | -0.67099750325 | 3.10E+09 | *** |
| | CD95 | -0.02498940925 | 0.995852014370055 | ns |
| | CD99 | -136,710,816,975 | 6.21E+08 | *** |
| | CD20 | -0.1192600145 | 0.375618832821768 | ns |
| | CD1d | -0.75629090875 | 1.61E+09 | *** |
| | IgM | -0.612307013 | 6.10E+09 | *** |
| **resting DN1** | CD40 | 0.635099668 | 0.000125347863267036 | *** |
| | CD38 | 0.6632388845 | 0.0419677698926638 | * |
| | CD86 | -0.2115888655 | 0.0734918097725564 | ns |
| | CD268 | 0.64625126025 | 0.00653506203817657 | ** |
| | CD24 | -0.13665605625 | 0.456831978375915 | ns |
| | HLA-DR | 0.27866984775 | 0.136015536814727 | ns |
| | IgD | 0.00500160725 | 0.672929381565716 | ns |
| | CD43 | -0.30109883925 | 0.00183827477021223 | ** |
| | CD22 | 0.506580153 | 0.00593207277018695 | ** |
| | CD21 | 0.68013525725 | 0.0336960508625353 | * |
| | CD27 | -100,254,074,475 | 0.000237995345732927 | *** |
| | CD11c | -0.17801172275 | 0.00135929750779692 | ** |
| | CD73 | 13,260,731,875 | 0.00653506203817657 | ** |
| | CD11a | 0.399550103 | 0.0695158343556909 | ns |
| | CD71 | 0.12887253775 | 0.59907908281269 | ns |
| | FCRL5 | 0.01868716125 | 0.767841254836371 | ns |
| | CD45RB | -1,135,509,074 | 6.28E+09 | *** |
| | IgG | 0.10770847875 | 0.75416706129664 | ns |
| | CD95 | -0.901228449 | 9.71E+09 | *** |
| | CD99 | 0.0167897545 | 0.891086221650957 | ns |
| | CD20 | -0.16023107125 | 0.0381613101238847 | * |
| | CD1d | -0.05324019225 | 0.969099564462485 | ns |
| | IgM | 0.017885473 | 0.928751050632021 | ns |

**Table S10. Mixed effects model of phenotypic stages across timepoints**

| Stage | Comparison | Adjusted P-value (BH) | Significance |
|---|---|---|---|
| **activated** | 1-2_months_post_vacc_2 - 1_month_post_vacc_1 | 0.999961760514947 | ns |
| | 1_2_months_post_vacc_2 - 1_week_post_vacc_1 | 0.999961760514947 | ns |

| | | | |
|---|---|---|---|
| | 1_2_months_post_vacc_2 - 1_week_post_vacc_2 | 0.999961760514947 | ns |
| | 1_2_months_post_vacc_2 - 3_8_months_post_vacc_2 | 0.999961760514947 | ns |
| | 1_2_months_post_vacc_2 - 8_10_months_post_infection | 0.999961760514947 | ns |
| | 1_month_post_vacc_1 - 1_week_post_vacc_1 | 0.999961760514947 | ns |
| | 1_month_post_vacc_1 - 1_week_post_vacc_2 | 1.16E-228 | *** |
| | 1_month_post_vacc_1 - 3_8_months_post_vacc_2 | 2.77E-89 | *** |
| | 1_month_post_vacc_1 - 8_10_months_post_infection | 0.999961760514947 | ns |
| | 1_week_post_vacc_1 - 1_week_post_vacc_2 | 0.999961760514947 | ns |
| | 1_week_post_vacc_1 - 3_8_months_post_vacc_2 | 0.999961760514947 | ns |
| | 1_week_post_vacc_1 - 8_10_months_post_infection | 0.999961760514947 | ns |
| | 1_week_post_vacc_2 - 3_8_months_post_vacc_2 | 0.000562618147690535 | *** |
| | 1_week_post_vacc_2 - 8_10_months_post_infection | 0.999961760514947 | ns |
| | 3_8_months_post_vacc_2 - 8_10_months_post_infection | 0.999961760514947 | ns |
| **early intermediate stage 1** | 1_2_months_post_vacc_2 - 1_month_post_vacc_1 | 0.999912820166858 | ns |
| | 1_2_months_post_vacc_2 - 1_week_post_vacc_1 | 0.999912820166858 | ns |
| | 1_2_months_post_vacc_2 - 1_week_post_vacc_2 | 0.999912820166858 | ns |
| | 1_2_months_post_vacc_2 - 3_8_months_post_vacc_2 | 0.999912820166858 | ns |
| | 1_2_months_post_vacc_2 - 8_10_months_post_infection | 0.999912820166858 | ns |
| | 1_month_post_vacc_1 - 1_week_post_vacc_1 | 0.999912820166858 | ns |
| | 1_month_post_vacc_1 - 1_week_post_vacc_2 | 0.999912820166858 | ns |
| | 1_month_post_vacc_1 - 3_8_months_post_vacc_2 | 0.999912820166858 | ns |
| | 1_month_post_vacc_1 - 8_10_months_post_infection | 0.999912820166858 | ns |
| | 1_week_post_vacc_1 - 1_week_post_vacc_2 | 0.999912820166858 | ns |
| | 1_week_post_vacc_1 - 3_8_months_post_vacc_2 | 0.999912820166858 | ns |
| | 1_week_post_vacc_1 - 8_10_months_post_infection | 4.00E-106 | *** |

| | | | |
|---|---|---|---|
| | 1_week_post_vacc_2 - 3_8_months_post_vacc_2 | 0.999912820166858 | ns |
| | 1_week_post_vacc_2 - 8_10_months_post_infection | 0.999912820166858 | ns |
| | 3_8_months_post_vacc_2 - 8_10_months_post_infection | 0.999912820166858 | ns |
| **early intermediate stage 2** | 1_2_months_post_vacc_2 - 1_month_post_vacc_1 | 2.40E-103 | *** |
| | 1_2_months_post_vacc_2 - 1_week_post_vacc_1 | 6.44E-252 | *** |
| | 1_2_months_post_vacc_2 - 1_week_post_vacc_2 | 1.65E-135 | *** |
| | 1_2_months_post_vacc_2 - 3_8_months_post_vacc_2 | 3.87E-220 | *** |
| | 1_2_months_post_vacc_2 - 8_10_months_post_infection | 1.11E-232 | *** |
| | 1_month_post_vacc_1 - 1_week_post_vacc_1 | 0 | *** |
| | 1_month_post_vacc_1 - 1_week_post_vacc_2 | 0 | *** |
| | 1_month_post_vacc_1 - 3_8_months_post_vacc_2 | 0 | *** |
| | 1_month_post_vacc_1 - 8_10_months_post_infection | 0 | *** |
| | 1_week_post_vacc_1 - 1_week_post_vacc_2 | 1.52E-35 | *** |
| | 1_week_post_vacc_1 - 3_8_months_post_vacc_2 | 6.54E-35 | *** |
| | 1_week_post_vacc_1 - 8_10_months_post_infection | 7.06E-47 | *** |
| | 1_week_post_vacc_2 - 3_8_months_post_vacc_2 | 3.62E-103 | *** |
| | 1_week_post_vacc_2 - 8_10_months_post_infection | 1.35E-113 | *** |
| | 3_8_months_post_vacc_2 - 8_10_months_post_infection | 0.0202436094489809 | * |
| **late intermediate stage** | 1_2_months_post_vacc_2 - 1_month_post_vacc_1 | 0.999971334917754 | ns |
| | 1_2_months_post_vacc_2 - 1_week_post_vacc_1 | 0.999971334917754 | ns |
| | 1_2_months_post_vacc_2 - 1_week_post_vacc_2 | 0.999971334917754 | ns |
| | 1_2_months_post_vacc_2 - 3_8_months_post_vacc_2 | 6.15E-42 | *** |
| | 1_2_months_post_vacc_2 - 8_10_months_post_infection | 0.500688787610647 | ns |
| | 1_month_post_vacc_1 - 1_week_post_vacc_1 | 0.999971334917754 | ns |
| | 1_month_post_vacc_1 - 1_week_post_vacc_2 | 0.999971334917754 | ns |

| | | | |
|---|---|---|---|
| | 1_month_post_vacc_1 - 3_8_months_post_vacc_2 | 0.999971334917754 | ns |
| | 1_month_post_vacc_1 - 8_10_months_post_infection | 0.999971334917754 | ns |
| | 1_week_post_vacc_1 - 1_week_post_vacc_2 | 0.999971334917754 | ns |
| | 1_week_post_vacc_1 - 3_8_months_post_vacc_2 | 0.999971334917754 | ns |
| | 1_week_post_vacc_1 - 8_10_months_post_infection | 0.999971334917754 | ns |
| | 1_week_post_vacc_2 - 3_8_months_post_vacc_2 | 0.999971334917754 | ns |
| | 1_week_post_vacc_2 - 8_10_months_post_infection | 0.999971334917754 | ns |
| | 3_8_months_post_vacc_2 - 8_10_months_post_infection | 2.03E-33 | *** |
| **resting** | 1_2_months_post_vacc_2 - 1_month_post_vacc_1 | 3.40E-21 | *** |
| | 1_2_months_post_vacc_2 - 1_week_post_vacc_1 | 4.07E+08 | *** |
| | 1_2_months_post_vacc_2 - 1_week_post_vacc_2 | 9.14E-46 | *** |
| | 1_2_months_post_vacc_2 - 3_8_months_post_vacc_2 | 4.25E-106 | *** |
| | 1_2_months_post_vacc_2 - 8_10_months_post_infection | 0 | *** |
| | 1_month_post_vacc_1 - 1_week_post_vacc_1 | 2.41E-18 | *** |
| | 1_month_post_vacc_1 - 1_week_post_vacc_2 | 7.84E+09 | *** |
| | 1_month_post_vacc_1 - 3_8_months_post_vacc_2 | 1.31E-30 | *** |
| | 1_month_post_vacc_1 - 8_10_months_post_infection | 7.77E-182 | *** |
| | 1_week_post_vacc_1 - 1_week_post_vacc_2 | 1.03E-26 | *** |
| | 1_week_post_vacc_1 - 3_8_months_post_vacc_2 | 5.15E-57 | *** |
| | 1_week_post_vacc_1 - 8_10_months_post_infection | 7.28E-146 | *** |
| | 1_week_post_vacc_2 - 3_8_months_post_vacc_2 | 2.66E-11 | *** |
| | 1_week_post_vacc_2 - 8_10_months_post_infection | 3.52E-166 | *** |
| | 3_8_months_post_vacc_2 - 8_10_months_post_infection | 8.55E-51 | *** |

**Table S11. Pairwise Wilcoxon Test of B cell subset composition per phenotypic stage**

| Cluster | stage | Adjusted P-value (BH) | Significance |
|---|---|---|---|
| DN2 | Stage | 0.473245778611632 | ns |
| | early intermediate stage 2 | 0.031385681162922 | * |
| | late intermediate stage | 0.763193161139246 | ns |
| | antigen-unspecific | 0.0107922510701337 | * |
| | activated | 0.101152506030555 | ns |
| | resting | 0.000276363003841672 | *** |
| DN3 | early intermediate stage 1 | 0.347842401500938 | ns |
| | early intermediate stage 2 | 0.592467674246727 | ns |
| | late intermediate stage | 0.347842401500938 | ns |
| | antigen-unspecific | 6.14E+09 | *** |
| | activated | 1 | ns |
| | resting | 0.000292052908529538 | *** |
| CD24+ intermediate | early intermediate stage 1 | 0.852479228088984 | ns |
| | early intermediate stage 2 | 0.118130724866359 | ns |
| | late intermediate stage | 0.109946536133956 | ns |
| | antigen-unspecific | 0.206874498016988 | ns |
| | activated | 0.118130724866359 | ns |
| | resting | 0.000276363003841672 | *** |
| activated | early intermediate stage 1 | 0.303866312228651 | ns |
| | early intermediate stage 2 | 0.447850173397028 | ns |
| | late intermediate stage | 0.0729233506723895 | ns |
| | antigen-unspecific | 0.0656836935321024 | ns |
| | activated | 0.0240100150372998 | * |
| | resting | 0.324492923689181 | ns |
| CD73+ resting memory | early intermediate stage 1 | 0.60400481453113 | ns |
| | early intermediate stage 2 | 0.185333853365412 | ns |
| | late intermediate stage | 0.619936535726009 | ns |
| | antigen-unspecific | 1.89E+06 | *** |
| | activated | 0.0610570084254295 | ns |
| | resting | 0.682126286290872 | ns |
| CD73+ activated | early intermediate stage 1 | 0.625312248726883 | ns |
| | early intermediate stage 2 | 0.965242709409439 | ns |
| | late intermediate stage | 0.625312248726883 | ns |
| | antigen-unspecific | 8.16E+05 | *** |
| | activated | 0.0681854730635218 | ns |
| | resting | 0.0489790112987274 | * |
| resting DN1 | early intermediate stage 1 | 0.604120369704514 | ns |
| | late intermediate stage | 0.0774555709467603 | ns |
| | antigen-unspecific | 1.81E+09 | *** |
| | activated | 0.435563106672319 | ns |

| | | | |
|---|---|---|---|
| | early intermediate stage 2 | 0.458187714566384 | ns |
| | resting | 0.458187714566384 | ns |
| **atypical resting intermediate** | early intermediate stage 1 | 0.473245778611632 | ns |
| | early intermediate stage 2 | 0.371646243880632 | ns |
| | late intermediate stage | 0.706289645314035 | ns |
| | antigen-unspecific | 8.16E+05 | *** |
| | activated | 0.121361565264004 | ns |
| | resting | 0.121361565264004 | ns |
| **CD24 resting memory** | early intermediate stage 1 | 0.642958748221906 | ns |
| | late intermediate stage | 0.933334927142977 | ns |
| | antigen-unspecific | 2.54E+06 | *** |
| | early intermediate stage 2 | 0.624910349960285 | ns |
| | resting | 0.058321479374111 | ns |
| | activated | 0.058321479374111 | ns |
| **activated memory** | early intermediate stage 1 | 0.763193161139246 | ns |
| | early intermediate stage 2 | 0.022290800664281 | * |
| | late intermediate stage | 0.763193161139246 | ns |
| | antigen-unspecific | 1.58E+08 | *** |
| | activated | 0.0347717323327079 | * |
| | resting | 0.763193161139246 | ns |
| **CD73+/CD24+ resting memory** | early intermediate stage 1 | 0.534124083597768 | ns |
| | late intermediate stage | 0.235301090564248 | ns |
| | antigen-unspecific | 1.42E+06 | *** |
| | activated | 0.00617135353977459 | ** |
| | early intermediate stage 2 | 0.702859884284033 | ns |
| | resting | 0.235301090564248 | ns |

**Table S12. Spectral Flow Cytometry phenotypic panel**

| Cell Surface Marker | Laser | Channel | Fluorophore | Clone | Company | REF ID | Dilution |
|---|---|---|---|---|---|---|---|
| CD24 | UV | UV2 | BUV395 | ML5 | BD | 566221/563818 | 1/80 |
| Viability | | UV6 | Live/Dead Blue | - | thermofisher | L34961 | 1/125 |
| HLA-DR | | UV7 | BUV496 | G46-6 | BD | 749866 | 1/80 |
| IgD | | UV9 | BUV563 | IA6-2 | BD | 741394 | 1/160 |
| CD43 | | UV10 | BUV661 | 1G10 | BD | 750301 | 1/160 |
| Tetanus toxoid | | UV11 | BUV615 | NA | NA | NA | NA |
| CD22 | | UV14 | BUV737 | HIB22 | BD | 741830 | 1/20 |
| CD21 | | UV16 | BUV805 | B-ly4 | BD | 742008 | 1/160 |

| | | | | | | | |
|---|---|---|---|---|---|---|---|
| SARS-CoV-2 spike | Violet | V1 | BV421 | NA | NA | NA | NA |
| CD20 | | V3 | eFluor450 | 2H7 | thermofisher | 48-0209-41/48-0209-42 | 1/160 |
| CD27 | | V5 | BV480 | L128 | BD | 566188 | 1/20 |
| CD3 (DUMP) | | V7 | BV510 | UCHT1 | Biolegend | 300447/300448 | 1/80 |
| CD4 (DUMP) | | V7 | BV510 | OKT4 | Biolegend | 317443/317444 | 1/80 |
| CD16 (DUMP) | | V7 | BV510 | 3G8 | Biolegend | 302047/302048 | 1/80 |
| CD14 (DUMP) | | V7 | BV510 | 63D3 | BioLegend | | 1/80 |
| CD56 (DUMP) | | V7 | BV510 | HCD56 | Biolegend | 318339/318340 | 1/80 |
| CD19 | | V8 | BV570 | HIB19 | Biolegend | 302235/302236 | 1/40 |
| CD11c | | V10 | BV605 | B-ly6 | BD | 563930/563929 | 1/160 |
| CD73 | | V11 | BV650 | AD2 | BD | 742633 | 1/80 |
| CD11a | | V13 | BV711 | HI111 | Biolegend | 301237 | 1/20 |
| CD71 | | V14 | BV750 | M-A712 | BD | 747308 | 1/80 |
| FcRL5 | | V15 | BV785 | 509F6 | BD | 749602 | 1/100 |
| SARS-CoV-2 spike | Blue | B1 | BB515 | NA | NA | NA | NA |
| IgM | | B4 | Sparkblue550 | MHM-88 | Biolegend | 314555 | 1/80 |
| CD1d | | B10 | PerCP-eFluor710 | 51.1 | thermofisher | 46-0016-42 | 1/20 |
| CD99 | Yellow/Green | YG9 | PE-Cy7 | 3B2/TA8 | BioLegend | 371314 | 1/20 |
| CD45RB | | YG1 | PE | MEM-55 | Biolegend | 310204 | 1/80 |
| IgG | | YG3 | PE-CF594 | G18-145 | BD | 562538 | 1/160 |
| CD95 | | YG5 | PE-Cy5 | DX2 | Biolegend | 305610 | 1/320 |
| CD268 | Red | R2 | AF647 | 11C1 | BD | 564817 | 1/80 |
| CD86 | | R4 | APC-R700 | 2331 (FUN-1) | BD | 565149 | 1/320 |
| CD40 | | R7 | APC-Fire750 | 5C3 | BioLegend | 334344 | 1/20 |
| CD38 | | R8 | APC-Fire810 | HIT2 | Biolegend | 303549/303550 | 1/160 |

125

#### Supplementary materials and methods

##### Probe labeling methods for SARS-CoV-2-specific B cell selection

We tested several labeling methods to generate labelled SARS-CoV-2 proteins for detection of antigen-specific B cells by flow cytometry and scRNAseq (Supplementary figure S2A): Barcoded streptavidin, in-house produced barcoded streptavidin, direct conjugation of proteins of interest, and dCode Klickmer probes based on a dextramer backbone with up to 20 streptavidin acceptor sites. Direct conjugation of an oligonucleotide barcode to the protein of interest was performed using the Protein-Oligonucleotide Conjugation kit by Vector Laboratories. An amino-oligonucleotide with the correct barcode set-up for 5' sequencing by 10x Genomics was conjugated to HIV-1 Envelope protein GT1.1 according to the manufacturer's protocol. Conjugated protein was separated from unconjugated protein and unbound oligonucleotide by size-exclusion chromatography.

All SARS-CoV-2 proteins (WT S, Delta S, Omicron BA.1 RBD, Omicron BA.2 RBD, Omicron BA4/5 S, H1N1 trimer) were prepared in two different colors to detect double positive SARS-CoV-2-specific B cells and eliminate any fluorochrome-specific B cells. Barcoded streptavidin probes were prepared either by incubating protein with TotalSeqC barcoded streptavidin (BioLegend) (Barcoded streptavidin approach) or streptavidin with a fluorochrome (BioLegend) (In-house barcoded streptavidin, direct protein conjugation approaches) in a 2:1 protein to streptavidin ratio for 1 h in the dark at 4°C. For the in-house produced barcoded streptavidin probe, a biotinylated oligonucleotide barcode was added in either a 1:1 or 2:1 oligonucleotide to streptavidin ratio in addition to protein. For the conjugated protein approach, direct conjugation of an oligonucleotide barcode to the protein of interest was performed using the Protein-Oligonucleotide Conjugation kit by Vector Laboratories. An amino-oligonucleotide with the correct barcode set-up for 5' sequencing by 10x Genomics was conjugated to HIV-1 Envelope protein GT1.1 according to the manufacturer's protocol. Conjugated protein was separated from unconjugated protein and unbound oligonucleotide by size-exclusion chromatography. This conjugated protein was incubated with streptavidin in a 2:1 protein ratio as described above. After protein loading, 10 μM biotin solution was added to block any open streptavidin acceptor sites and avoid aspecific binding of probes to each other causing a false positive result and probes were incubated for 30 min in the dark at 4°C. SARS-CoV-2 dextramer probes were prepared using the (dCode) Klickmer products from Immudex, which contain both single stranded oligonucleotides and fluorophores for detection by flow cytometry and single-cell RNA sequencing. First, proteins were incubated with either PE dCode Klickmer or APC Klickmer (Immudex) in a 5:1 protein to dextramer ratio for 1 h in the dark at 4°C. Next, 10 μM biotin solution was added. Probes were incubated with biotin for 30 min in the dark at 4°C. After this step, the different colour probes were mixed and used for cell staining.

All labeling methods were tested by flow cytometry bead assay or PBMC staining for their binding abilities (Figure S2B-E). For the bead assay, anti-mouse Ig/negative control CompBeads (BD Biosciences) were stained with 0.125 μg mouse anti-human Ig primary antibody, followed by 0.5 μg of the antibody to test. For PBMC staining, isolated PBMCs from healthy donor buffy coats were thawed briefly in the water bath at 37°C and placed in 40 mL RPMI supplemented with 20% FCS. Next, SARS-CoV-2 probes were added to either the beads or PBMCs and incubated for 30 min. Beads or cells were washed with FACS buffer. Next, cells were stained with a B cell phenotype antibody panel for FACS detection of B cell populations (CD4 eF780, CD3 eF780, CD14 eF780, CD16 eF780, viability dye eF780, CD19 AF700, IgD BV785, IgG BV605, CD27 BV421) for 30 min at 4°C in the dark. After incubation, cells were washed with FACS buffer. Cells and beads were analyzed using flow cytometry analysis on the BD FACS Aria IIu SORP 4 laser sorter or BD Fortessa machine.
